# Rational Design of Unsaturated, Thioether Ionizable Lipids for Enhanced *In vivo* mRNA Delivery

**DOI:** 10.1101/2024.12.30.630759

**Authors:** Eleni Samaridou, Johanna Simon, Moritz Beck-Broichsitter, Gary Davidson, Pavel A. Levkin

## Abstract

Therapies based on mRNA technology have offered hope to millions of patients worldwide by disrupting the way we treat diseases. The safe and functional delivery of the delicate mRNA molecules to the target tissue is a crucial step in the development of effective vaccines and therapeutic interventions. Lipid nanoparticles (LNP) are the most clinically advanced delivery vehicles for mRNA drugs. Key to the success of LNP is the inclusion of an ionizable cationic lipid. However, the structure-function relationships between ionizable lipids and efficient *in-vivo* mRNA delivery are still poorly understood. In this work, we focused on the rational design and sequential structural optimization of previously identified ionizable lipids that performed well *in vitro*, but not *in vivo*. Through two distinct iterative optimization cycles — one targeting the lipid tail and the other the headgroup — we aimed to understand how the fusogenicity and apparent p*K*a of the ionizable lipid contribute to LNP delivery performance *in vivo*. By engineering unsaturated lipids with more hydrophobic, less protonatable amino headgroups with longer alkyl group at the tertiary nitrogen, we achieved both significant improvement of protein expression *in vitro*, reduced hemolysis risk, and more than 200-fold improvement of *in vivo* mRNA delivery. When compared head-to-head to a market-approved LNP benchmark, the newly developed ionizable lipids/LNP resulted in equally highly efficient *in vivo* mRNA delivery, with strong liver and spleen tropism upon intravenous injection, while matching the safety of the approved platform. Our findings are pivotal for the development of next-generation mRNA-LNP therapies and vaccines.

## Introduction

Lipid nanoparticles (LNP) have revolutionized the development of nucleic acid-based preventive and therapeutic applications^1-4^. Apart from the established role of these delivery vehicles in the success of mRNA vaccines against infectious diseases (i.e., SARS-CoV-2 and respiratory syncytial virus), more than 50 different mRNA-LNP drug products are currently in clinical development, with particular focus on infectious diseases, oncology and genetic disorders^5^.

LNP are typically composed of four lipidic ingredients: an ionizable cationic lipid, a phospholipid (also referred to as helper lipid), cholesterol and a polymer-lipid conjugate. Each component has a specific role in delivering RNA drugs, with the ionizable cationic lipid being the main performance driver of the LNP as delivery vehicle^6^. Owing to the presence of the ionizable lipid, the charge state of the LNP varies as a function of the environmental pH. An apparent p*K*_a_ value of the formulation (defined as the pH value at which the LNP becomes protonated) between 6.0 and 7.0 has been described to be ideal for RNA delivery, since the formulation presents no-to-low charge at physiological pH (reducing the risk for toxicity), while in the acidic endosomes, it acquires a higher overall positive charge, leading to the endosomal membrane disruption and release of the RNA into the cytoplasm.^7^

Given the importance of the ionizable cationic lipid component for the success of the LNP, extensive scientific efforts over the last decades have been dedicated to the structural optimization of these lipidic excipients, aiming to further improve the LNP specificity and efficacy in cargo delivery, as well as immunogenicity and shelf-life, while keeping the manufacturing costs low. The quest for the optimal design of ionizable lipids began in the early 2000s when Semple *et al*.^8^ demonstrated the potential of an ionizable aminolipid, DODAP (1,2-dioleoyl-3-dimethylammonium propane), for improving the pharmacokinetics of LNP formulations compared to delivery systems containing permanently-charged aminolipids or cationic liposome-based complexes. A few years later, Heyes *et al*. showed that increasing the degree of unsaturation in the hydrophobic domain of ionizable lipids enhanced their fusogenicity, thereby supporting endosomal escape and increasing the potency of LNP formulations^9^. This pioneering work led to the development of DLinDMA (1,2-dilinoleyloxy-N,N-dimethyl-3-aminopropane), the first ionizable lipid to enter human clinical trials^10^, subsequently leading to the “birth” of DLin-MC3-DMA (dilinoleyl-methyl-4-dimethylaminobutyrate) commonly known as “MC3”, the ionizable lipid used in the first marketed RNA-LNP drug (Onpattro™). Since then, several groups have tried to understand the contribution of different structural features of the ionizable lipids to their potency, biodegradability and/or their ability to specifically target tissues of interest^11-17^. The next generation of ionizable lipid design has focused on the number of tertiary amines, the number and branching of the alkyl tails, and the presence of biodegradable motifs^16^, leading to the development of the ionizable lipids used in the two marketed Covid-19 mRNA vaccines, namely ALC-315 (used in Comirnaty^®^) and SM-102 (used in SpikeVax^®^)^18,19^.

Current research relies on the use of combinatorial chemistry, high-throughput synthesis and screening approaches with the aim of identifying ionizable lipids that allow for site-specific RNA delivery, while minimizing off-target effects, in a rapid and cost-effective manner^20-23^. Such approaches may enable the rapid identification of well-performing lipid structures *de novo in vitro*, but potential pitfalls are shifted to later stages of development, such as a lack of scalability of lipid synthesis and LNP formulation at a reasonable purity and price as well as *in vivo* performance.

Concurrently, artificial intelligence (AI) and machine learning approaches are being used to process complex data sets and generate models that could enable intuitive lipid design^11,24,25^. Nevertheless, these approaches are still under development and, therefore, several challenges need to be addressed before their full potential can be realized. Examples are the limited access to large quantity of high-quality data to properly train and create a reliable and accurate AI model, the inability of these approaches to address several outputs simultaneously (such as toxicity concerns, apart from efficacy and biodistribution) and the issues of organization and interpretation of the diverse available data in the field and literature^26^.

In this work, we instead focused on the rational design of the ionizable lipid structure, building on advanced lipid chemistry and customizing it for RNA delivery, through specific and targeted structural optimizations. This approach resulted in a drastic increase in the efficacy of mRNA delivery both *in vitro* and *in vivo*. Previously, we developed a facile two-step chemical synthesis of cationic thioether lipids based on the thiol-yne photoclick chemistry^27^ and demonstrated the general applicability of these lipids for *in vitro* cell transfection^28,29^. To further explore the potential of these lipids for use in LNP for *in vivo* mRNA delivery, we focused on the role of fusogenicity and apparent *pK*_a_ in cellular uptake, endosomal escape and, importantly, the hemolytic activity of the formulation. For this, we sequentially modified the lipid structures by first including longer, unsaturated tail moieties (hydrophobic alkyl groups), followed by modifying the ionizable head groups. The new lipids, resulting from our sequential optimizations, were incorporated into LNP and three of these novel lipids showed highly efficient *in vitro* and *in vivo* mRNA delivery, improved p*K*_a_ values and, consequently, low hemolytic activity. These results pave the way for the effective design of LNP-based therapeutic tools and provide valuable insights into the critical structural components of lipid moieties in the development of novel vaccines and mRNA therapeutics.

## Results and discussion

### Thioether lipids for LNP-mediated mRNA delivery

In a previous work, we presented a facile modular and scalable approach employing thiol-yne “click” chemistry and amide coupling for the high-throughput combinatorial synthesis of a novel library of cationic thioether lipids with two saturated hydrophobic tails of variable lengths and a linker group structurally mimicking the glycerol core of phospholipids (Figure 1A)^28^. From this library, we were able to identify several top-performing lipids in terms of *in vitro* delivery of distinct nucleic acid payloads (such as siRNA and plasmid DNA) in “difficult-to-transfect” cell lines. Specifically, with lipid A1C11 (Figure 1A), as transfection reagent (lipoplex formulation), a strong siRNA silencing and a high plasmid DNA transfection rate *in vitro* were induced.

**Figure 1:**
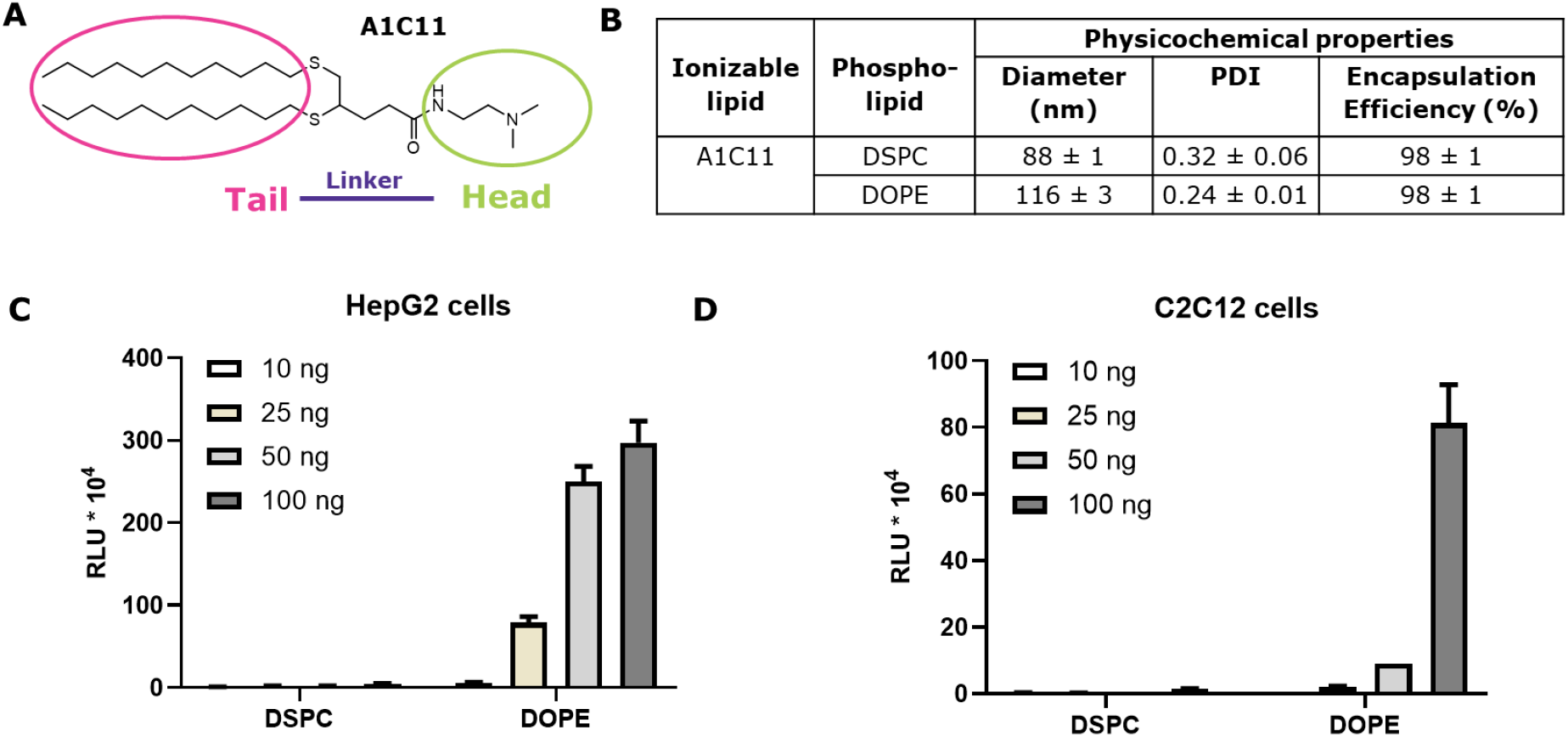
A) Structure of the thioether lipid A1C11. B) Physicochemical properties of mRNA-loaded LNP formulations containing A1C11 (with varying helper lipids DSPC and DOPE). In vitro luciferase activity in HepG2 (C) and C2C12 cells (D) after incubation with LNP formulations (at four different mRNA doses). Mean values from triplicates ± standard deviations are shown.

As a first step, we tested lipid A1C11 for its applicability in LNP-mediated mRNA delivery (Figure 1). To this end, LNP were formulated using the same LNP composition as the one found in the marketed Onpattro™ LNP formulation^30,31^. Since the market approval of Onpattro™, its specific LNP composition has been frequently applied in multiple applications, including mRNA delivery for therapeutics and vaccines^16,32,33^. MC3, the ionizable lipid found in Onpattro™, was used as the benchmark for the performance evaluation of the novel ionizable lipid structures tested here.

LNP formulations were then prepared with a reporter mRNA (encoding firefly luciferase) using the thioether lipid A1C11 (Figure 1A) and varying the phospholipid type (i.e., DSPC versus DOPE). In line with what has been reported in the literature,^33-36^ LNP containing DOPE exhibited a larger particle size distribution compared to those containing DSPC (Figure 1B). Both LNP based on A1C11 exhibited high encapsulation efficiency of the mRNA (∼98%), while the type of phospholipid had no effect on the apparent p*K*_a_ of the formulation, which was in both cases >7, indicating that the formulation was positively-charged at physiological pH. Interestingly, when the LNP were tested *in vitro* using a hepatocyte representative cell line (HepG2, as indication for the preference of the LNP to transfect hepatocyte cells, relevant for therapeutic applications) and muscle representative cell line (C2C12, as indication for the preference of the LNP to transfect muscle cells, relevant for vaccine applications) (Figure 1C, D), the DOPE-containing LNP showed nearly 10-fold higher luciferase activity compared to their DSPC counterparts, while both formulations showed overall good cell viability at the four doses tested (SI Figures 2 and 3).

We suspected that a potential reason for this result was associated with the specific contribution of the chosen helper lipid on the membrane fluidity of LNP, which is required for cellular uptake and, more importantly, for endosomal escape. As previously reported, the structural differences of DSPC and DOPE results in differential packing within the LNP, with DOPE adopting a less stable hexagonal phase, while DSPC adopts a more stable lamellar phase^16,36-38^. The cone-like geometry of DOPE enables a higher membrane fluidity and fusogenicity, leading to enhanced endosomal release of the RNA cargo^4,36^. However, DOPE-containing LNP have been associated with tolerability issues^39^ when tested *in vivo* and, also, with manufacturing and stability challenges (due to the risk of particle fusion upon handling and storage)^36^.

### Optimizing the ionizable lipid tail chemistry to increase LNP fusogenicity

For this reason, we focused on increasing the fusogenicity of the ionizable cationic lipid itself, rather than relying on the effects of helper lipids. Heyes *et al*. reported that decreasing the degree of saturation in the alkyl chains of the ionizable lipid, by introducing 1, 2 or 3 double bonds per alkyl chain, led to a significant increase in the formulation’s fusogenicity and, consequently, its biological performance^9^. With this insight, we designed five novel structures using A1C11 as the starting point (Figure 2 and Table S1). In our initial iteration, we evaluated the impact of introducing double bonds into the alkyl tails of the A1C11 lipid. This resulted in the lipid A1C11_D5, where a single set of double bonds was introduced at the C5 position of each C11 tail. The double bond position was carefully selected to be in the middle of each alkyl chain, as this has been described to have the strongest influence on the lipid curvature, leading to a decrease in phase transition temperature and an increase in fluidity^40,41^.

**Figure 2:**
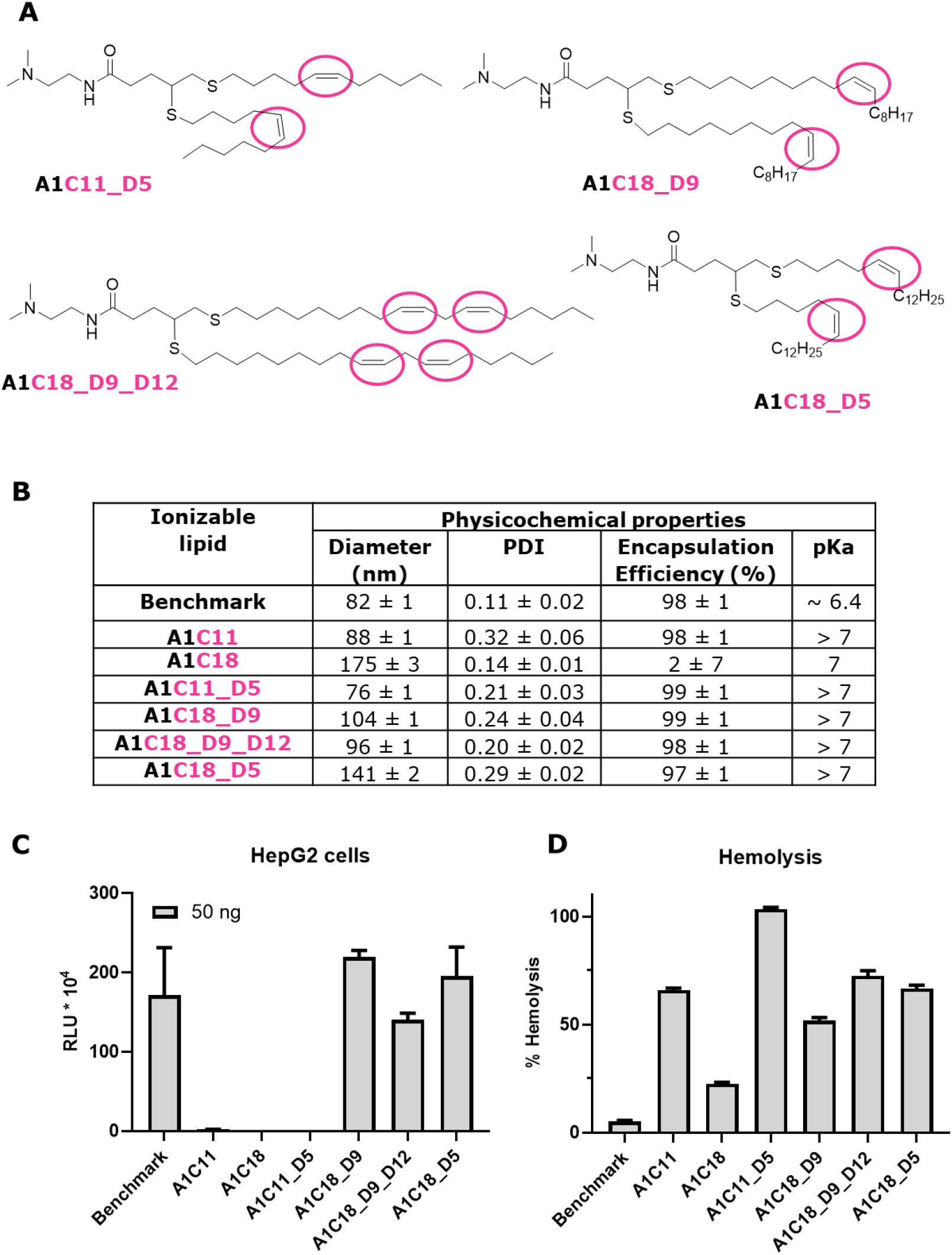
A) Structures of tail-group optimized thioether lipids. B) Physicochemical properties of mRNA-loaded LNP formulations containing novel tail-optimized cationic thioether lipids and the benchmark lipid (MC3). C) In vitro luciferase activity in HepG2 cells after incubation with LNP formulations. D) Hemolytic effect of LNP formulations containing novel tail-optimized cationic thioether lipids and the benchmark lipid (MC3). Mean values from triplicates ± standard deviations are shown. All LNP tested at the same composition (only the ionizable lipid was varied) and at the same mRNA dose levels.

In addition, we hypothesized that the two C11 alkyl chains may be too short to enable high structural curvature and synthesized four additional lipids with two C18 chains, to mimic the structure of the MC3 benchmark (Figure 2A and SI Table S1). At the same time, we introduced one or two sets of double bonds per chain. The position of the double bonds in the new 2xC18 lipids were placed at the middle (C9) of the C18 chains for lipid A1C18_D9, while for lipid A1C18_D9_D12, the double bonds were positioned at both C9 and C12 on each tail. A1C18 served as the saturated 2xC18 control to distinguish between the effect of the alkyl chain length and their saturation. Additionally, we introduced a control lipid, A1C18_D5, with one set of double bonds at the C5 position of each C18 chain, allowing for a direct comparison with lipid A1C11_D5, where only the alkyl chain length differed.

For functional testing, mRNA encoding luciferase was again encapsulated into LNP formed using these ionizable cationic lipids (Figure 2B), together with DSPC as the helper lipid. Luciferase activity and cell viability tests were performed in HepG2 (Figure 2C) and C2C12 cells (SI Figures 3, 4). For comparison, LNP were also formulated with DOPE as the helper lipid (SI Figure 1-4) and characterized as described above.

All LNP displayed comparable physicochemical properties (particle size <140 nm, PDI <0.3, and encapsulation efficiency >90%), except for the saturated A1C18-based LNP that showed only 2% encapsulation efficiency (Figure 2B). This was expected due to the more rigid bilayer structure of the saturated lipid A1C18^9^.

Notably, significantly enhanced mRNA delivery was observed for the longer C18 tails harboring one (A1C18_D5 and A1C18_D9) or two (A1C18_D9_D12) sets of double bonds (Figure 2C, SI Figure 3). Neither inclusion of double bonds alone to the shorter C11 chains or lengthening the chains from C11 to C18 alone showed this effect (SI Figures 1, 3). Interestingly, the helper lipid DOPE did not further enhance the fusogenicity of LNP composed of unsaturated lipids, whereas the DSPC containing LNP showed efficient mRNA delivery, with expression levels similar to the ones of market-approved LNP benchmark (HepG2: Figure 2C and SI Figure 1; C2C12: SI Figure 3).

Nevertheless, the p*K*_a_ remained >7 for all tested formulations (Figure 2B), in contrast to the observations made by Heyes *et al*., where the p*K*_a_ correlated with the degree of saturation^9^. Considering the optimum p*K*_a_ range is 6-7 for delivery^14,42,43^, this raised concerns for potential toxicity. This led us to perform *in vitro* hemolysis assays (Figure 2D)^44,45^ since it has been reported that the hemolytic activity is a linear function of their surface charge / zeta-potential^46^. According to the assay, membrane lysis observed at pH 7.4 is an indication of toxicity, whereas lysis observed at pH 5.5 is a good model for the ability of the LNP to escape vesicular structures upon protonation^45^. As expected, the LNP containing the novel unsaturated lipids exhibited similar membrane fusion levels to the benchmark formulation (SI Figure 5), however, their hemolysis potential was found to be significantly higher compared to the benchmark formulation, likely due to their high p*K*_a_ values and overall charge of LNP at physiological pH (Figure 2D and SI Figure 5).

### Fine tuning the lipid head group to decrease the p*K*_a_ of the LNP

Following these results, our next iterative step was to optimize the p*K*_a_ of the LNP to the desired value between 6 and 7. Initially, we focused on the lipid A1C18_D5, modifying the head group of the ionizable lipid as indicated in Figure 3A, generating the lipids A2C18_D5, A3C18_D5, and A4C18_D5. The piperazine derivatives A3C18_D5 and A4C18_D5 containing an additional tertiary amine did not show reduction of p*K*_a_ and led to higher hemolysis. Surprisingly, the introduction of a bulkier, hydrophobic pentyl group (A2) on the head amine group, while retaining just one tertiary amine, significantly reduced the LNP’s p*K*_a_ (Figure 3B). Good particle size distribution and mRNA encapsulation efficiency were maintained. Interestingly, although A4C18_D5 contained the same pentylated amine, it did not improve the p*K*_a_ value and resulted in strong hemolysis. With the optimized tails identified above, we then additionally synthesized the lipids A2C18_D9 and A2C18_D9_D12 and formulated them as described above using DSPC as helper lipid. Remarkably, when tested *in vitro* in HepG2 (Figure 3D, and SI Figures 6 and 10) and C2C12 cells (SI Figures 8 and 12), the LNP’s based on lipids A2C18_D5, A2C18_D9 and A2C18_D9_D12 displayed greatly improved delivery efficacy, well above the levels seen for the marketed benchmark composition. Importantly, testing in the same cells showed no significant decrease in cell viability for the novel lipids (SI Figures 7, 9, 11 and 13).

**Figure 3:**
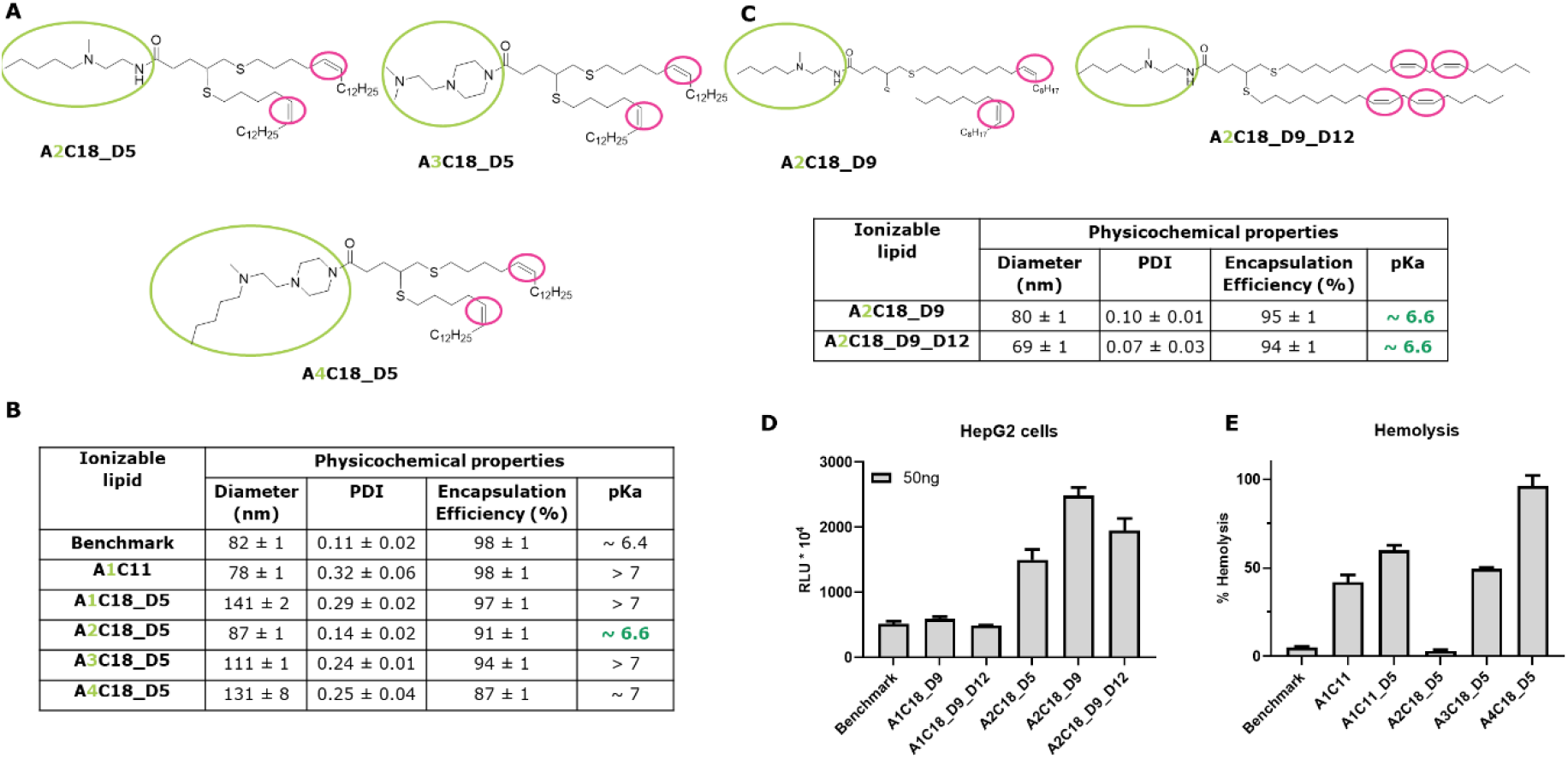
A) and C) Structures of head-group optimized thioether lipids. B) Physicochemical properties of mRNA-loaded LNP formulations containing novel head optimized thioether lipids and the benchmark lipid (MC3). D) In vitro luciferase activity in HepG2 cells incubated with LNP formulations. E) Hemolytic effect of LNP formulations. Mean values from triplicates ± standard deviations are shown.

In agreement with the reduced p*K*_a_ values, LNP containing the novel ionizable lipids A2C18_D5, A2C18_D9 and A2C18_D9_D12 showed greatly reduced hemolysis (Figure 3E), while maintaining high levels of fusogenicity (SI Figures 14 and 15). We suspect this may be attributed to the bulkier hydrophobic head group sterically hindering the ability of the tertiary amine to interact with the plasma membrane of blood cells.

### Altering the *in vivo* biodistribution with novel designed LNP formulations

Having optimized the *in vitro* performance of the novel lipids, we next evaluated the effectiveness of the optimized LNP *in vivo*. We injected luciferase mRNA-loaded LNP formulations (0.25 mg/kg), based on the novel lead lipids A2C18_D5, A2C18_D9 and A2C18_D9_D12, intravenously to Balb/c mice and studied the distribution of bioluminescence *in vivo* (whole body bioluminescence) and *ex vivo* at 6 hours post injection in selected tissues (liver, spleen, kidneys, lungs). As controls, LNP formulations based on the MC3 (benchmark) and our starting point, LNP based on the A1C11 lipid, were used.

As shown in Figure 4, LNP containing the lipids A2C18_D5, A2C18_D9, and A2C18_D9_D12 demonstrated excellent *in vivo* performance, matching the benchmark formulation. Indeed, over 200-fold enhancement in protein expression could be demonstrated for these LNP when compared to the A1C11-based formulation (Figure 4C). Notably, the *ex vivo* analysis of bioluminescence distribution six hours after intravenous injection revealed a distinct tissue tropism of the novel LNP toward the liver. LNP-mediated RNA delivery to the liver offers great therapeutic value for protein replacement therapies and for treating hepatic diseases (such as but not limited to liver fibrosis, nonalcoholic fatty liver disease, and drug-induced liver injury). However, there are several disease indications that would benefit from extra-hepatic LNP distribution (such as cancer, neurological, cardiovascular and vaccine applications). To this point, the A2C18_D9-based LNP exhibited significant localization in the spleen, resulting in a higher spleen-to-liver ratio compared to the marketed LNP benchmark (Figure 4D),. Additionally, the tolerability of the novel LNP was evaluated *in vivo* in terms of unwanted immune responses to the LNP components, which may lead to increased production of different secretory proinflammatory molecules, such as Interleukin 6 (IL-6) or monocyte chemoattractant protein 1 (MCP-1). The induction of IL-6 and MCP-1 cytokines upon *in vivo* injection was measured by ELISA-based assay in the animal plasma collected at different time points post injection (6 and 24 hours post injection). Interestingly, all novel LNP formulations displayed plasma cytokine expression levels (MCP-1 and IL-6) comparable to those observed in the marketed LNP benchmark and the untreated animals, with all cytokine values returning to base levels at 24 hours post injection (Figure 4F).

**Figure 4:**
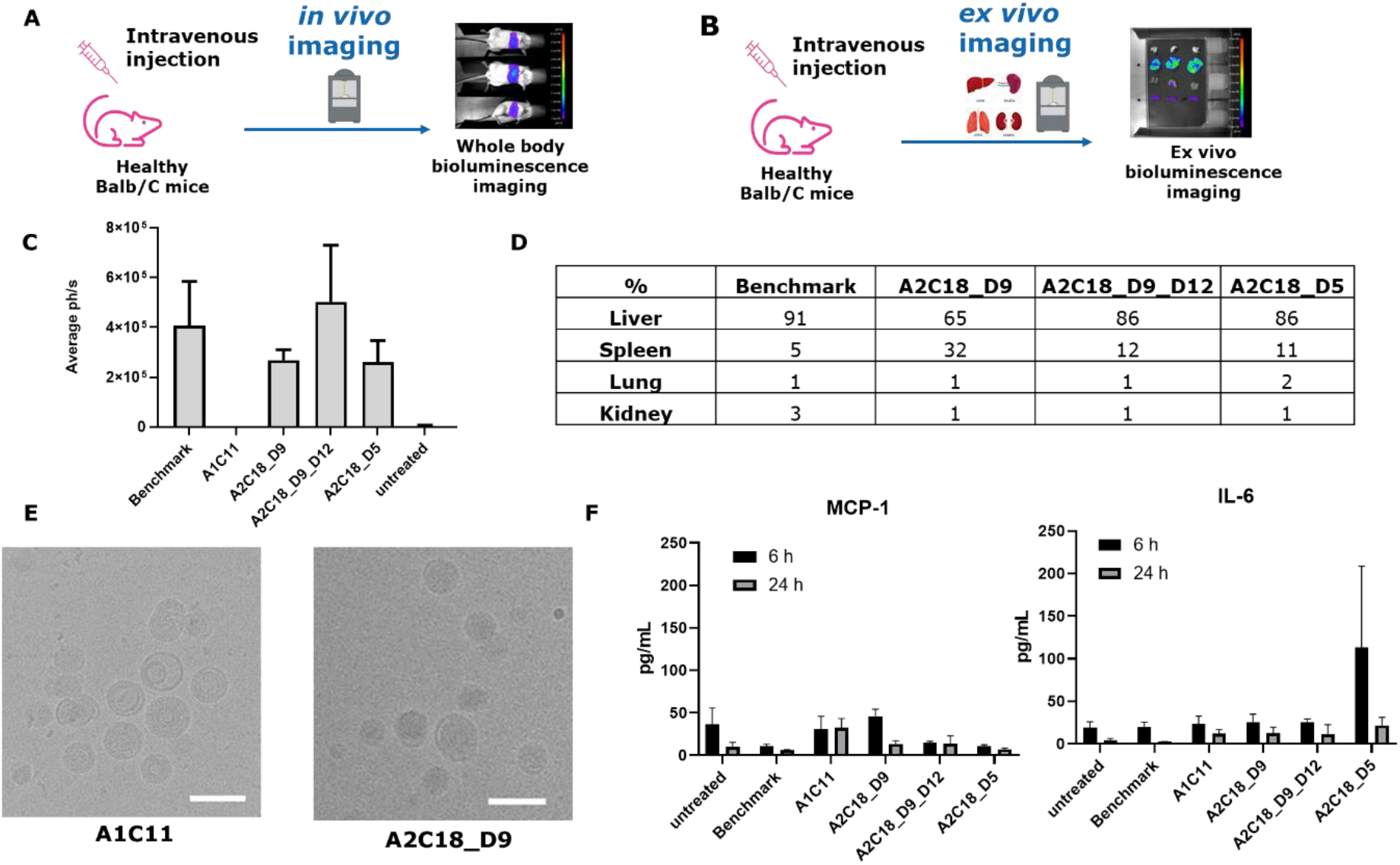
A) In vivo imaging study design. B) Ex vivo imaging study design. C) Whole body in vivo bioluminescence signal 6 h post dosing. D) Ex vivo bioluminescence signal 6 h post dosing. E) CryoTEM images of LNP containing A1C11 or A2C18_D9 as ionizable lipid. Scale bar shown in white = 100 nm. F) Cytokine levels 6 h and 24 h post dosing. Mean values from triplicates ± standard deviations are shown.

### Understanding LNP morphological differences

To evaluate further the potential effect of the different ionizable lipid structures on LNP morphology, electron micrographs of mRNA–LNP, based either on our starting point lipid, A1C11, or the novel lead lipid A2C18_D9 (chosen based on the higher spleen tissue localization), were evaluated. Interestingly, whereas both mRNA-LNP were confirmed to be spherical and at the expected size range (in accordance with the average size and distribution as determined by DLS measurements), significant structural differences were visualized for LNP containing A1C11 vs. A2C18_D9 as ionizable lipid (Figure 4E). LNP containing A1C11 demonstrated bilayer features (like what is observed in liposomal formulations) and a heterogeneous morphology, including multi-lamellar structures. In contrast, LNP containing A2C18_D9 exhibited an electron-dense core, as expected for the typical LNP morphology and, as reported previously by Leung et al.^47^, with a very homogeneous morphology. The morphological differences observed are believed to be directly connected to the structural differences of the ionizable lipids used in each case, owing to their different ratio of the cross-sectional area of their head group to their overall length (e.g., C11 versus unsaturated C18 alkyl chains) resulting in different membrane curvatures^48^. Therefore, these striking morphological differences in the resulting LNP are expected to be related to the different performance observed in the *in vivo* luciferase activity levels between the two tested LNP formulations (Figure 4C).

## Conclusion

In this work, we successfully designed and tested novel ionizable thioether lipids for mRNA delivery, by engineering synthetic unsaturated analogues with more hydrophobic and less exposed protonatable amino head groups. Our novel lipids demonstrated improved physicochemical as well as biological performance, which was particularly evident with respect to *in vitro* and *in vivo* mRNA delivery, showing equal performance compared to the current market approved benchmark LNP formulation. The introduction of unsaturated hydrophobic tails and modified ionizable head groups resulted in optimal physicochemical properties, including reduced p*K*_a_ values and decreased hemolytic activity. The strategic modifications led to effective mRNA delivery with selective organ accumulation, particularly in the liver and spleen. These results indicate that the newly developed lipid structures facilitate efficient cellular uptake and protect mRNA from degradation, addressing the critical issue of translating *in vitro* efficiency to *in vivo* performance. Moreover, the novel LNP formulations exhibited low toxicity and reduced hemolytic membrane disruption *in vitro*.

Overall, the advancements presented in this study offer significant potential for the development of next-generation mRNA vaccines and therapeutic interventions. The newly developed lipid structures provide a robust platform for efficient, targeted, and safe delivery of mRNA, paving the way for improved treatment options in various biomedical applications. These findings also contribute valuable insights into the structure-function relationships in lipid design, which are essential for the continued advancement of LNP-based delivery systems.

## Acknowledgements

PAL and GD thank A. Levkina and M. Mannuss for the support with the project as well as Dr. Khartulyari (Genosynth GmbH) for the help with the synthesis.

## Materials & Methods

### Materials

1,2-Dioleoyl-sn-glycero-3-phosphoethanolamine (DOPE), 1,2-distearoyl-sn-glycero-3-phosphocholine (DSPC), 1,2-dimyristoyl-rac-glycero-3-methoxypolyethylene glycol-2000 (DMG-PEG2000), cholesterol, phosphate-buffered saline (PBS) (1x dPBS) and 6-(*p*-Toluidino)-2-naphthalenesulfonic acid (TNS) were purchased from Sigma-Aldrich (Germany). mRNA encoding for firefly luciferase (CleanCap^®^ FLuc mRNA (5moU)) was acquired from TriLink BioTechnologies (US). Pur-A-Lyzer™ Maxi Dialysis Kit (MWCO 12-14 kDa), Amicon Ultra (MWCO 30 kDa) centrifugal units and Millex^®^-GV filter unit syringe filters (0.22 μm, 13 mm, PVDF) were purchased from MilliporeSigma (Germany). RiboGreen was obtained from ThermoFischer (Germany). Hept-1-yne, potassium thioacetate (KSAc), ammonia (7M), mesyl chloride, N,N-dimethylethane-1,2-diamine, N,N-dimethylaminoethylamine, 1-octadecanethiol, BOC-protected amines (various), pentylchloride, 5% Pd/CaCO_3_, Pd/BaSO4, ethanol, methanol, petroleum ether, dry methanol were purchased from Fisher Scientific (Schwerte, Germany). Dodecyl iodide was purchased from Sigma (Schnelldorf, Germany). N-methyl-1,2-ethylenediamine and trifluoroacetic acid (TFA), dichloromethane (DCM), anhydrous dimethylformamide, diethyl ether, acetic acid, sodium sulfate (Na2SO4), celite (for filtration), molecular sieves (for drying solvents) were bought from Carl Roth (Karlsruhe, Germany). Lindlar catalyst (5% Pd/CaCO_3_ poisoned with Pd or Pd/BaSO4) and sodium methylate were obtained from TCI (Eschborn, Germany). Quinoline was received from Merck KGaA (Darmstadt, Germany). Hydrogen gas was purchased from Air Liquide (Ludwigshafen, Germany). 2-(4-Iodobutoxy)tetrahydro-2H-pyran (1) and dibromide (25) were obtained from GenoSynth GmbH (Berlin, Germany). Oleyl alcohol, linoleyl alcohol ((9Z,12Z)-octadecadien-1-ol), N^1^-BOC, N^2^-methylethane-1,2-diamine, N-methyl-N-pentylamine, BOC-protected compound <u>30</u>, Hexynol 33, dodecyl iodide, n-butyllithium, triisopropylsilylchloride were purchased from Sigma (Germany). All other chemicals and solvents were commercially available and used without further purification.

### Synthesis and characterization of lipids

See Supplementary Information

### LNP formulation and characterization

LNP with a lipid composition of 50:38.5:10:1.5 mol% (ionizable lipid:cholesterol:phospholipid:DMG-PEG_2000_) were prepared by mixing an ethanolic lipid solution with an mRNA aqueous solution in 50 mM citrate buffer at pH 4 at a mRNA concentration of 0.15 mg/mL, using a commercial microfluidic-based mixing device (mRNA:lipid phase flow rate ratio = 3:1, total flow rate = 12 mL/min), to result in mRNA formulated in LNP at a nitrogen-to-phosphate (N/P) ratio of 6. The formulations were dialyzed overnight (at 2-8 ^°^C) against 100-fold volume of 1x dPBS or the appropriate storage matrix buffer. After dialysis, the formulations were up-concentrated to the desired concentration, introduced to the appropriate storage matrix by dilution and finally sterile filtered.

Particle size and polydispersity were determined using a DynaPro™ Plate Reader III (Wyatt Technology, USA). The mRNA concentration and encapsulation efficiency of the final formulations were determined using the RiboGreen assay and a mRNA standard curve and comparing fluorescence in the presence and absence of Triton X-100, using a Tecan Infinite M200 Pro Multi Mode Microplate Reader (Tecan, USA).

### Evaluation of in-situ p*K*_a_ by TNS assay

For the TNS assay, 20 mM citrate/phosphate buffer series (in 150 mM NaCl), covering a pH range between 4.0 – 8.0 (with increments of 0.2), were prepared. In the wells of a 96-well plate, 10 μL of the LNP solution (0.2 mM total lipid), 88 μL of each of the buffers above and 2 μL of the TNS solution (0.3 mM stock solution in water) were mixed. The fluorescence of the TNS (ex: 322 nm, em: 431 nm) was measured using a microplate reader. Fluorescence for each formulation at the various pH values was then normalized to the value at pH 4.0. By assuming that minimum fluorescence represents zero charge, and maximum fluorescence represents 100% charge, p*K*_*a*_ was estimated by measuring the pH at the point exactly halfway between the values of minimum and maximum charge using Prism10.2.1 (GraphPad, USA).

### Hemolysis and membrane fusion assay

Human blood (healthy donors from blood donation bank at Merck KGaA, Darmstadt, Germany) was centrifuged for 5 min at 500 × g. The plasma was aspirated, the isolated human red blood cells were washed twice with 1x dPBS and diluted in either 1x dPBS or citrate buffer saline at pH 5.5 (CBS, 20 mM citrate buffer, 130 mM NaCl) to a 4% vol/vol red blood cell suspension. In a 96-well plate, 40 μL of LNP formulated at a mRNA concentration (0.01 mg/mL) were added to 40 μL of the 4% vol/vol red blood cell suspension in either 1x dPBS or CBS and heated to 37 °C for 1 h. After cooling, the plate was centrifuged at at 500 × g and 4 °C for 5 min; the supernatant was transferred into another 96-well assay plate and the absorption was read at 540 nm. Positive and negative controls were carried out with 0.2% Triton-X (100%) and 1x dPBS alone, respectively.

### *In vitro* transfection

*In vitro* transfection was performed using HepG2 and C2C12 cells (ATCC, USA). HepG2 cells were maintained at 37 °C in a 5% (vol/vol) CO_2_ atmosphere in Eagle’s Minimium Essential Medium) with 10% (vol/vol) fetal bovine serum (FBS), 1x non-essential amino acid (ThermoFisher), 200 mM glutamine and 1x PenStrep (all from Sigma-Aldrich, Germany). C2C12 cells were maintained at 37 °C in a 5% (vol/vol) CO_2_ atmosphere in Dulbecco’s Modified Eagle Medium (high glucose) with 10% (vol/vol) FBS and 1x PenStrep) for a maximum of 20 passages. Cells were passaged 2-3 times per week. Before transfection 10.000 HepG2 or 5.000 C2C12 cells were seeded in white 96-well plates and allowed to attach overnight. The next day, the media was replaced with fresh media containing LNP at a dose of 10, 25, 50 and 100 ng (mRNA dose). Relative firefly luciferase activity and cell viability were assessed ∼24 h after LNP addition using ONE-Glo™ + Tox Luciferase Reporter and Cell Viability Assay (Promega, Germany).

### *In vivo* biodistribution and tolerability

Animal studies were carried out in collaboration with the Innovation Campus Berlin (Nuvisan ICB GmbH) and in strict accordance with AAALAC guidelines and the European Directive 2010/63/EU as well as the German Animal Welfare Act (Tierschutzgesetz). Animal experiments were approved by the Institutional Animal Care and Use Committee and competent regional Animal Care and Use Committees according to §15 TierSchG (application number E0155/23; Berlin, Germany).

Eight-week-old female BALB/c mice were purchased from Janvier Labs and were housed in a Specific and Opportunistic Pathogen Free animal facility. Mice were allowed to acclimatize for at least 1 week. Mice had free access to food and water and were exposed to 12-h light/dark cycles.

Groups of six mice each received one intravenous injection (single dose: 0.25 mg/kg of encapsulated mRNA in 10 mL/kg) of each tested LNP formulation per group. At 6 hours post dosing, 3 animals of each group underwent whole body live Bioluminescence imaging (BLI). For BLI, 3 animals per time point from each group received Luciferin (150mg/kg, i.p.) and underwent full body bioluminescence in the BERTHOLD TECHNOLOGIES NightOwl device 10 min later under isofluran narcosis. After whole body BLI, the 3 animals from each group were sacrificed and selected organs (lung, liver, heart, spleen,) were harvested for *ex vivo* BLI. The rest of 3 animals from each group were sacrificed by heart puncture at 24 hours post dosing, and blood plasma for cytokine level analysis was harvested. Concentrations of cytokines and chemokines in the animal plasma, collected at both 6 and 24 hours post dosing, were quantified using validated singleplex V-plex kits from Meso Scale Discovery (MSD) according to the manufacturer’s instruction, for IL-6 and MCP-1 (V-PLEX Mouse MCP-1 Kit and V-PLEX Mouse IL-6 Kit, respectively).

### Cryogenic Transmission Electron Microscopy (Cryo-TEM) of LNP

Frozen samples at a mRNA concentration of 0.5 mg/mL were thawed at room temperature and 4 μl of the undiluted sample was applied to an ATEM regularly spaced holey carbon copper grid (ATEM *LNPCFoil Grids*) in H_2_O saturated atmosphere at 4 °C and incubated for 1 min. Excess sample was blotted away, leaving a 100 nm thin layer of liquid on the surface of the grid. Blotted grids were vitrified immediately by plunge-freezing in liquid ethane at 180°C. Samples were stored in sample specific containers under liquid nitrogen (lN_2_) until further use in the cryo-electron microscope. After grid preparation, all grids were processed for transfer into the microscope and evaluated in a 200 kV ThermoFisher Glacios cryo-transmission electron microscope equipped with a Falcon IVi direct electron detector and ThermoFisher EPU software.

