## Supplementary Information for "Rational Design of Unsaturated, Thioether Ionizable Lipids for Enhanced *In vivo* mRNA Delivery"

### Content

**Synthesis and characterization of thioether lipids.** .......... 2

**Biological testing of LNPs containing novel thioether lipids.** .......... 12

### Synthesis and characterization of thioether lipids.

**Reagents used for the synthesis:**

Hept-1-yne, potassium thioacetate (KSAc), ammonia (7M), mesyl chloride, N,N-dimethylethane-1,2-diamine, N,N-dimethylaminoethylamine, 1-octadecanethiol, BOC-protected amines (various), pentylchloride, 5% Pd/$CaCO_3$, Pd/$BaSO_4$, ethanol, methanol, petroleum ether, dry methanol were purchased from Fisher Scientific (Schwerte, Germany). Dodecyl iodide was purchased from Sigma (Schnelldorf, Germany). N-methyl-1,2-ethylenediamine and trifluoroacetic acid (TFA), dichloromethane (DCM), anhydrous dimethylformamide, diethyl ether, acetic acid, sodium sulfate ($Na_2SO_4$), celite (for filtration), molecular sieves (for drying solvents) were bought from Carl Roth (Karlsruhe, Germany). Lindlar catalyst (5% Pd/$CaCO_3$ poisoned with Pd or Pd/$BaSO_4$) and sodium methylate were obtained from TCI (Eschborn, Germany). Quinoline was received from Merck KGaA (Darmstadt, Germany). Hydrogen gas was purchased from Air Liquide (Ludwigshafen, Germany). 2-(4-Iodobutoxy)tetrahydro-2H-pyran (1) and dibromide (25) were obtained from GenoSynth GmbH (Berlin, Germany). Oleyl alcohol, linoleyl alcohol ((9Z,12Z)-octadecadien-1-ol), $N^1$-BOC, $N^2$-methylethane-1,2-diamine, N-methyl-N-pentylamine, BOC-protected compound 30, Hexynol 33, dodecyl iodide, n-butyllithium, triisopropylsilylchloride were purchased from Sigma (Germany).

#### Methods

1. **Lipid synthesis**

*Synthesis of A1C11_D5*

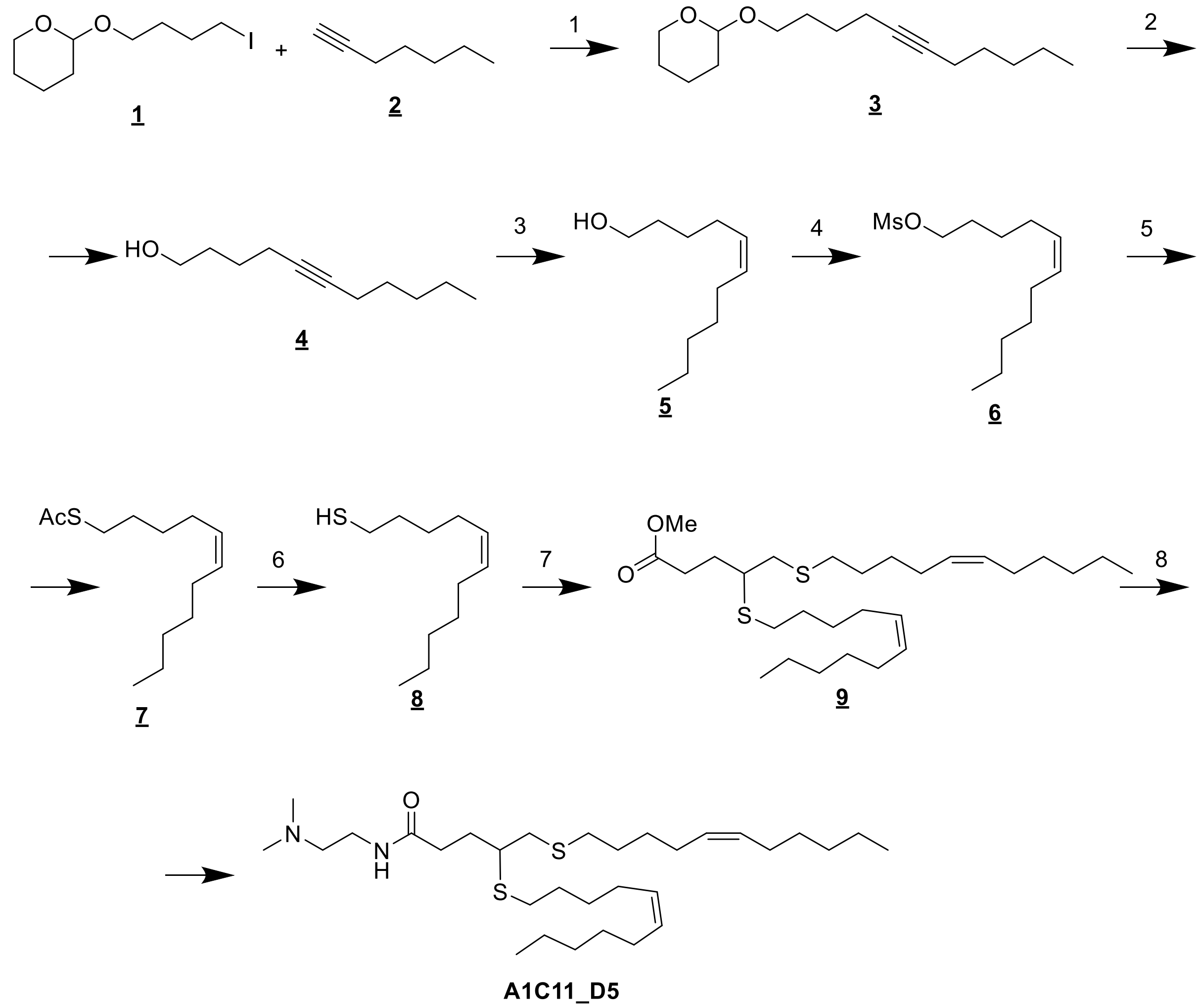


*Scheme 1. Synthesis scheme of A1C11_D5*

The synthesis began by alkylation of the hept-1-yne (**2**) with 2-(4-iodobutoxy)tetrahydro-2H-pyran (**1**) according to a published procedure.[1] This resulted in the formation of the protected alkyne **3**. Subsequent acid-catalyzed deprotection of the tetrahydropyranyl (THP) group afforded the corresponding alcohol **4**.

Alkyne **4** was then subjected to Lindlar reduction to selectively obtain the alkene **5**. The crude alkyne **4** (13.98 g, 83.08 mmol, 1.0 eq.) was added to a 1L three-necked reaction flask, followed by the addition of ethanol (500 mL) as solvent. To the reaction mixture, 5% Pd/$CaCO_3$ (700 mg, poisoned with Pd), Pd/BaSO4 (10% Pd, 700 mg), and quinoline (2 mL, 17.45 mmol, 0.21 eq.) were introduced under a nitrogen atmosphere. The system was purged with hydrogen gas, and the reaction was stirred at room temperature until completion. Upon completion, the reaction mixture was filtered through celite and concentrated under reduced pressure. The crude product was purified by flash chromatography, yielding the alkene **5** as a yellow oil with a 48% yield.

Alcohol **5** was mesylated using mesyl chloride to produce the mesylated compound **6**. Subsequently, **6** was reacted with potassium thioacetate (KSAc) to yield AcS-derivative **7**. The detailed procedure is as follows: In a 100 mL round-bottomed flask equipped with magnetic stirring, mesylate **6** (11.99 g, 48.3 mmol, 1 eq) was dissolved in anhydrous DMF (50 mL). Potassium thioacetate (11 g, 96 mmol, 2 eq) was added to the reaction mixture in one portion, and the reaction was stirred overnight at room temperature. Upon completion, the mixture was poured into water (500 mL) and extracted with diethyl ether (2 x 200 mL). The organic layers were dried over sodium sulfate ($Na_2SO_4$), and the solvent was removed under reduced pressure to afford the crude product **7**.

The obtained thioester **7** was hydrolyzed to yield thiol **8** according to the procedure: In a 100 mL round-bottomed flask with magnetic stirring, alkenyl thioacetate (11.2 g, 49 mmol, 1 eq) was dissolved in methanol (MeOH) containing 7 M ammonia (28 mL). The reaction was stirred at room temperature overnight. After completion, the reaction mixture was concentrated under reduced pressure and purified by flash chromatography, eluting with neat petroleum ether (PE), to yield thiol **8** as a colorless oil (76% yield).

Compound **9** was synthesized by coupling alkenyl thiol **8** with dibromide **25**. In a 500 mL round-bottomed flask with magnetic stirring, sodium methylate (2.17 g, 40.14 mmol, 2.2 eq) was suspended in dry methanol (80 mL, dried over molecular sieves). Alkenyl thiol **8** (7.48 g, 40.14 mmol, 2.2 eq), dissolved in diethyl ether (80 mL, dried over molecular sieves), was added dropwise to the sodium methylate suspension. The mixture was stirred for 30 minutes at room temperature. Dibromide **25** (5 g, 18.24 mmol, 1.0 eq) was then added in one portion, and the reaction was stirred overnight while monitoring progress by TLC. Afterward, acetic acid (1 mL) was added, and the solvent was evaporated under reduced pressure to yield the crude ester **9**.

The crude methyl ester **9** was directly subjected to amidation with neat N,N-dimethylaminoethylamine. The ester was dissolved in the amine, and the reaction mixture was heated at 60°C overnight. Upon completion, the solvent was evaporated under reduced pressure, and the crude product was purified by flash chromatography to afford the target lipid **A1C11_D5**.

*Synthesis of A1C18_D9*

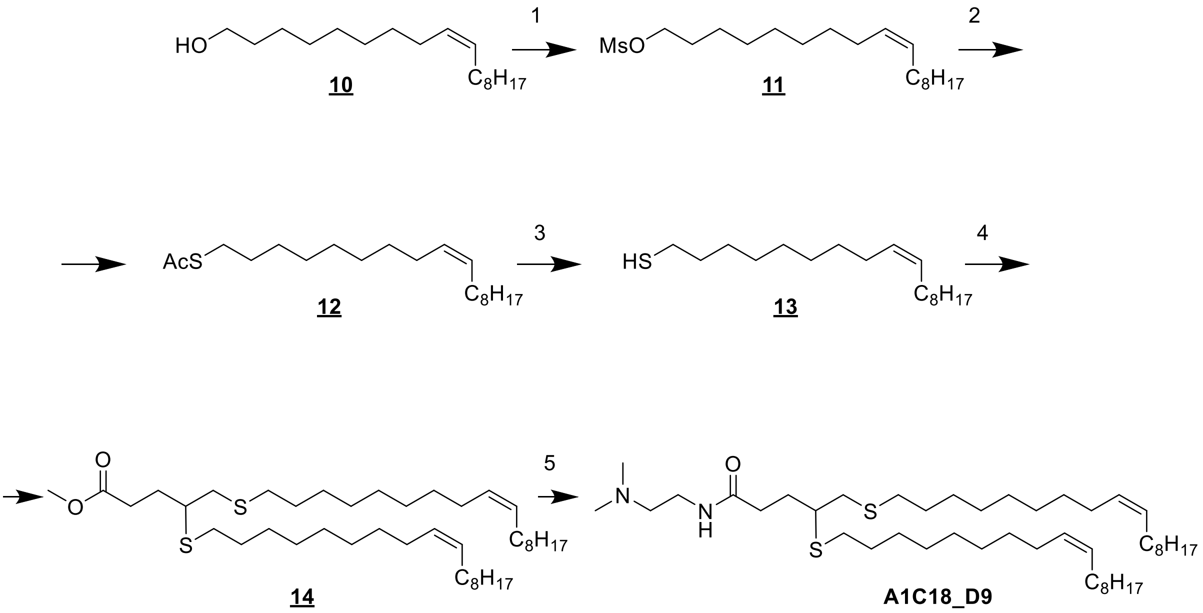


*Scheme 2. Synthesis scheme for A1C18_D9*

**A1C18_D9** was synthesized following a procedure analogous to that used for **A1C11_D5**, beginning with commercially available oleyl alcohol **<u>10</u>**. The alcohol was first mesylated using mesyl chloride, yielding mesylated intermediate **<u>11</u>**. This intermediate was then treated with potassium thioacetate (KSAc), resulting in the formation of the AcS-derivative **<u>12</u>**. Subsequent hydrolysis of compound **<u>12</u>** produced thiol **<u>13</u>**, which was further reacted with dibromide **<u>25</u>**. The final step involved direct amidation with dimethylaminoethyl, leading to the target lipid, **<u>A1C18_D9</u>**.

*Synthesis of A1C18_D9_D12*

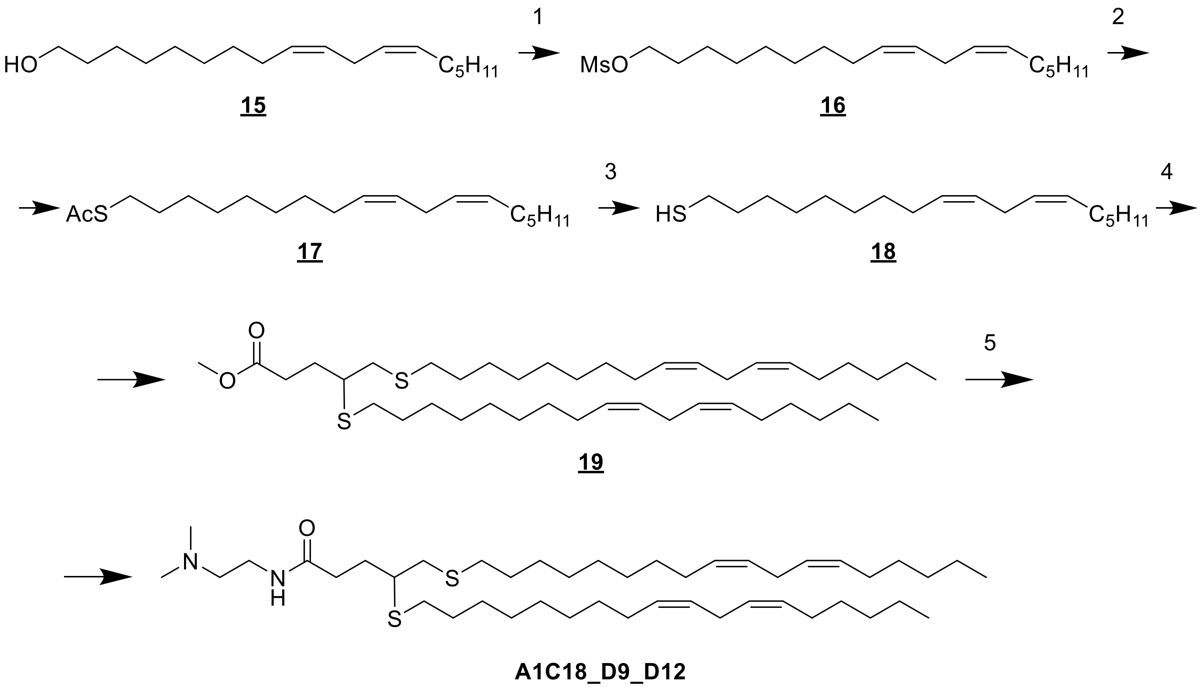


*Scheme 3. Synthesis scheme of A1C18_D9_D12*

**A1C18_D9_D12** was synthesized following the same procedure used for **A1C11_D5**, starting from linoleyl alcohol **15** ((9Z,12Z)-octadecadien-1-ol) as the initial material.

*Synthesis of A1C18*

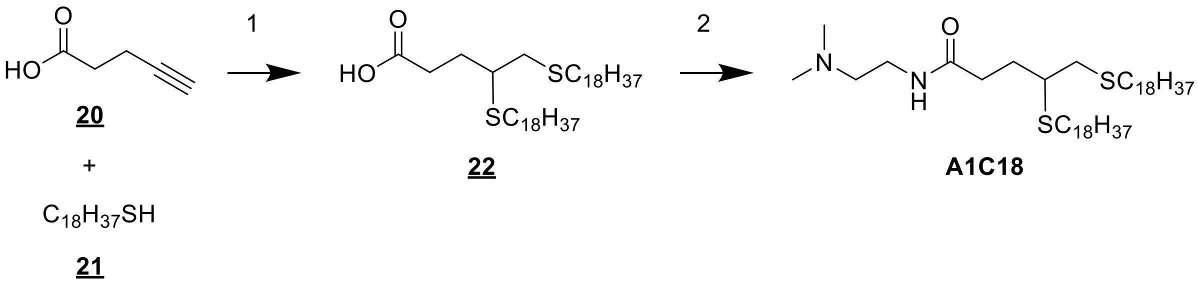


*Scheme 4. Synthesis scheme of A1C18*

**A1C18** was synthesized following a published two-step procedure. In the first step, alkyne **20** was reacted with 1-octadecanethiol through a UV-induced thiol-yne reaction. This was followed by the amidation of the resulting carboxylic acid with the corresponding amine, as previously described,[2] to give ***A1C18***.

*Synthesis of precursors, dibromide **25**, and amines **28** and **32***

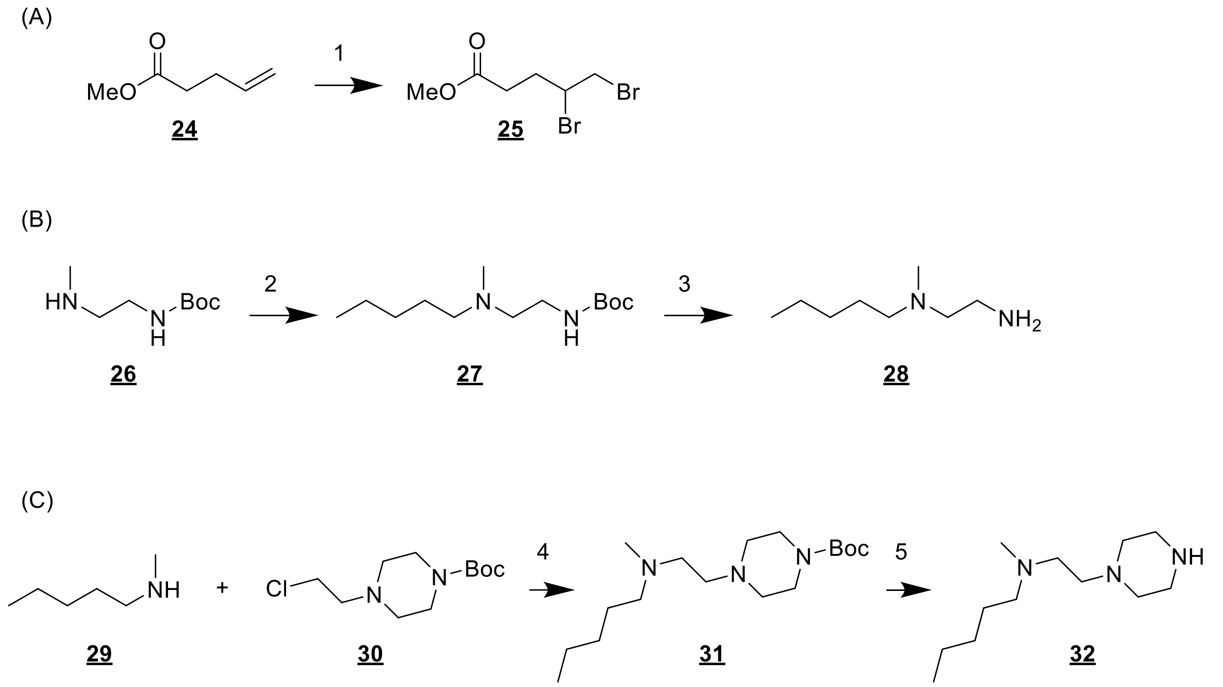


*Scheme 5. Synthesis scheme of dibromide 25, and amines 28 and 32*

Dibromide **25** was synthesized by addition of brom to alkene **24** following a standard procedure.[3]

Amine **28** was prepared by alkylating N[1]-BOC, N[2]-methylethane-1,2-diamine **26** with pentylchloride, followed by BOC deprotection using trifluoroacetic acid (TFA) in dichloromethane (DCM), yielding amine **28**.

In a similar manner, amine **32** was synthesized by alkylating *N*-methylpentan-1-amine **29** with the BOC-protected compound **30**. Subsequent BOC deprotection yielded the desired amine **32**.

*Synthesis of A1C18_D5, A2C18_D5, A3C18_D5 and A4C18_D5*

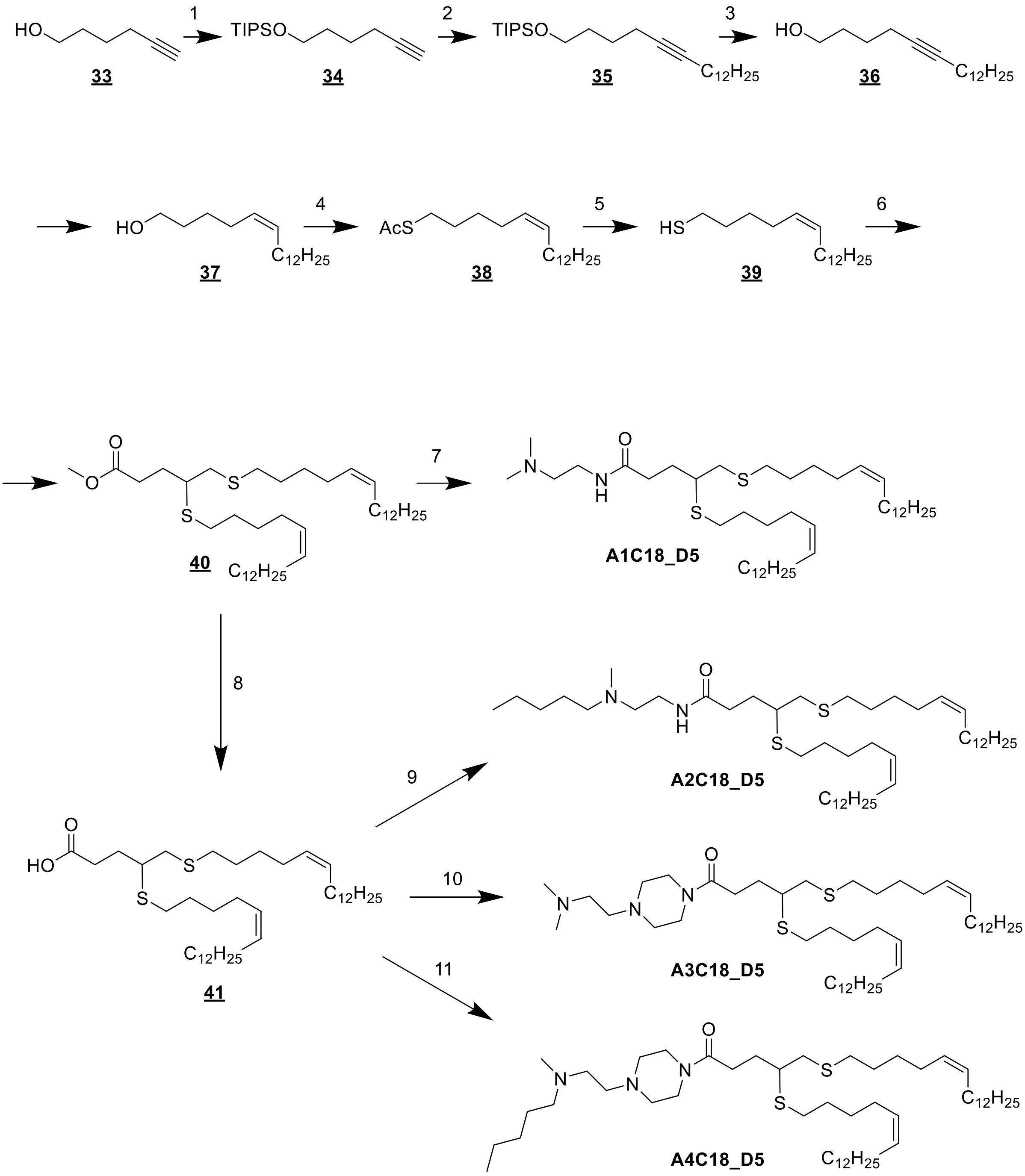


*Scheme 6. Synthesis schemes for A1C18_D5, A2C18_D5, A3C18_D5 and A4C18_D5*

Hexynol **33** was initially protected with a triisopropylsilyl (TIPS) group, followed by alkylation with dodecyl iodide ($C_{12}H_{25}I$) using n-butyllithium to yield alkyne **35**. Afterward, the TIPS group was removed to produce octadec-5-yn-1-ol **36**, which was then selectively reduced to (*Z*)-octadec-5-en-1-ol **37** using a Lindlar catalyst, following the procedure previously described for the synthesis of alkene **5**. Octadecenol **37** was converted to (Z)-octadec-5-ene-1-thiol **39** and subsequently transformed into methyl ester **40** bearing two C18_D5 tails, as outlined in the procedure used for compound **9**. Methyl ester **40** was hydrolyzed to obtain carboxylic acid **41**, which was then subjected to amidation with various amines — **28**, N,N-dimethyl-2-(piperazin-1-yl)ethan-1-amine, and amine **32** —to synthesize lipids **A2C18_D5**, **A3C18_D5**, and

**A4C18_D5**, respectively. For the synthesis of lipid **A1C18_D5**, methyl ester **40** was directly amidated with N,N-dimethylethane-1,2-diamine using the same method described earlier for compound **A1C11_D5**.

*Synthesis of A2C18_D9*

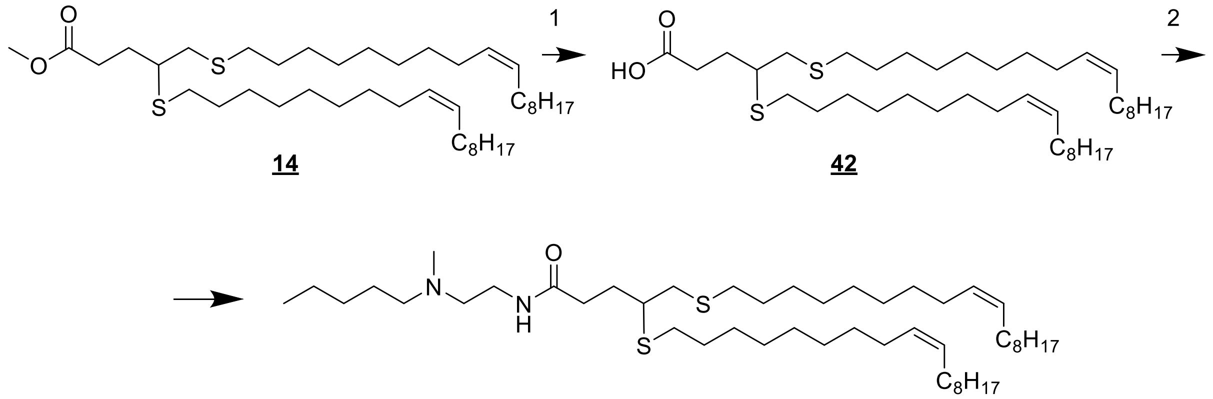


**14** **42** **A2C18_D9**

*Scheme 7. Synthesis scheme of A2C18_D9*

Lipid **A2C18_D9** was synthesized from methyl ester **14** by its hydrolysis to the carboxylic acid **42**, followed by its amidation using amine **28** to give the final lipid.

*Synthesis of A2C18_D9_D12*

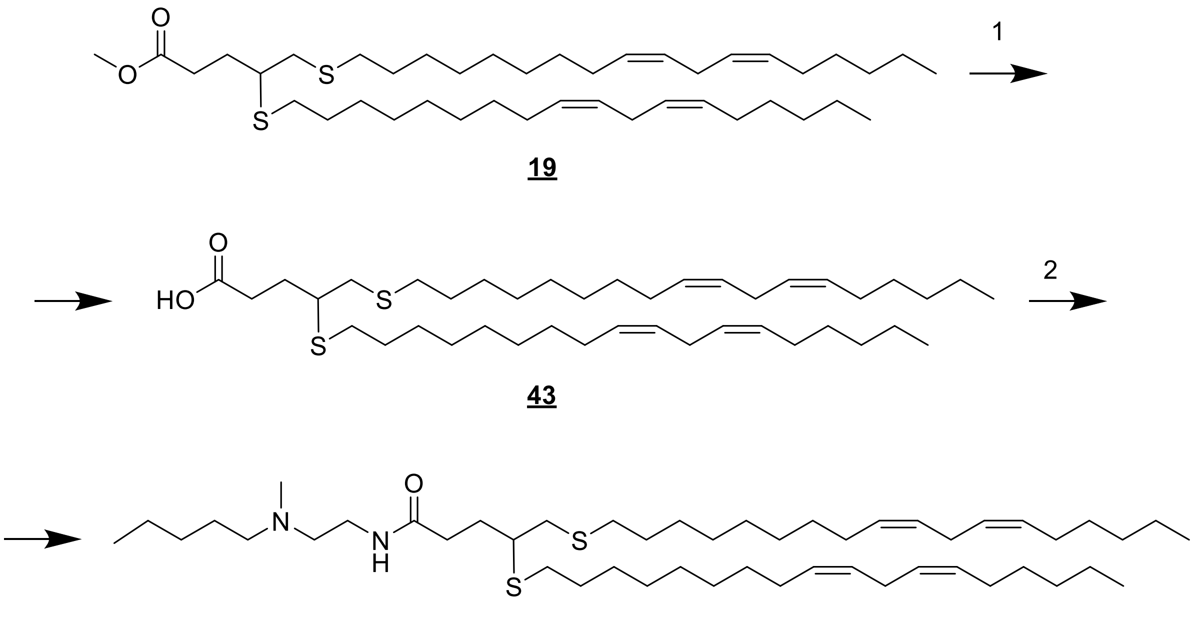


**A2C18_D9_D12**

*Scheme 8. Synthesis scheme of A2C18_D9_D12*

Lipid **A2C18_D9_D12** was synthesized from methyl ester **19** by its hydrolysis to the carboxylic acid **43**, followed by its amidation using the amine **28** to give the final lipid.

NMR spectra and LC-MS data of final products and selected intermediates can be found at the end of the Supporting Information.

*Table 1: Overview of lipids tested in the current work.*

| Name | Structure |
|---|---|
| **DLin-MC3-DMA** | 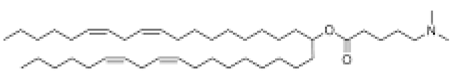 |
| **A1C11** | 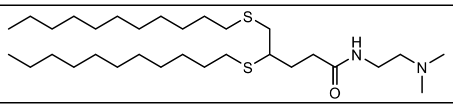 |
| **A1C11_D5** | 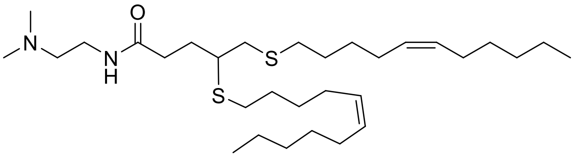 |
| **A1C18_D9** | 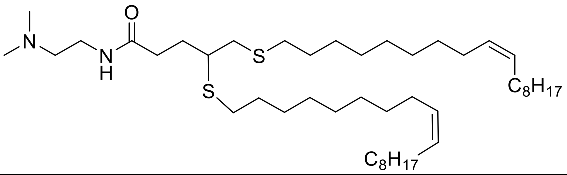 |

| | |
|---|---|
| **A1C18_D9_D12** | 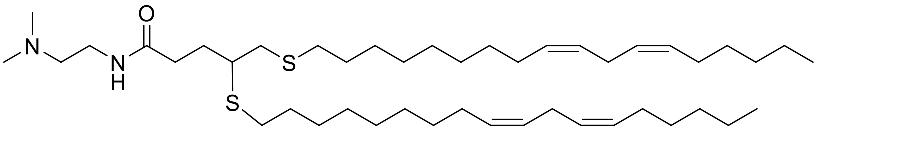 |
| **A1C18_D5** | 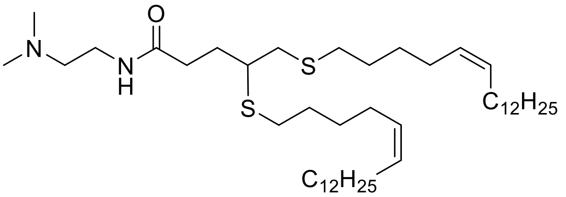 |
| **A1C18** | 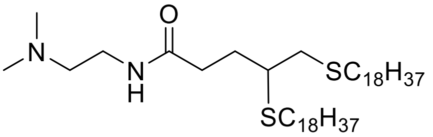 |
| **A2C18_D5** | 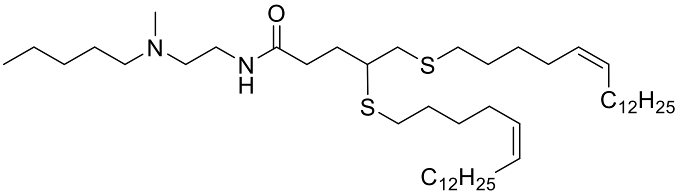 |
| **A3C18_D5** | 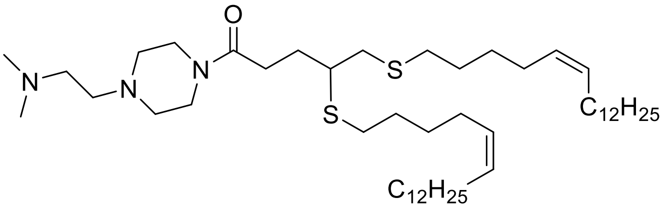 |
| **A4C18_D5** | 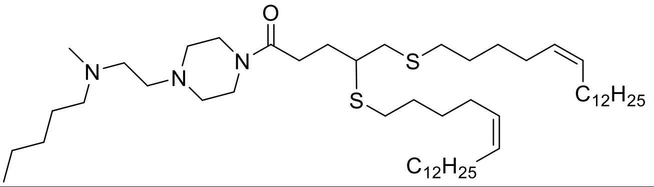 |
| **A2C18_D9** | 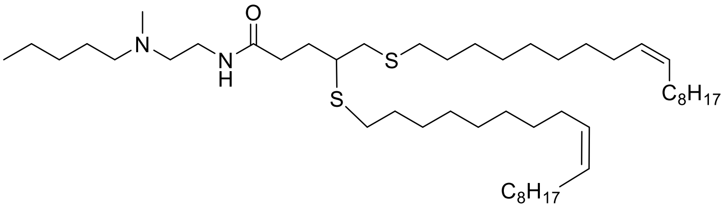 |
| **A2C18_D9_D12** | 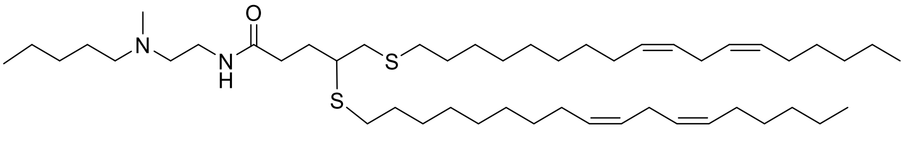 |

### Biological testing of LNPs containing novel thioether lipids.

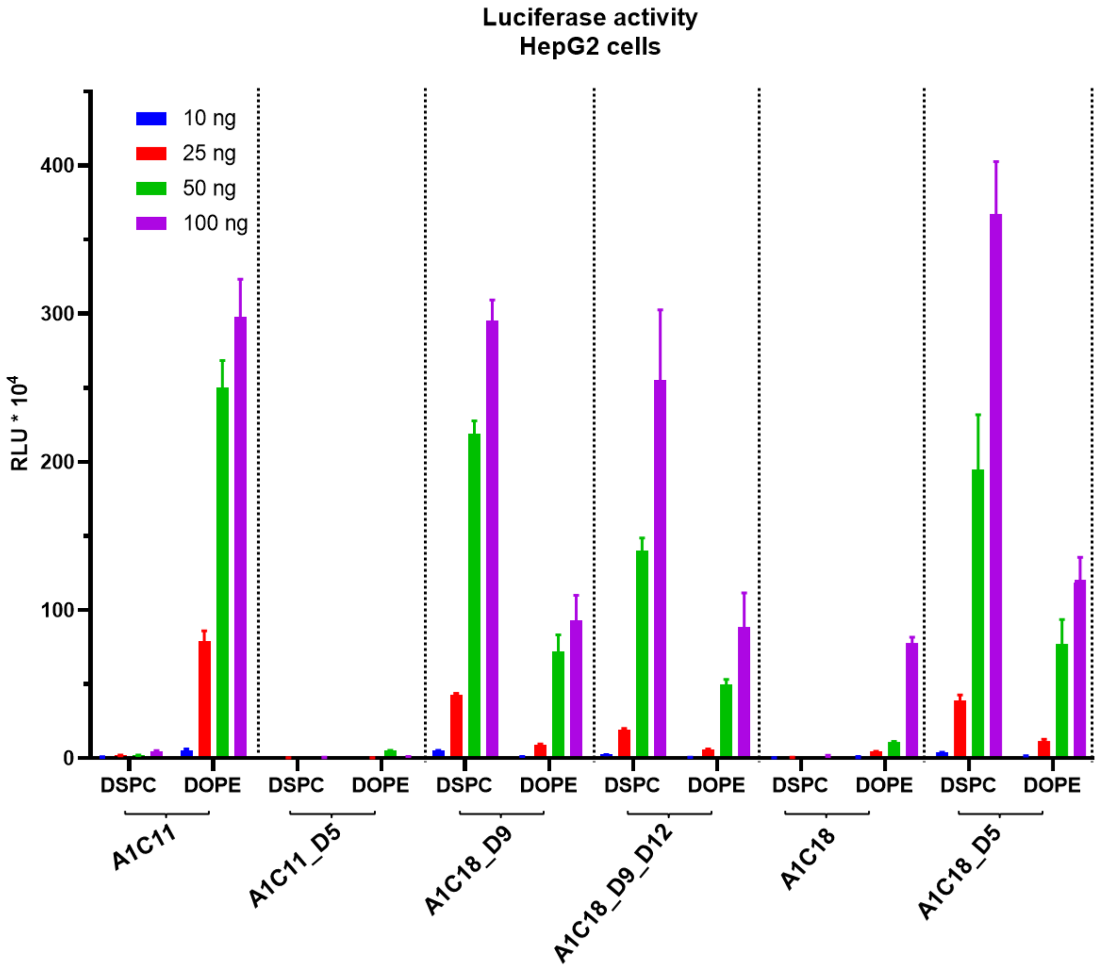


*Figure 1: HepG2 cells were treated with LNPs containing mRNA encoding for Luciferase (at four different mRNA doses) for 24 h and the luciferase activity was determined after 24 h. Selected data was extracted from this graph and is shown in Figure 1. Mean values from triplicates are shown ± standard deviations.*

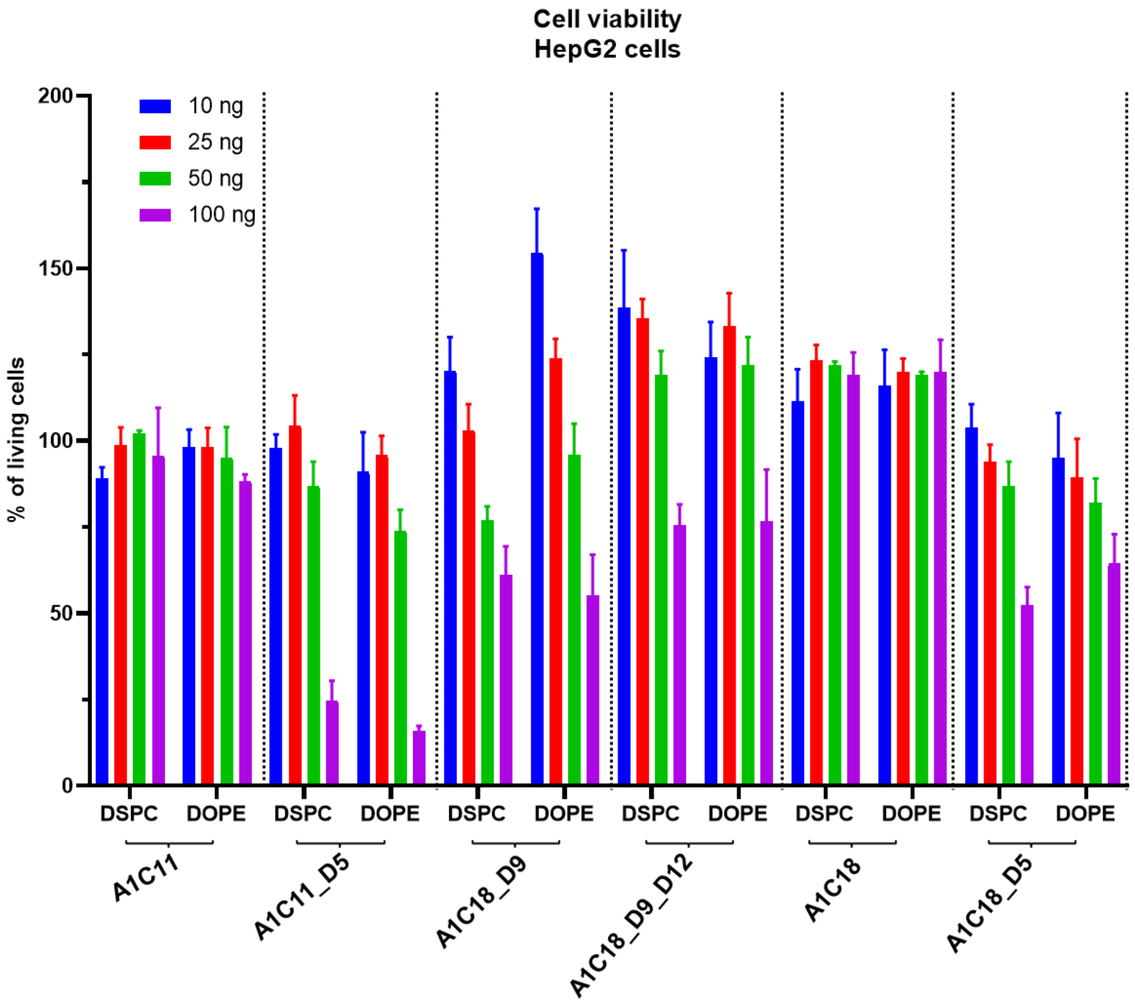


*Figure 2: HepG2 cells were treated with LNPs containing mRNA encoding for Luciferase (at four different mRNA doses) for 24 h and the cell viability was determined afterwards. Untreated cells were used as control and the cell viability was set to 100%. Mean values from triplicates are shown ± standard deviations.*

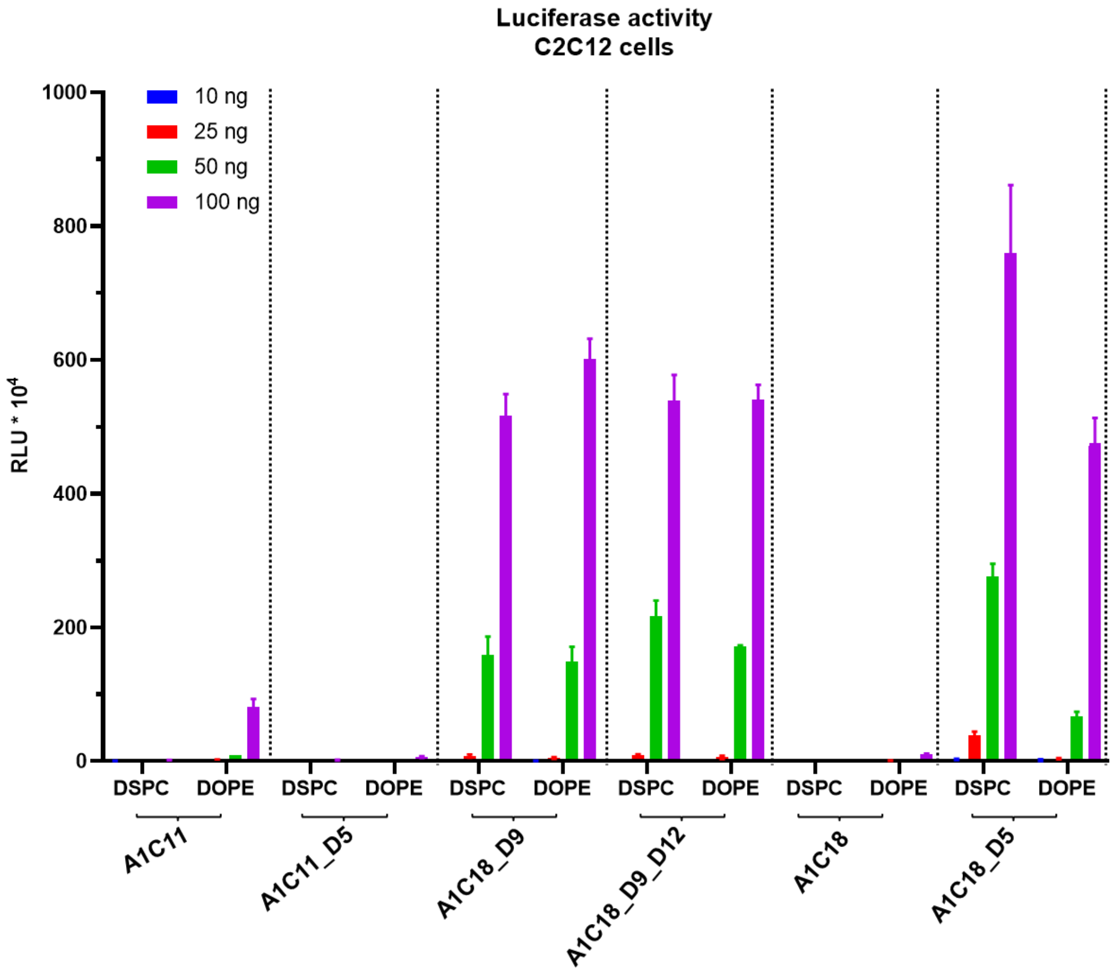


*Figure 3: C2C12 cells were treated with LNPs containing mRNA encoding for Luciferase (at four different doses 10 – 100 ng) for 24 h and the luciferase activity was determined afterwards. Selected data was extracted from this graph and is shown in Figure 1. Mean values from triplicates are shown ± standard deviations.*

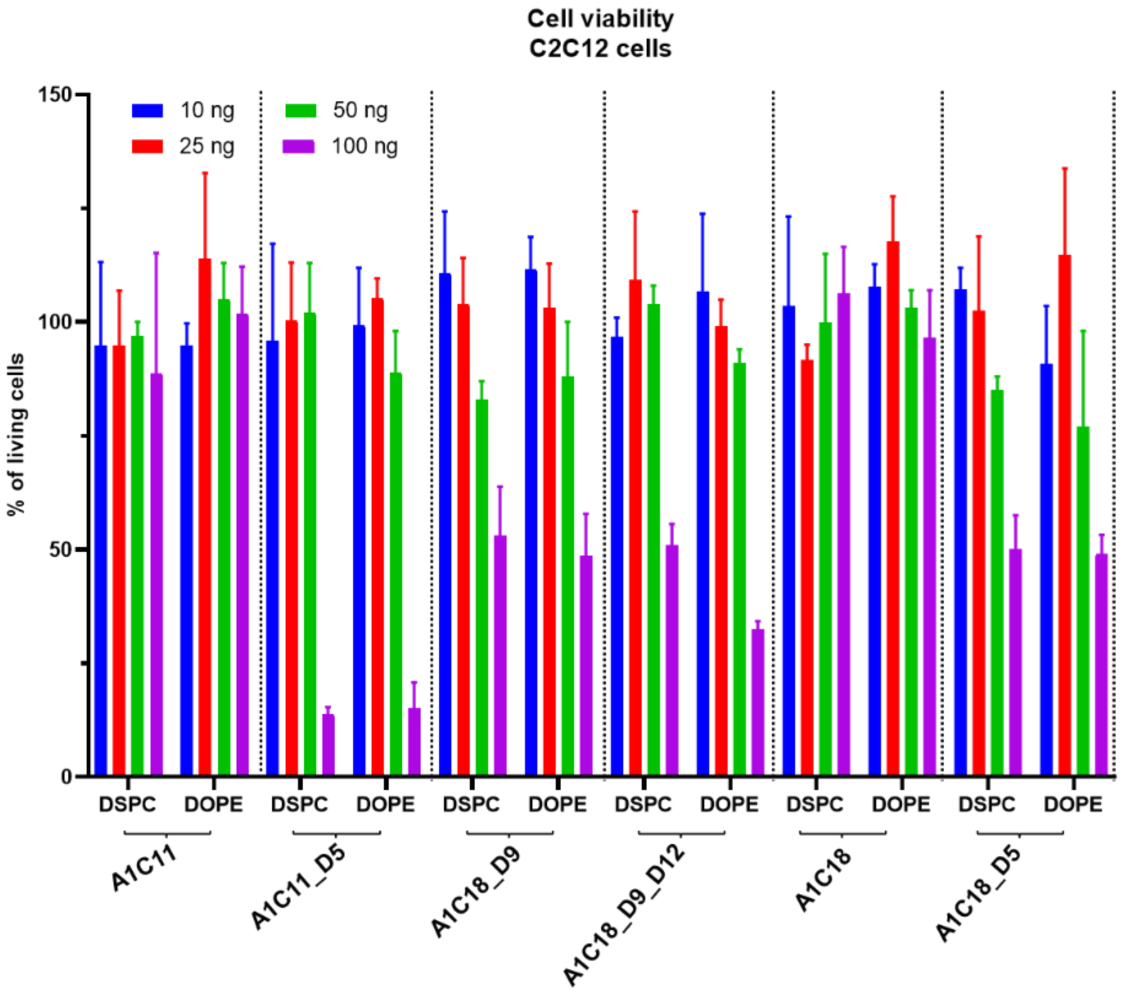


*Figure 4: C2C12 cells were treated with LNPs containing mRNA encoding for Luciferase (at four different doses 10 – 100 ng) for 24 h and the cell viability was determined afterwards. Untreated cells were used as control and the cell viability was set to 100%. Mean values from triplicates are shown ± standard deviations.*

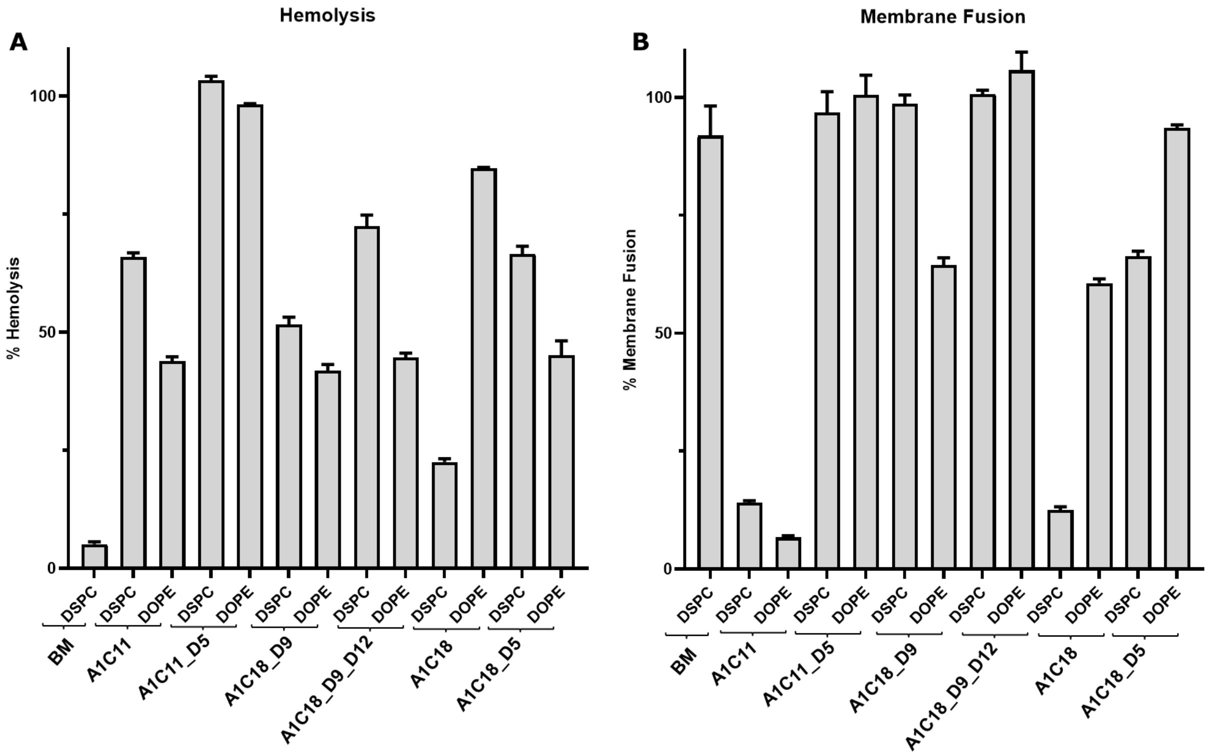


*Figure 5: Red blood cells were incubated in PBS at pH = 7.4 (A) or in buffer at pH = 5.5 with LNPs for 1 h. The absorption was measured afterwards at 540 nm. Selected data was extracted from this graph and is shown in Figure 2. Mean values from duplicates are shown ± standard deviations.*

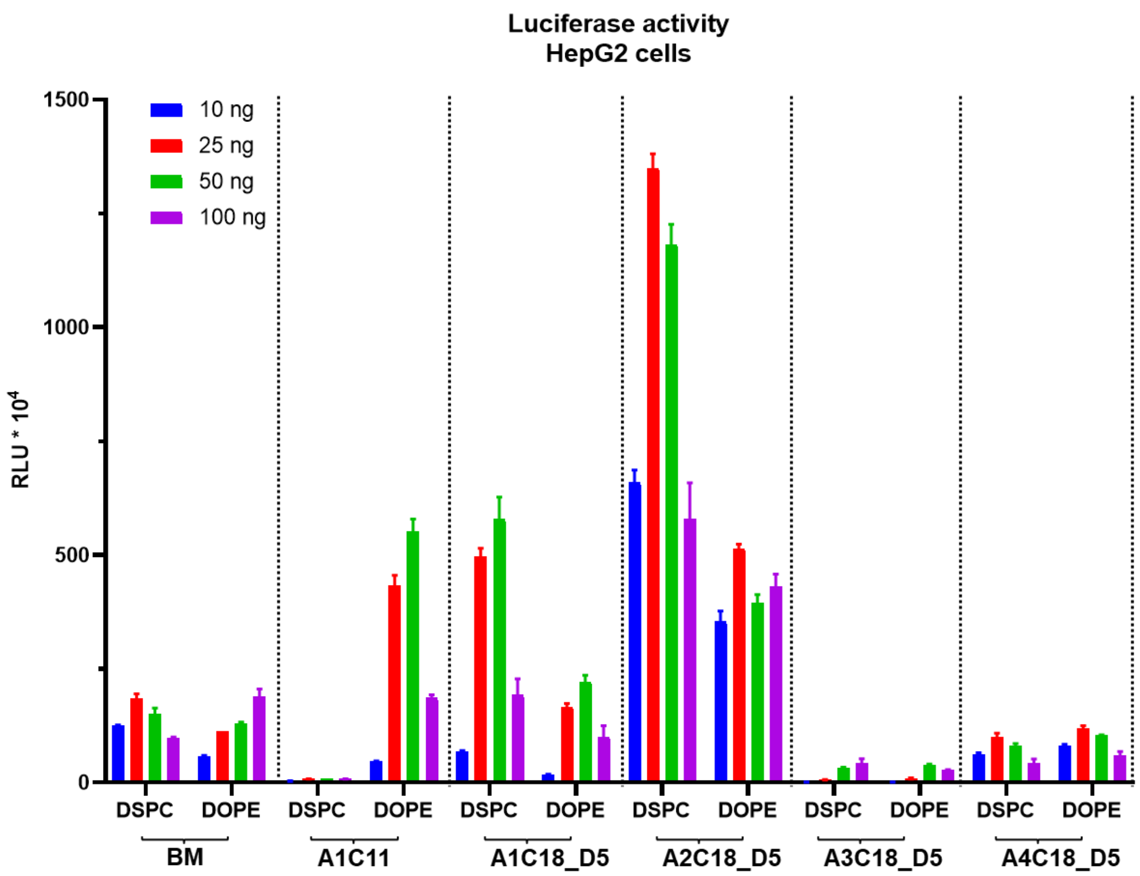


*Figure 6: HepG2 cells were treated with LNPs containing mRNA encoding for Luciferase (at four different doses 10 – 100 ng) for 24 h and the luciferase activity was determined afterwards. Mean values from triplicates are shown ± standard deviations. BM = Benchmark LNP*

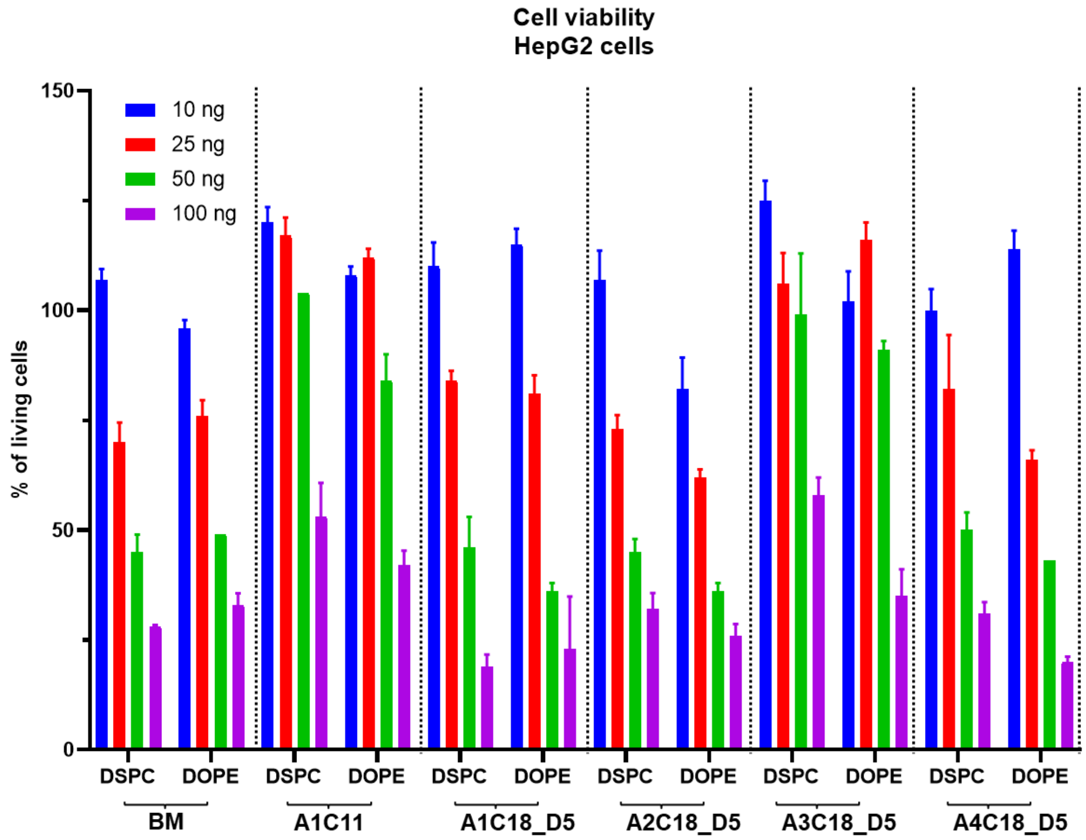


*Figure 7: HepG2 cells were treated with LNPs containing mRNA encoding for Luciferase (at four different doses 10 – 100 ng) for 24 h and the cell viability was determined afterwards. Untreated cells were used as control and the cell viability was set to 100%. Mean values from triplicates are shown ± standard deviations. BM = Benchmark LNP*

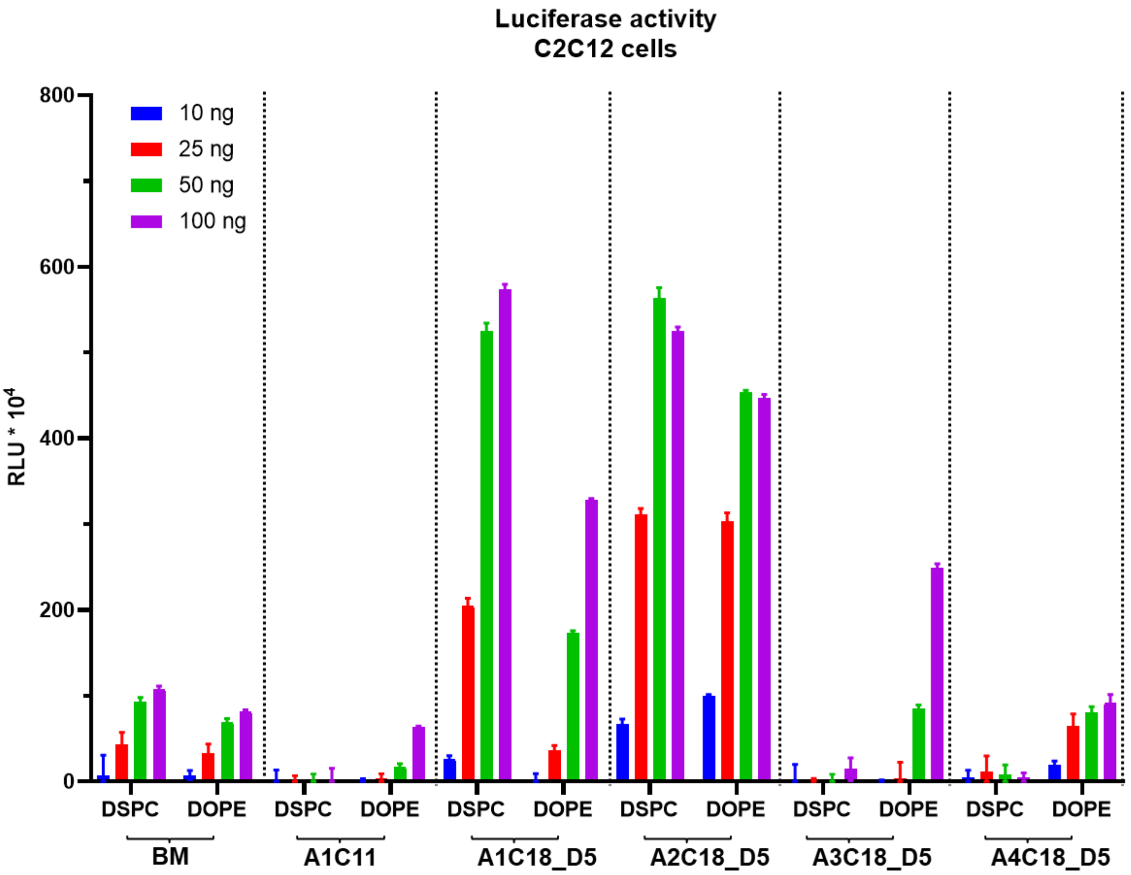


*Figure 8: C2C12 cells were treated with LNPs containing mRNA encoding for Luciferase (at four different doses 10 – 100 ng) for 24 h and the luciferase activity was determined afterwards. Mean values from triplicates are shown ± standard deviations. BM = Benchmark LNP*

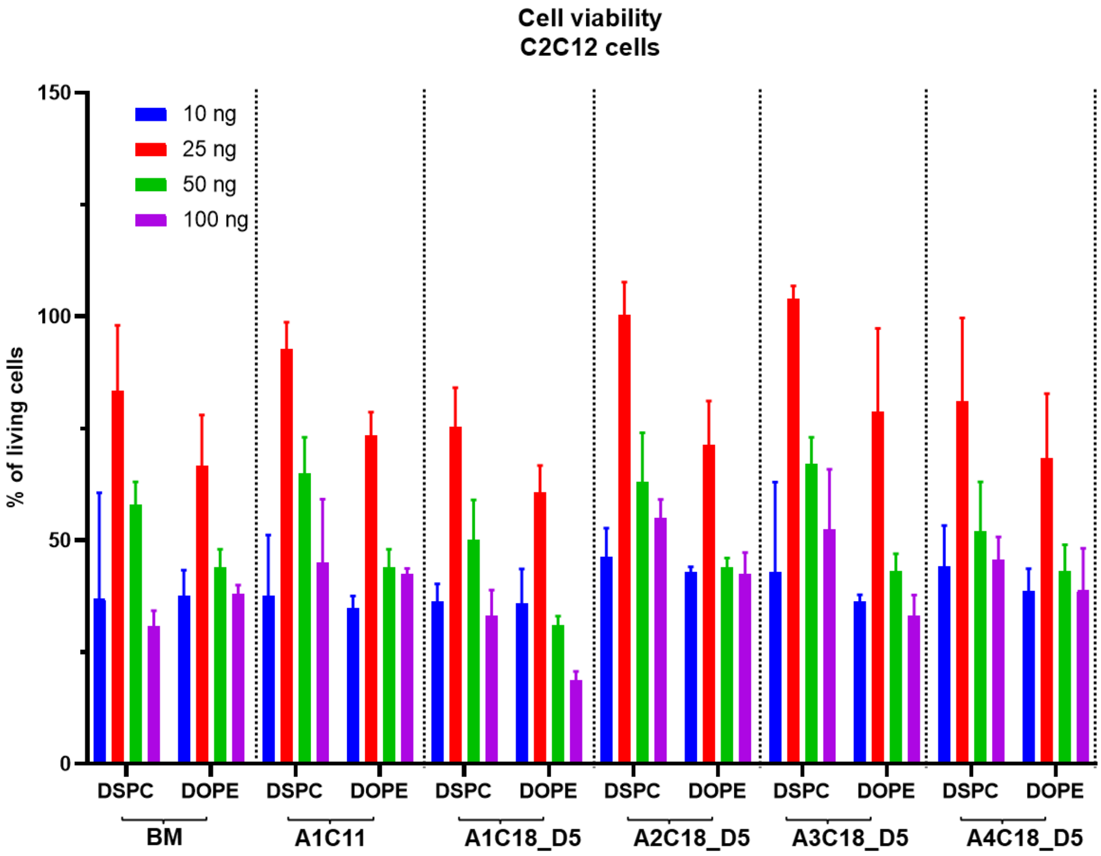


*Figure 9: C2C12 cells were treated with LNPs containing mRNA encoding for Luciferase (at four different doses 10 – 100 ng) for 24 h and the cell viability was determined afterwards. Untreated cells were used as control and the cell viability was set to 100%. Mean values from triplicates are shown ± standard deviations. BM = Benchmark LNP*

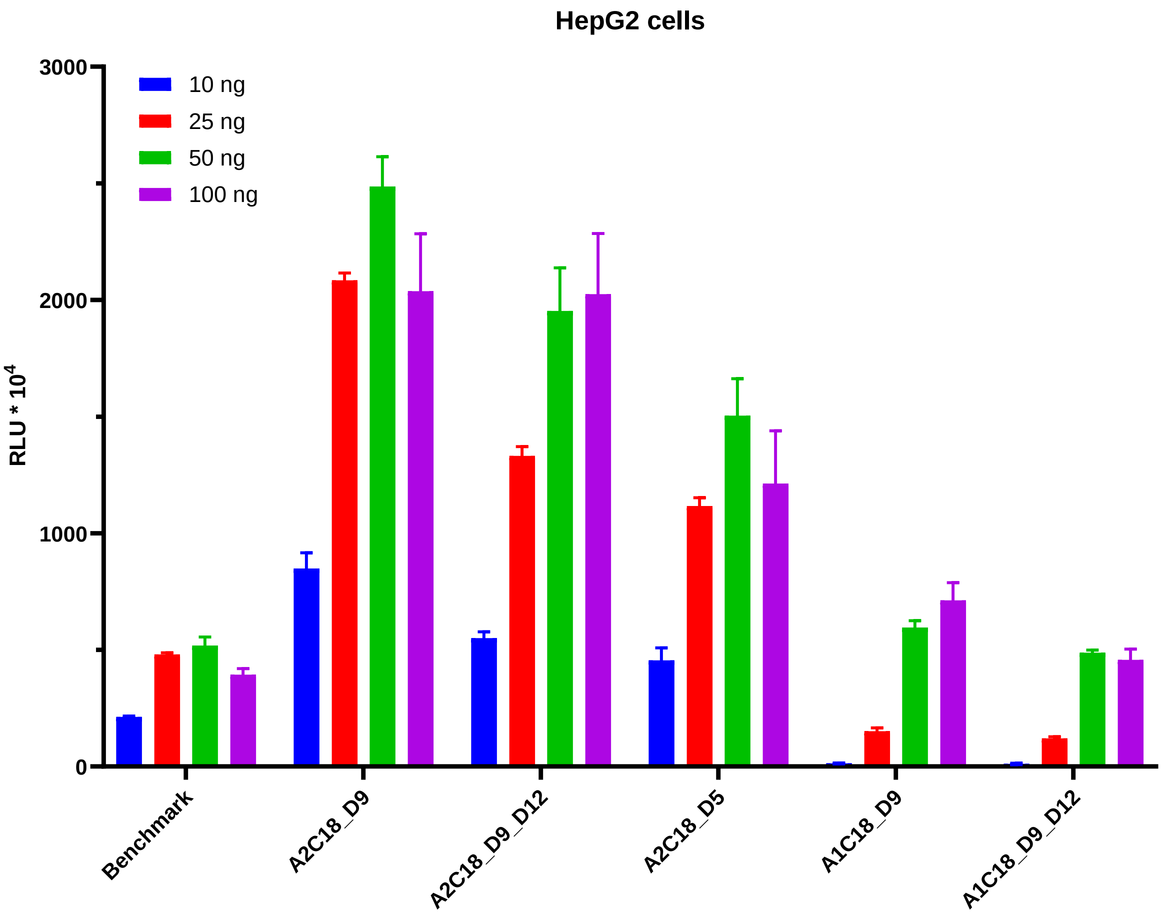


*Figure 10: HepG2 cells were treated with LNPs containing mRNA encoding for Luciferase (at four different doses 10 – 100 ng) for 24 h and the cell viability was determined afterwards. Untreated cells were used as control and the cell viability was set to 100%. Selected data was extracted from this graph and is shown in Figure 3. Mean values from triplicates are shown ± standard deviations.*

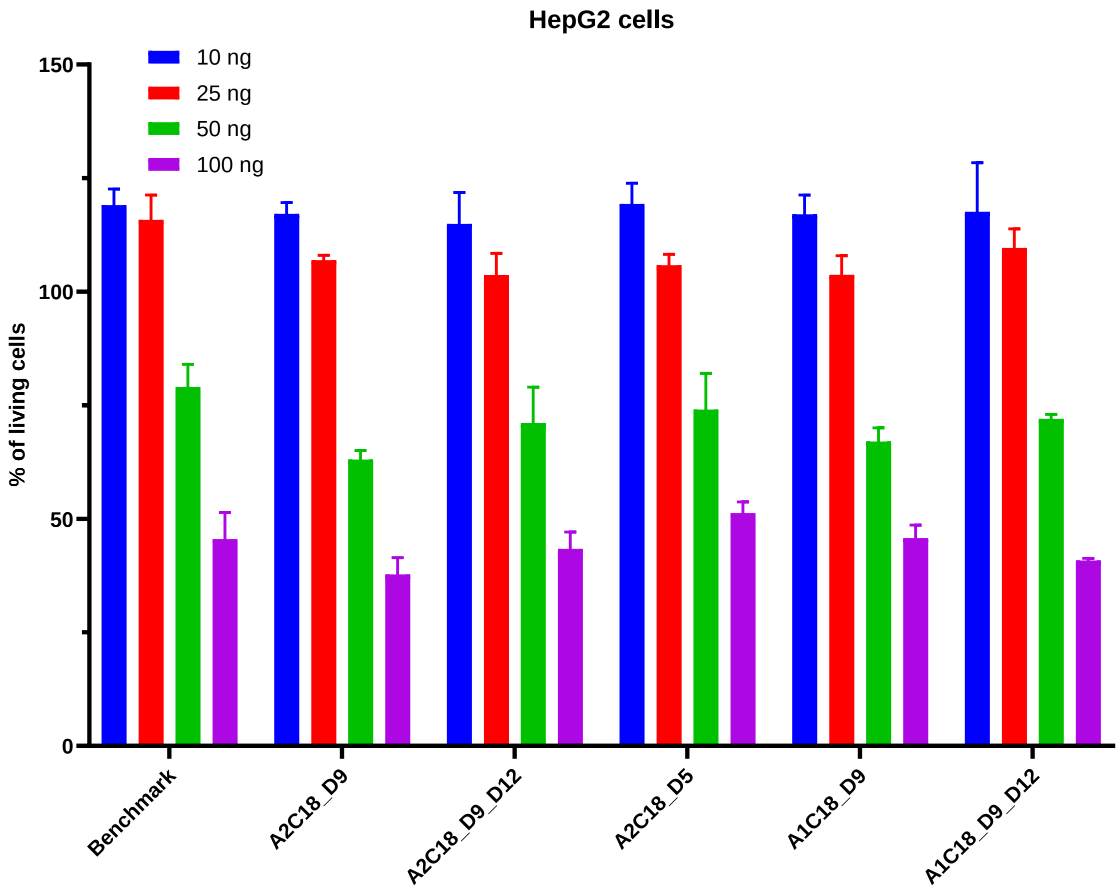


*Figure 11: HepG2 cells were treated with LNPs containing mRNA encoding for Luciferase (at four different doses 10 – 100 ng) for 24 h and the cell viability was determined afterwards. Untreated cells were used as control and the cell viability was set to 100%. Mean values from triplicates are shown ± standard deviations.*

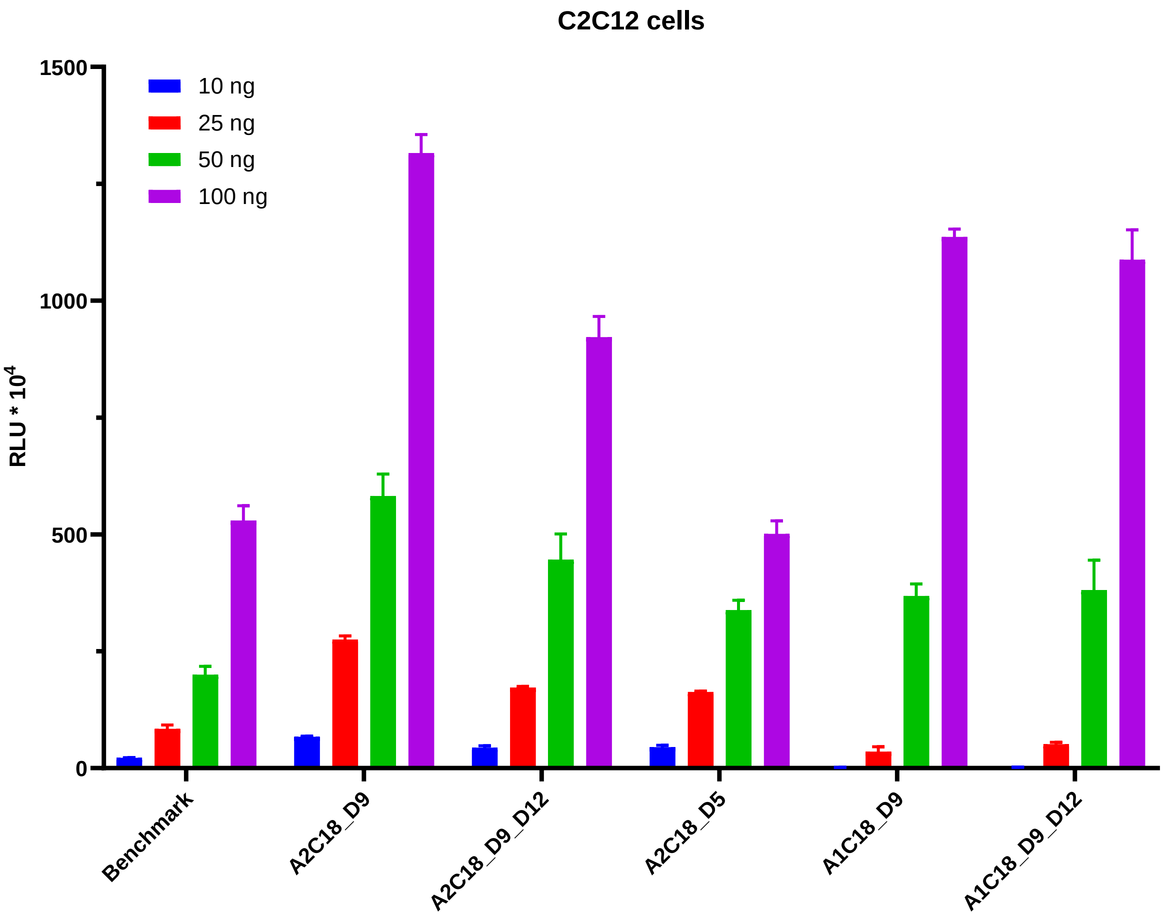


*Figure 12: C2C12 cells were treated with LNPs containing mRNA encoding for Luciferase (at four different doses 10 – 100 ng) for 24 h and the luciferase activity was determined afterwards. Mean values from triplicates are shown ± standard deviations.*

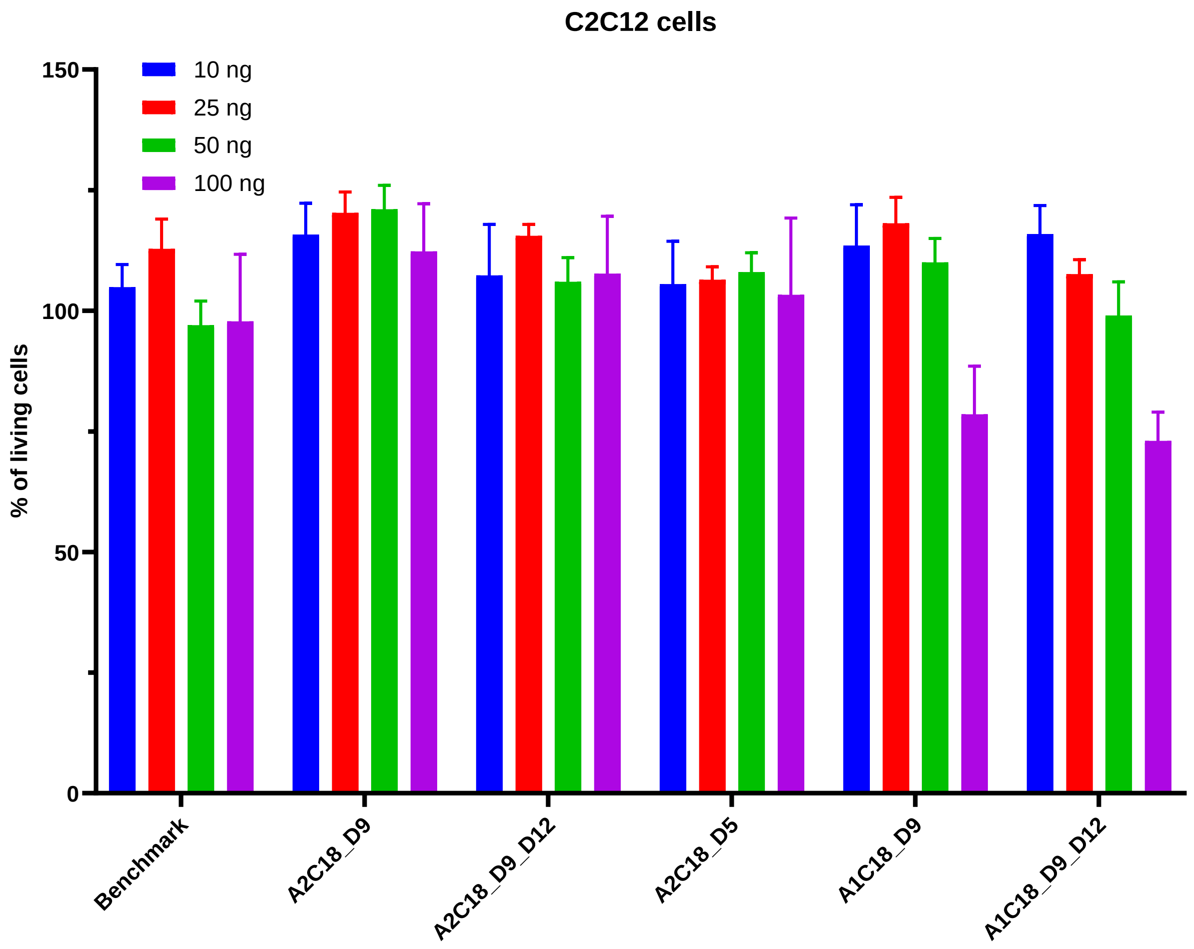


*Figure 13: C2C12 cells were treated with LNPs containing mRNA encoding for Luciferase (at four different doses 10 – 100 ng) for 24 h and the cell viability was determined afterwards. Untreated cells were used as control and the cell viability was set to 100%. Mean values from triplicates are shown ± standard deviations.*

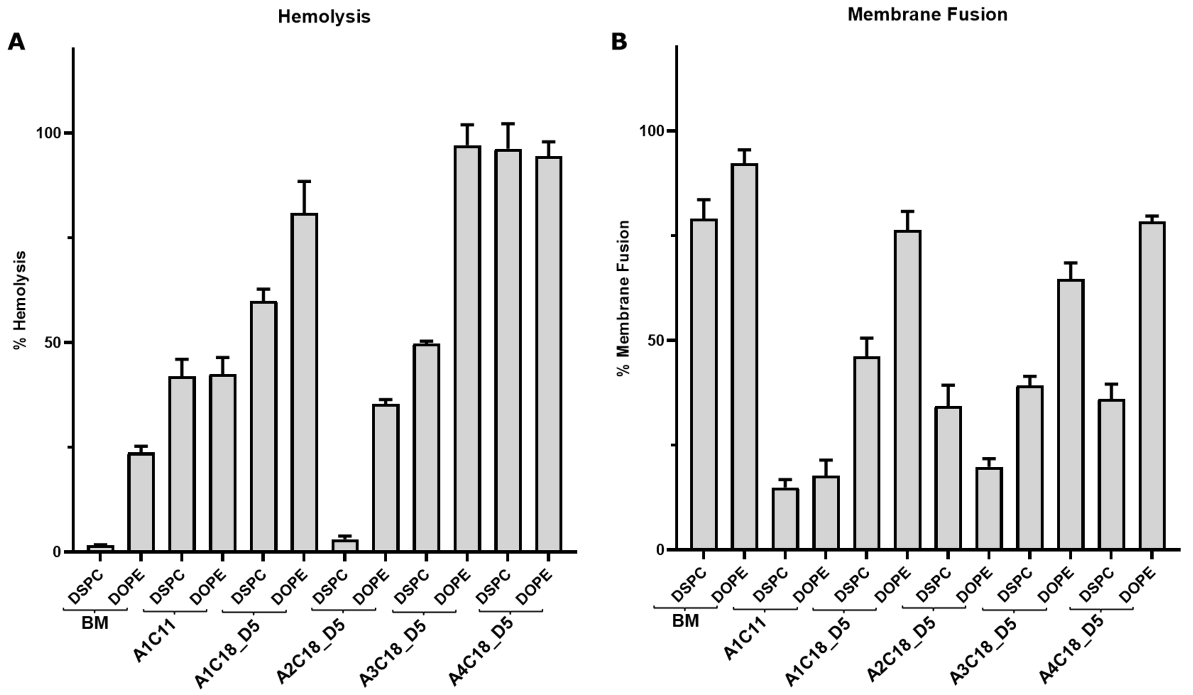


*Figure 14: Red blood cells were incubated in PBS at pH = 7.4 (A) or in buffer at pH = 5.5 with LNPs for 1 h. The absorption was measured afterwards at 540 nm. Selected data was extracted from this graph and is shown in Figure 3. Mean values from duplicates are shown ± standard deviations.*

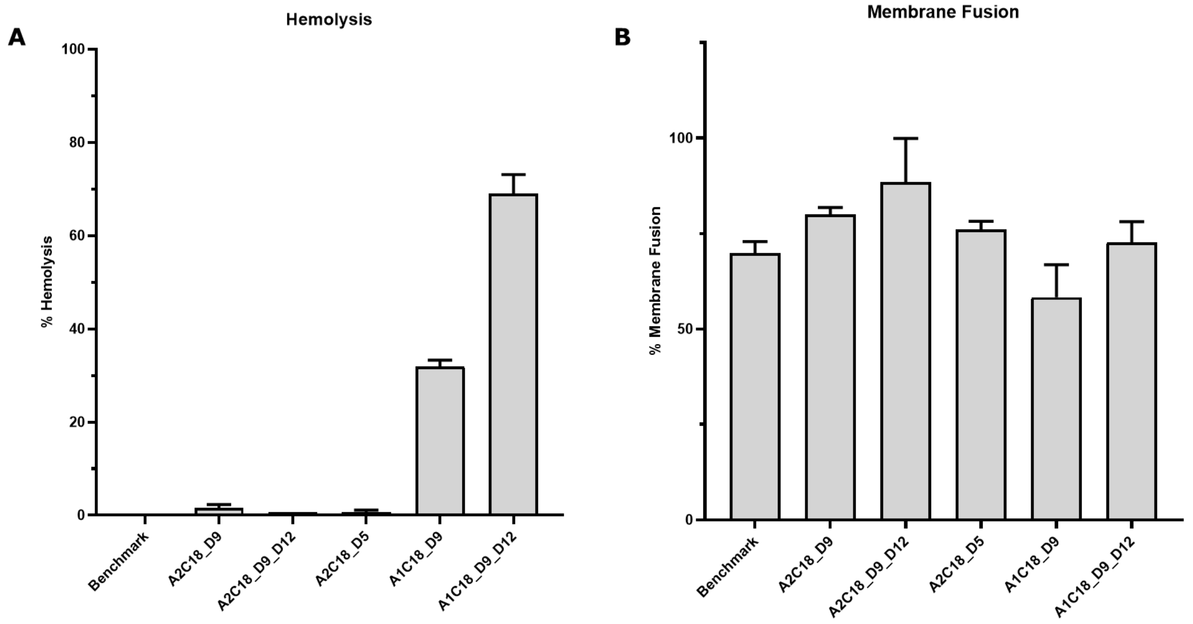


*Figure 15: Red blood cells were incubated in PBS at pH = 7.4 (A) or in buffer at pH = 5.5 with LNPs for 1 h. The absorption was measured afterwards at 540 nm. Mean values from duplicates are shown ± standard deviations.*

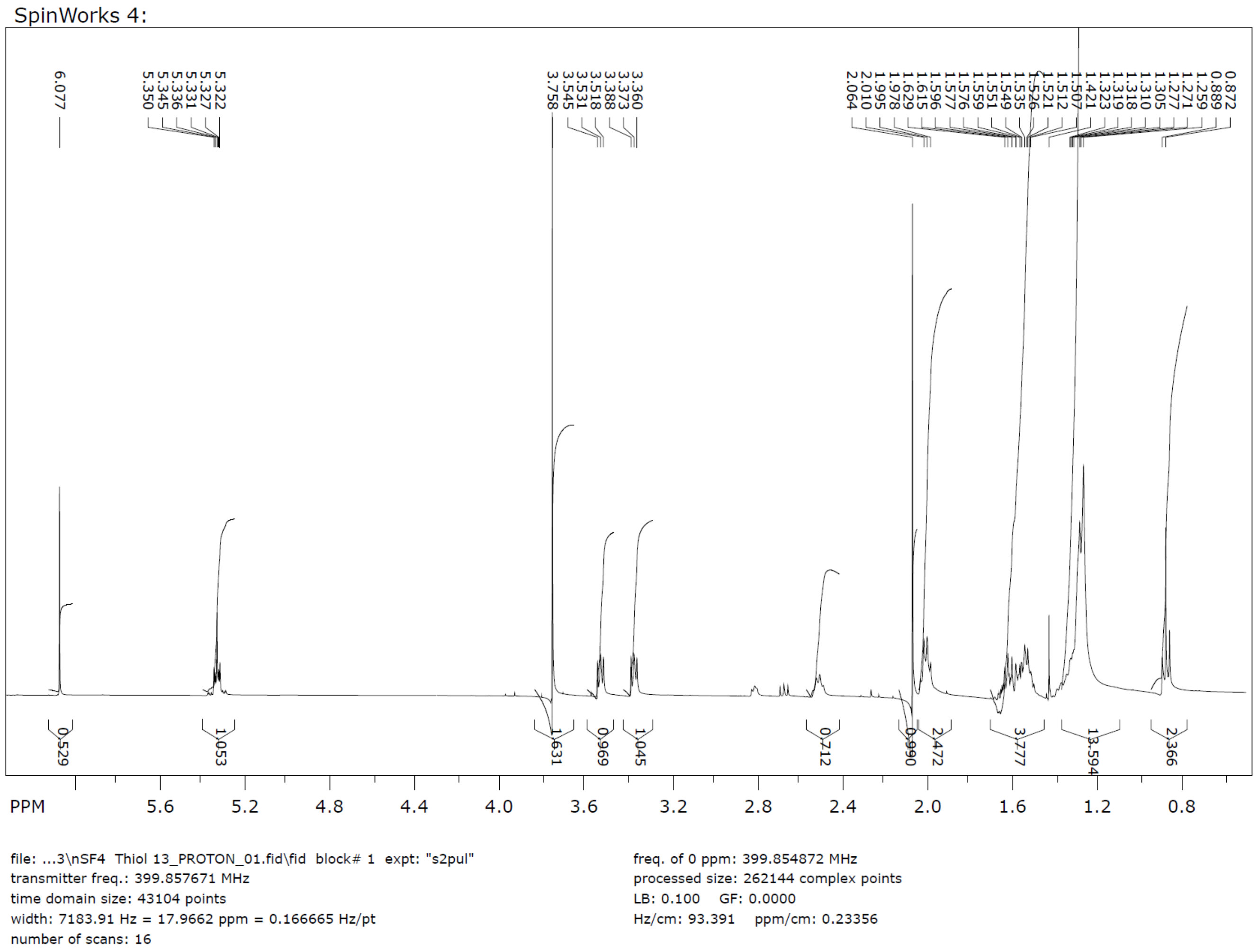


*Figure 16: 1H-NMR of compound 13.*

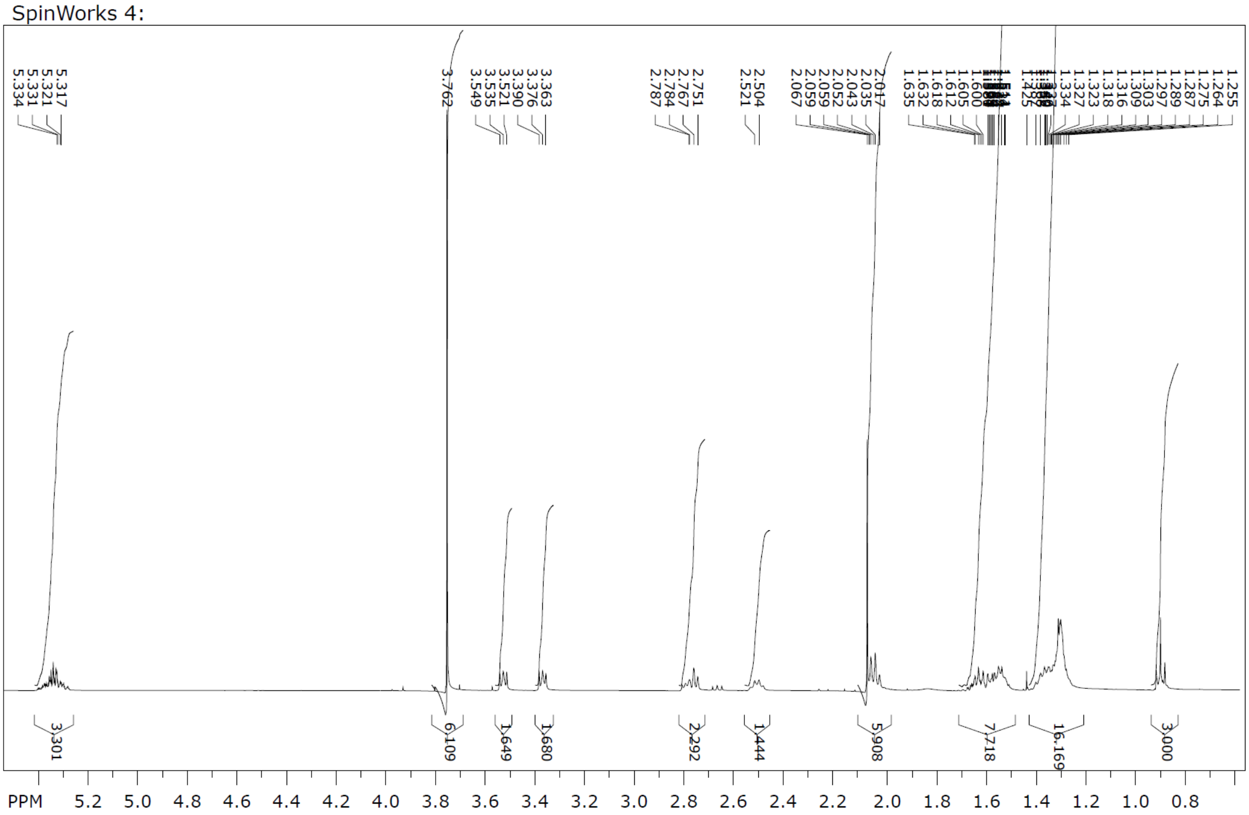

file: ...18\nSF5 Thiol 18_PROTON_01.fid\fid block# 1 expt: "s2pul"
transmitter freq.: 399.857671 MHz
time domain size: 43104 points
width: 7183.91 Hz = 17.9662 ppm = 0.166665 Hz/pt
number of scans: 16

freq. of 0 ppm: 399.854872 MHz
processed size: 262144 complex points
LB: 0.100 GF: 0.0000
Hz/cm: 79.730 ppm/cm: 0.19939

*Figure 17: 1H-NMR of compound 18.*

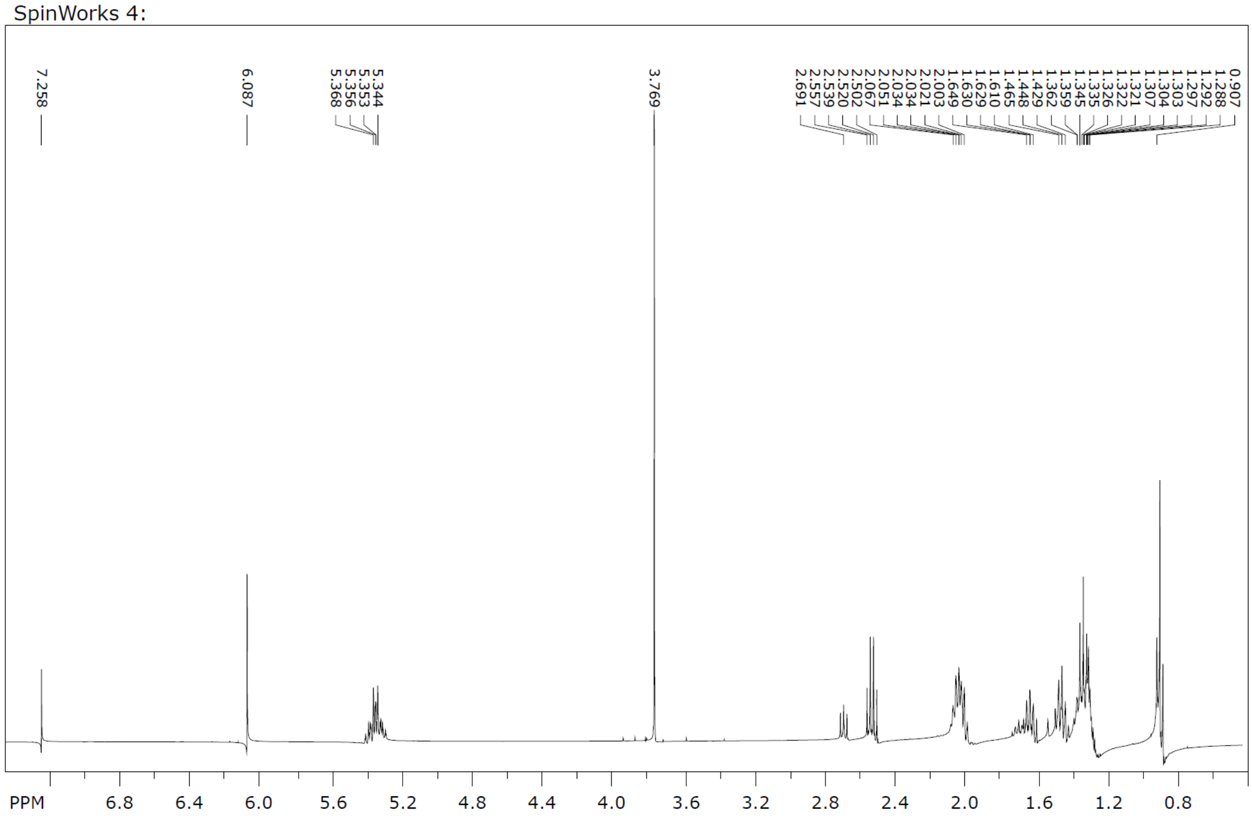

file: ...hiol 8 nSF1 -FC2_PROTON_01.fid\fid block# 1 expt: "s2pul"
transmitter freq.: 399.857671 MHz
time domain size: 43104 points
width: 7183.91 Hz = 17.9662 ppm = 0.166665 Hz/pt
number of scans: 16

freq. of 0 ppm: 399.854872 MHz
processed size: 262144 complex points
LB: 0.100 GF: 0.0000
Hz/cm: 112.705 ppm/cm: 0.28186

*Figure 18: 1H-NMR of compound 8.*

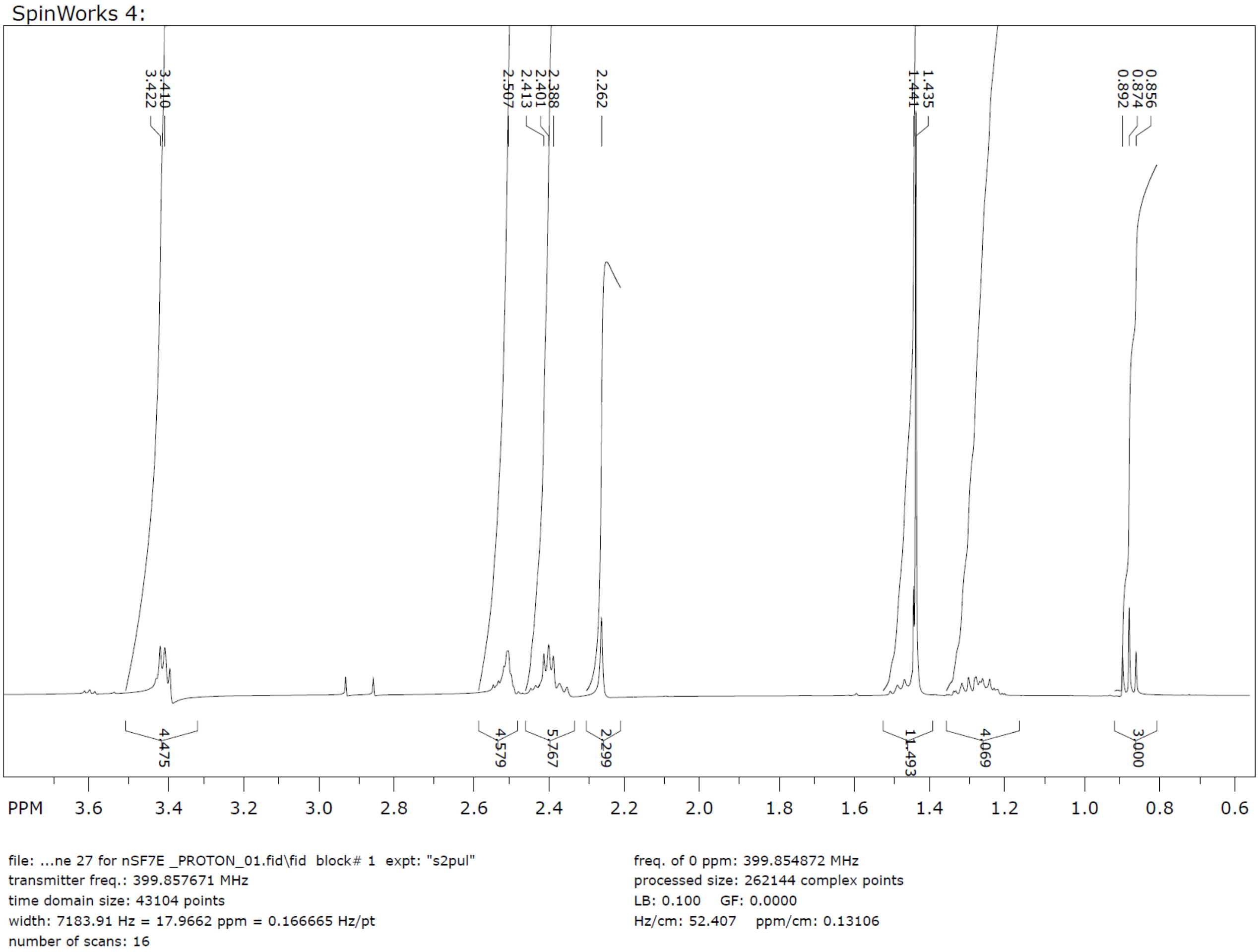


*Figure 19: 1H-NMR of compound 27.*

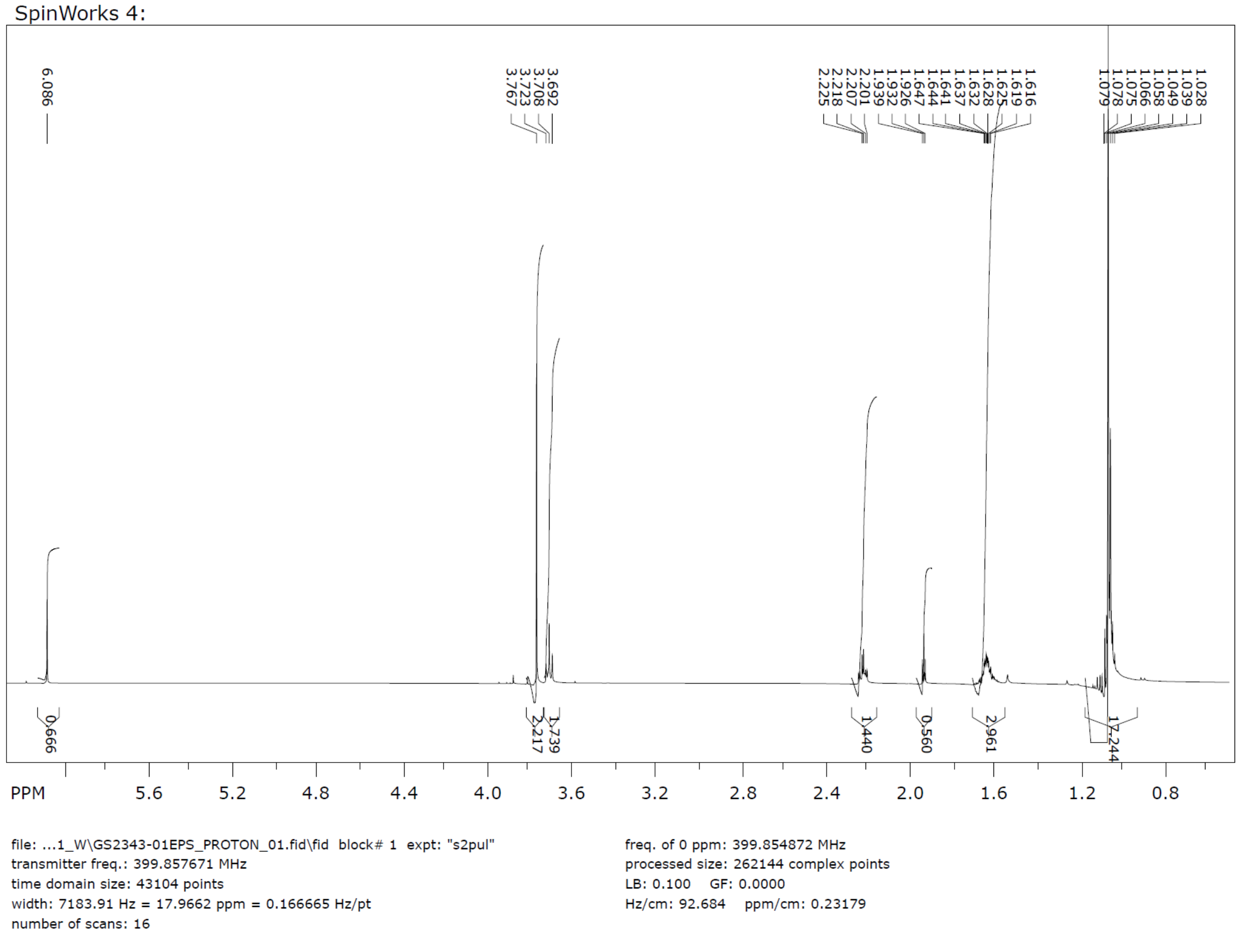


*Figure 21: 1H-NMR of compound 34.*

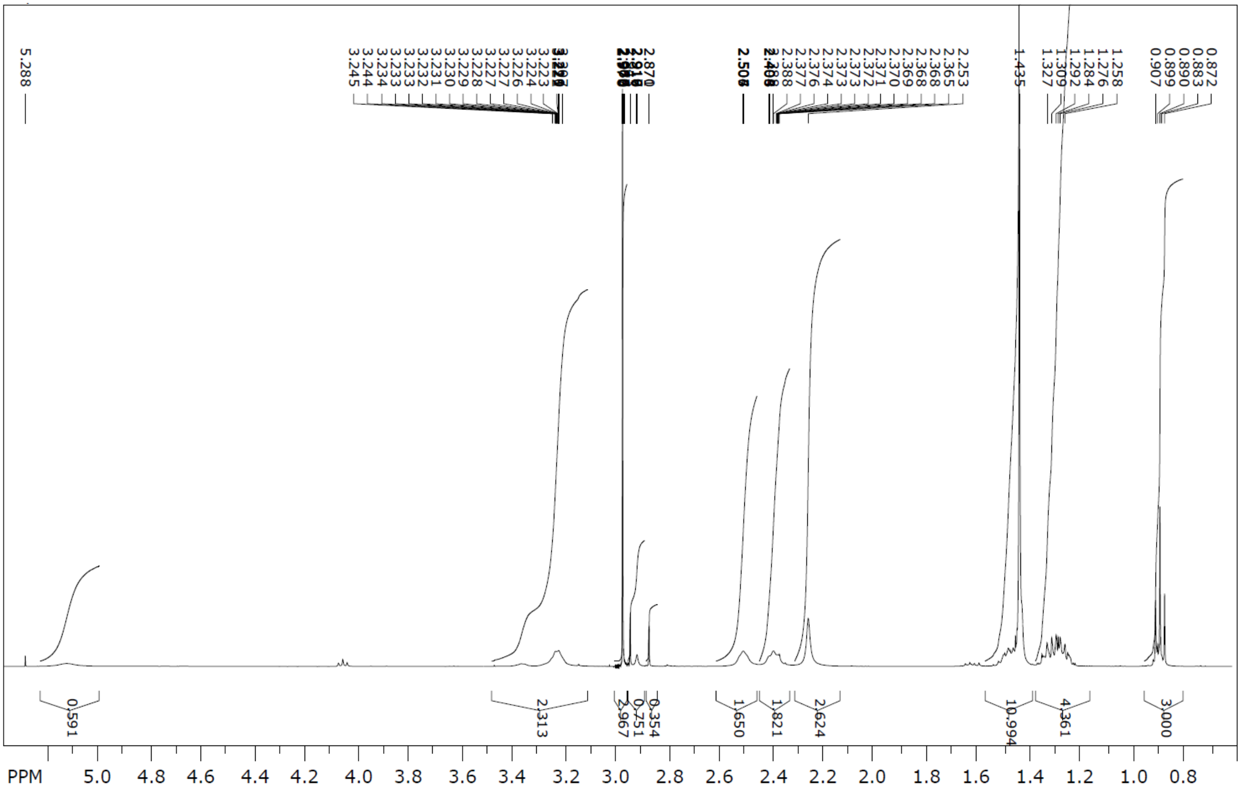


file: ... 27\Boc-Amine 27_PROTON_01.fid\fid block# 1 expt: "s2pul"
transmitter freq.: 399.857671 MHz
time domain size: 43104 points
width: 7183.91 Hz = 17.9662 ppm = 0.166665 Hz/pt
number of scans: 16

freq. of 0 ppm: 399.854872 MHz
processed size: 262144 complex points
LB: 0.100 GF: 0.0000
Hz/cm: 76.197 ppm/cm: 0.19056

*Figure 22: 1H-NMR of compound 27.*

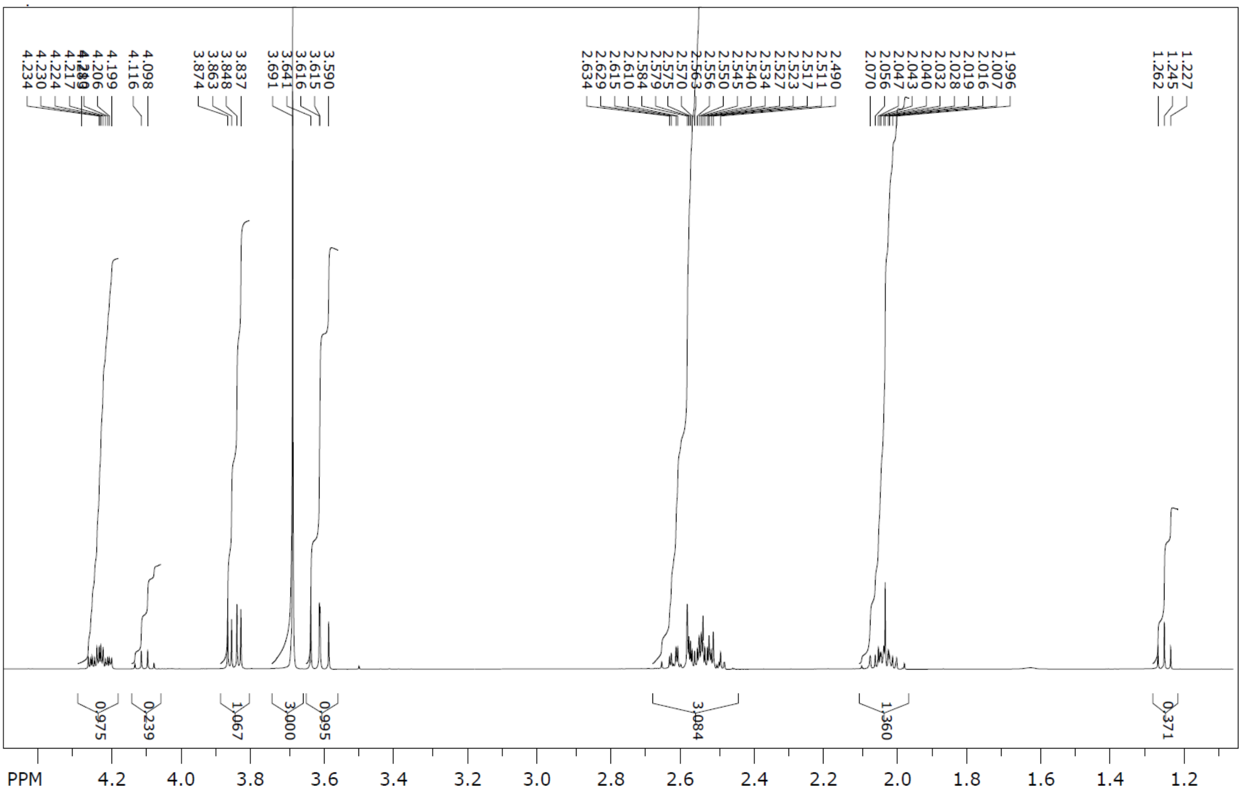


file: ... 25\dibromide 25_PROTON_01.fid\fid block# 1 expt: "s2pul"
transmitter freq.: 399.857671 MHz
time domain size: 43104 points
width: 7183.91 Hz = 17.9662 ppm = 0.166665 Hz/pt
number of scans: 16

freq. of 0 ppm: 399.854872 MHz
processed size: 262144 complex points
LB: 0.100 GF: 0.0000
Hz/cm: 55.234 ppm/cm: 0.13813

*Figure 23: 1H-NMR of compound 25.*

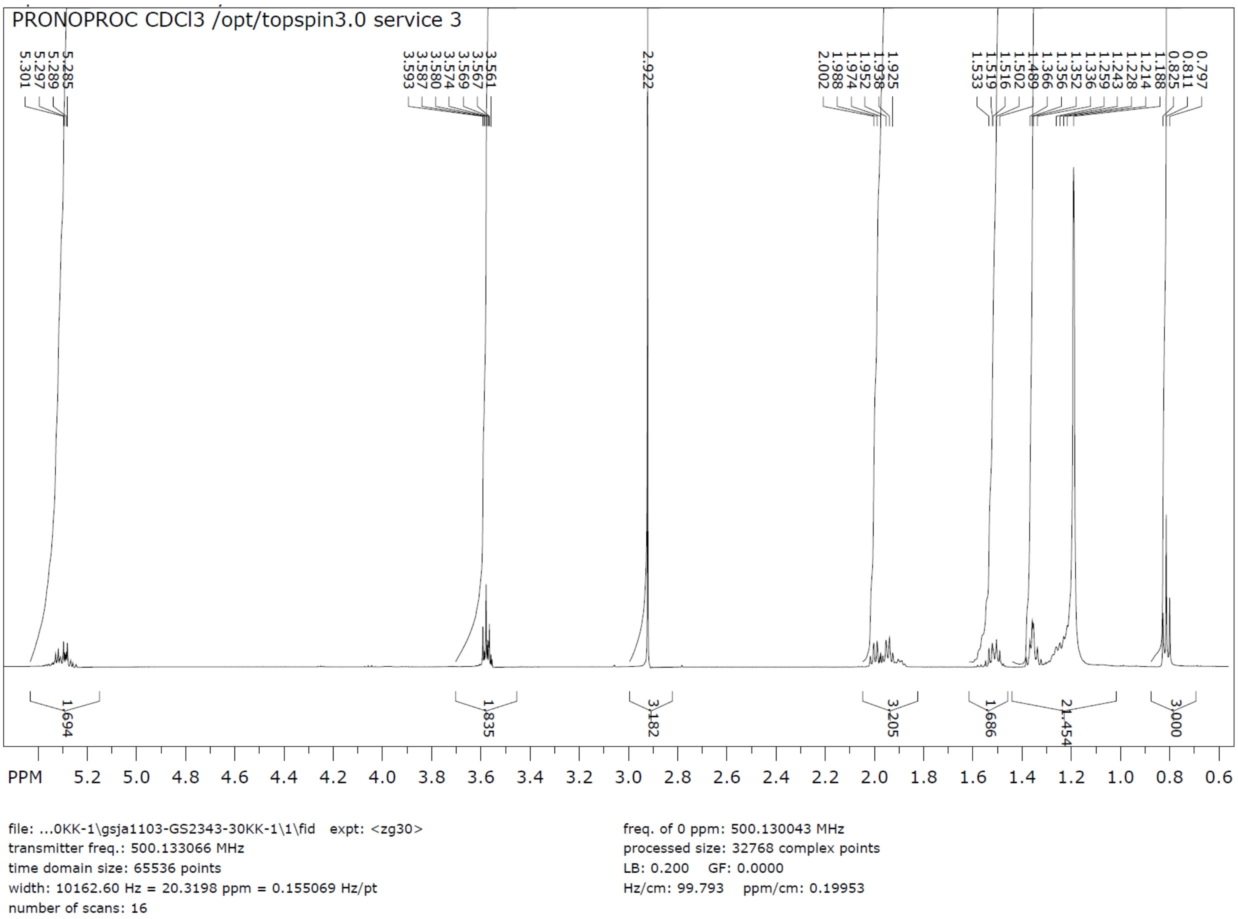


*Figure 24: 1H-NMR of compound 37.*

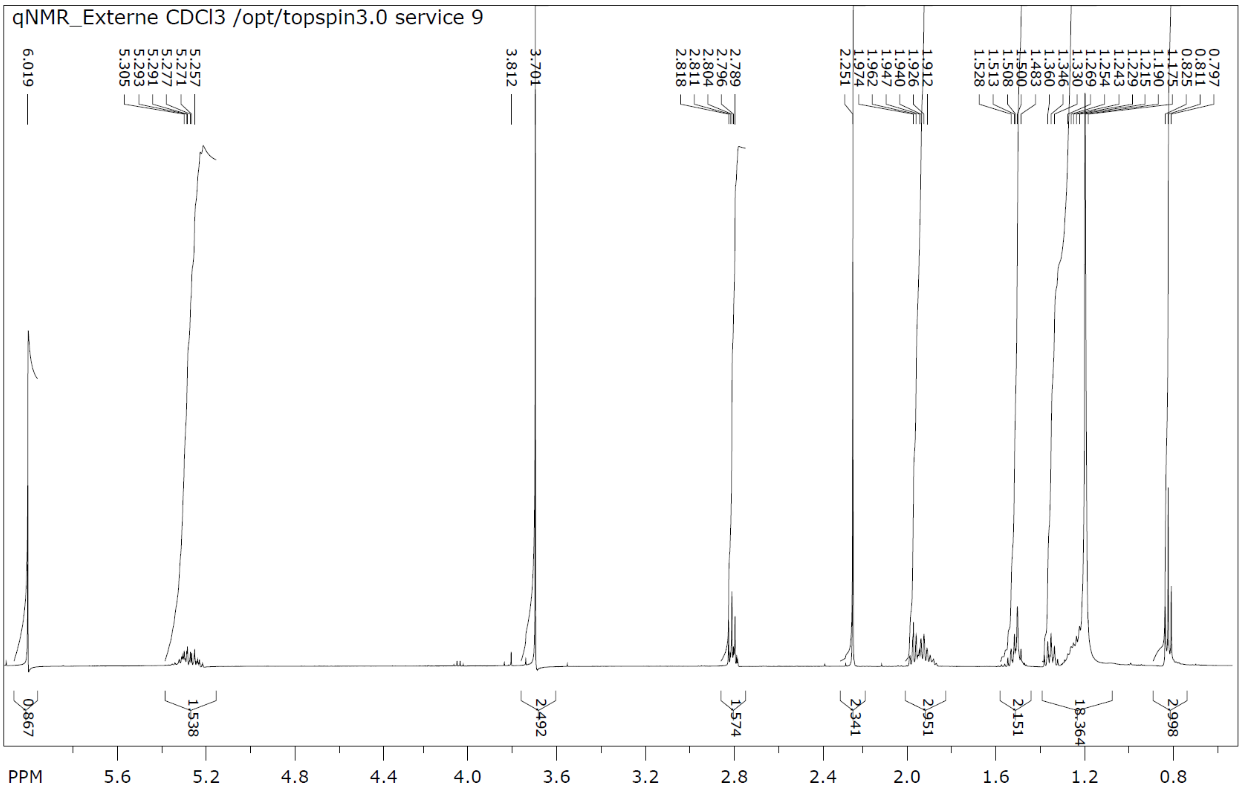


file: ...Anton\1H NMR 10. compound 38\1\fid expt: <zg30>
transmitter freq.: 500.133066 MHz
time domain size: 65536 points
width: 10162.60 Hz = 20.3198 ppm = 0.155069 Hz/pt
number of scans: 16

freq. of 0 ppm: 500.130046 MHz
processed size: 32768 complex points
LB: 0.200 GF: 0.0000
Hz/cm: 112.288 ppm/cm: 0.22452

*Figure 25: 1H-NMR of compound 38.*

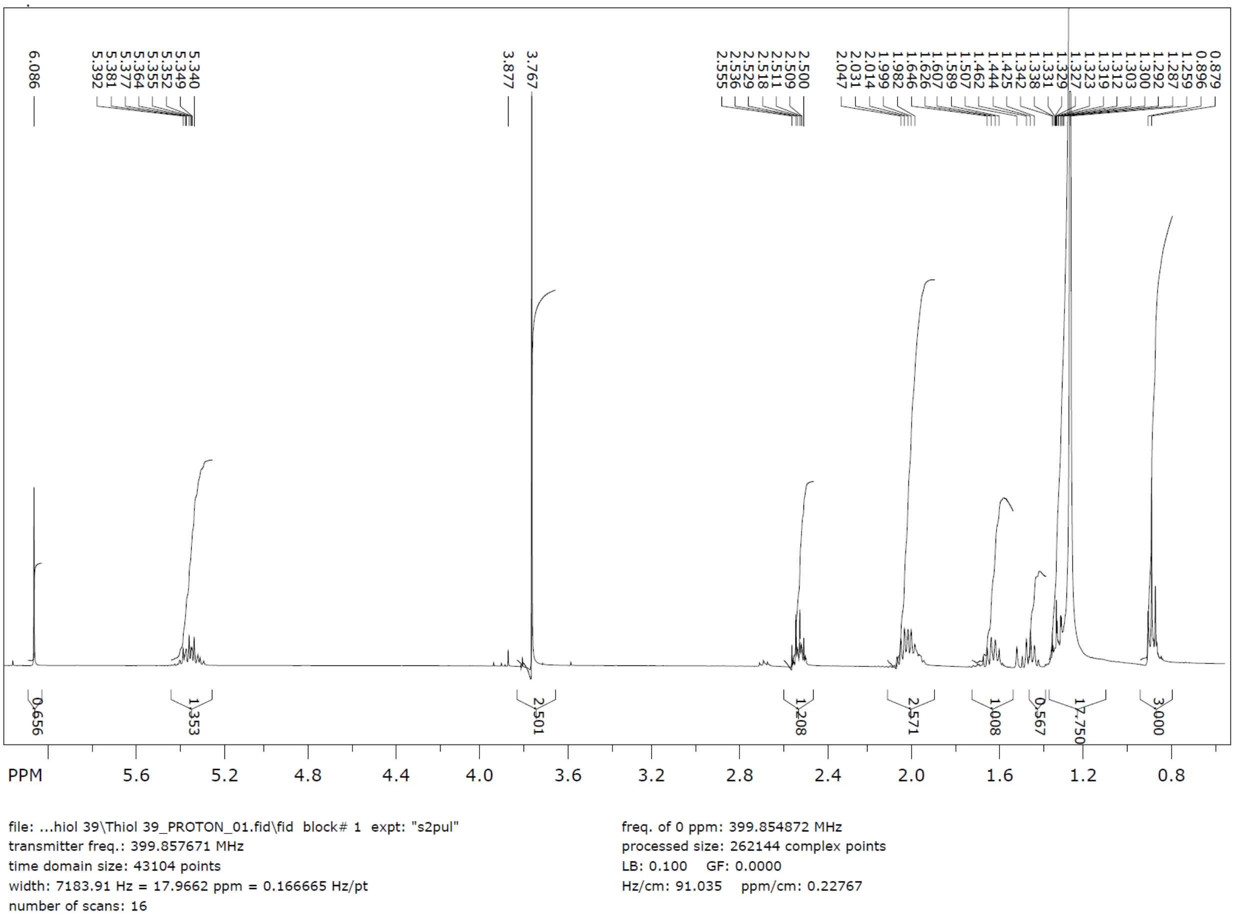


*Figure 26: 1H-NMR of compound 39.*

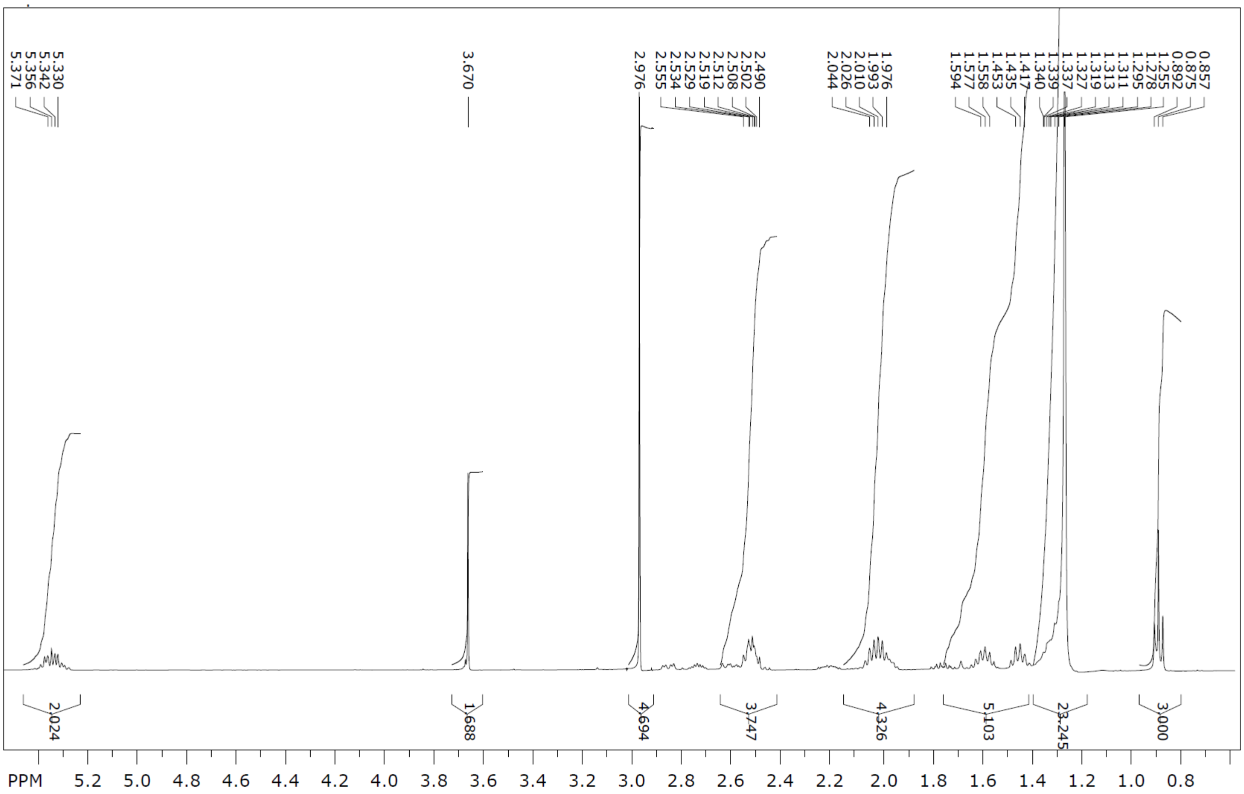

file: ...ster 40\ester 40_PROTON_01.fid\fid block# 1 expt: "s2pul"
transmitter freq.: 399.857671 MHz
time domain size: 43104 points
width: 7183.91 Hz = 17.9662 ppm = 0.166665 Hz/pt
number of scans: 16

freq. of 0 ppm: 399.854872 MHz
processed size: 262144 complex points
LB: 0.100 GF: 0.0000
Hz/cm: 79.730 ppm/cm: 0.19939

*Figure 27: 1H-NMR of compound 40.*

### High Resolution Mass Spectrum Analysis

found m/z = 737.6408, calc $[M+H]^+$ = 737.6411.

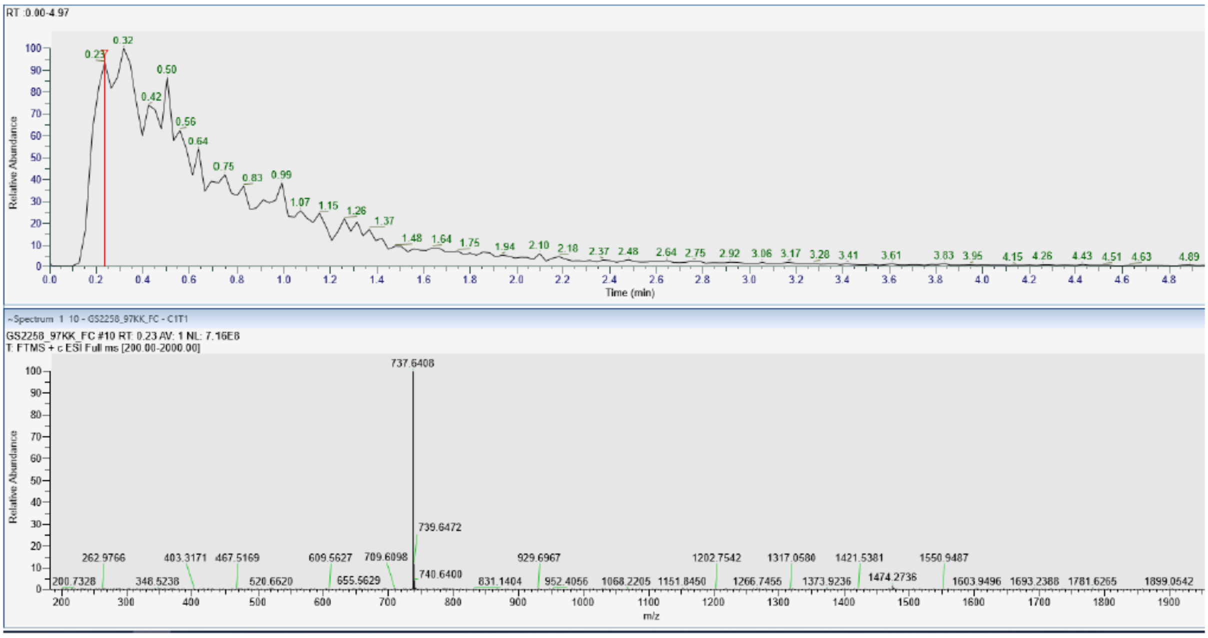


*Figure 28: High-resolution MS of compound A1C18_D5*

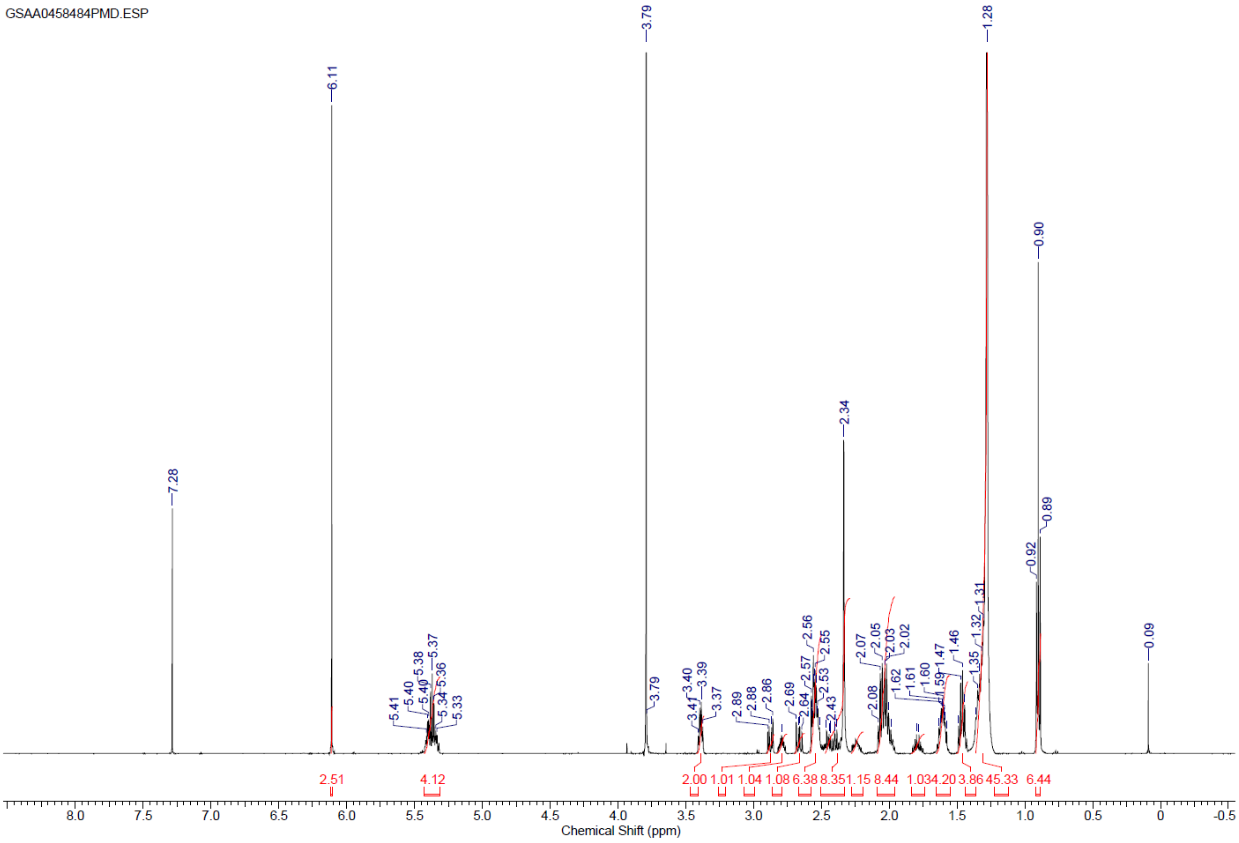


*Figure 29: 1H-NMR of A1C18_D5*

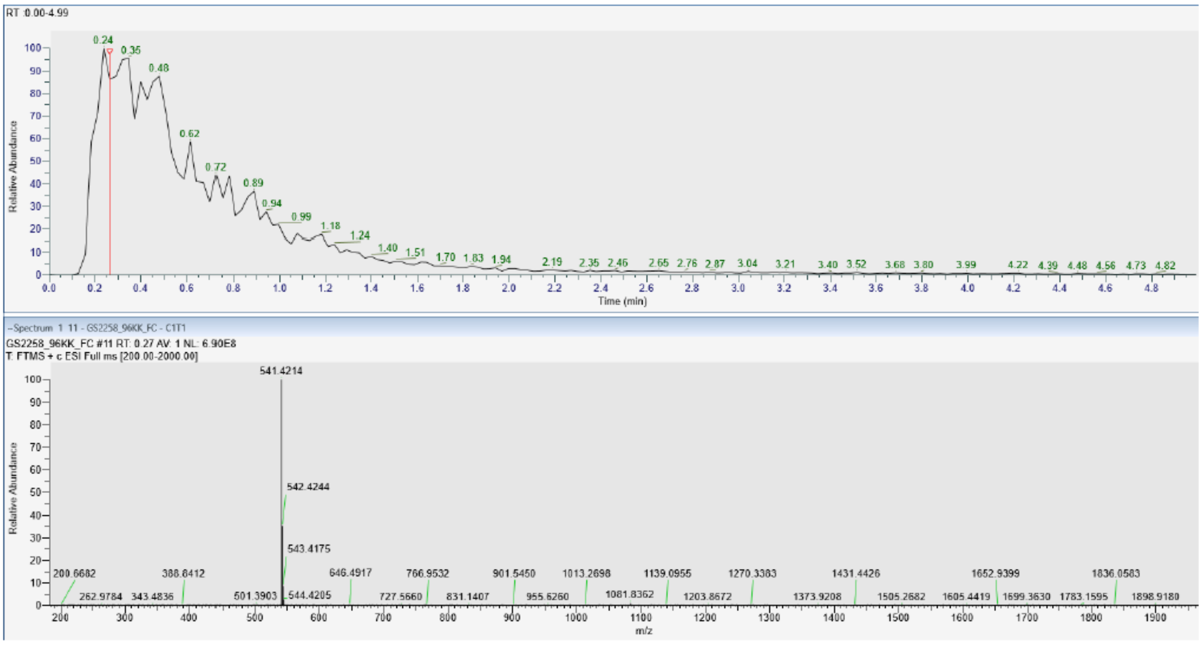


*Figure 30: High resolution MS of compound A1C11_D5 (calc. [M+H]+ = 541,4120)*

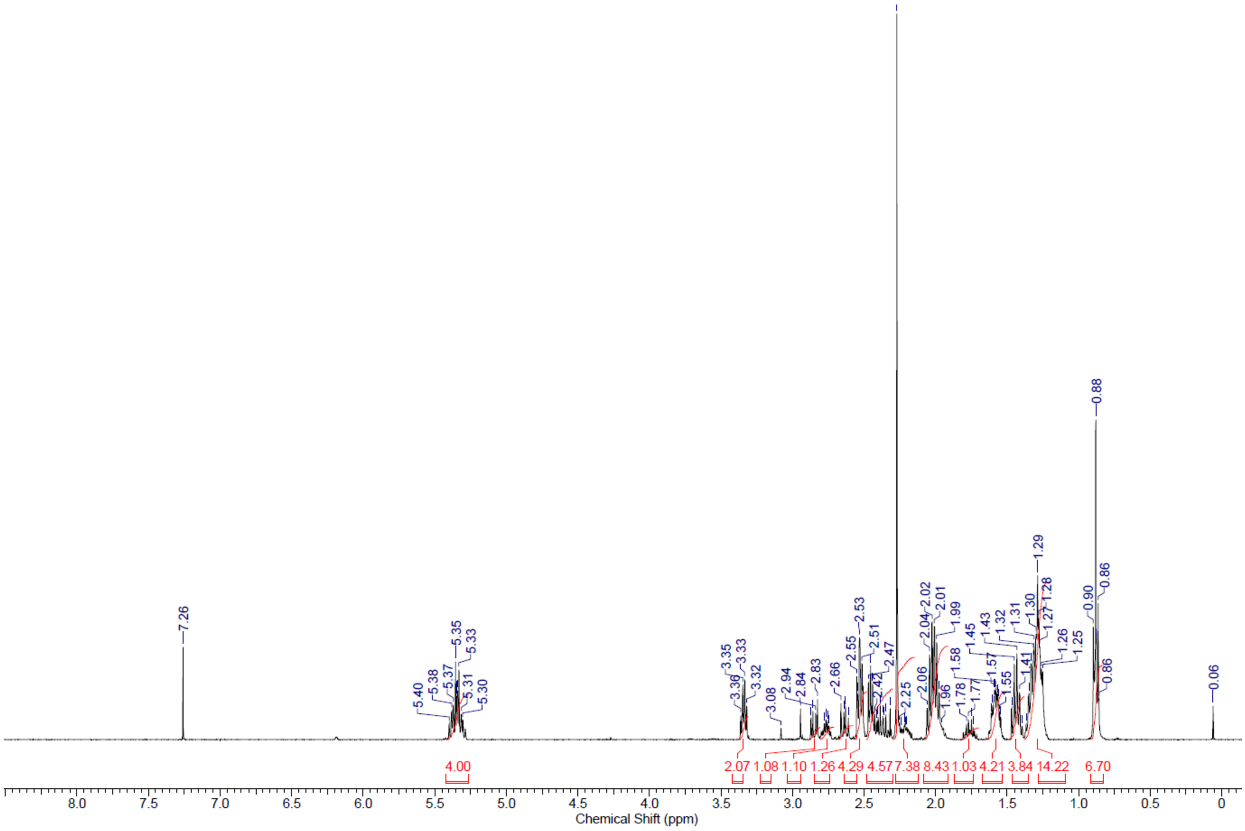


*Figure 31: 1H-NMR of compound A1C11_D5*

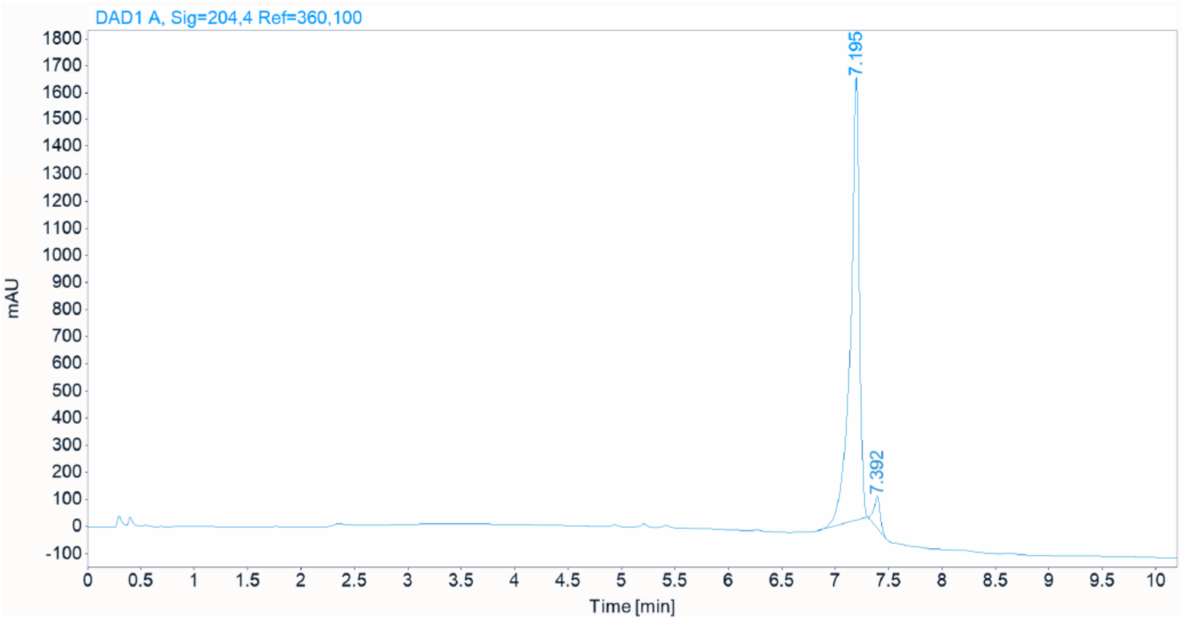


*Figure 32: HPLC trace of A1C18_D9_D12*

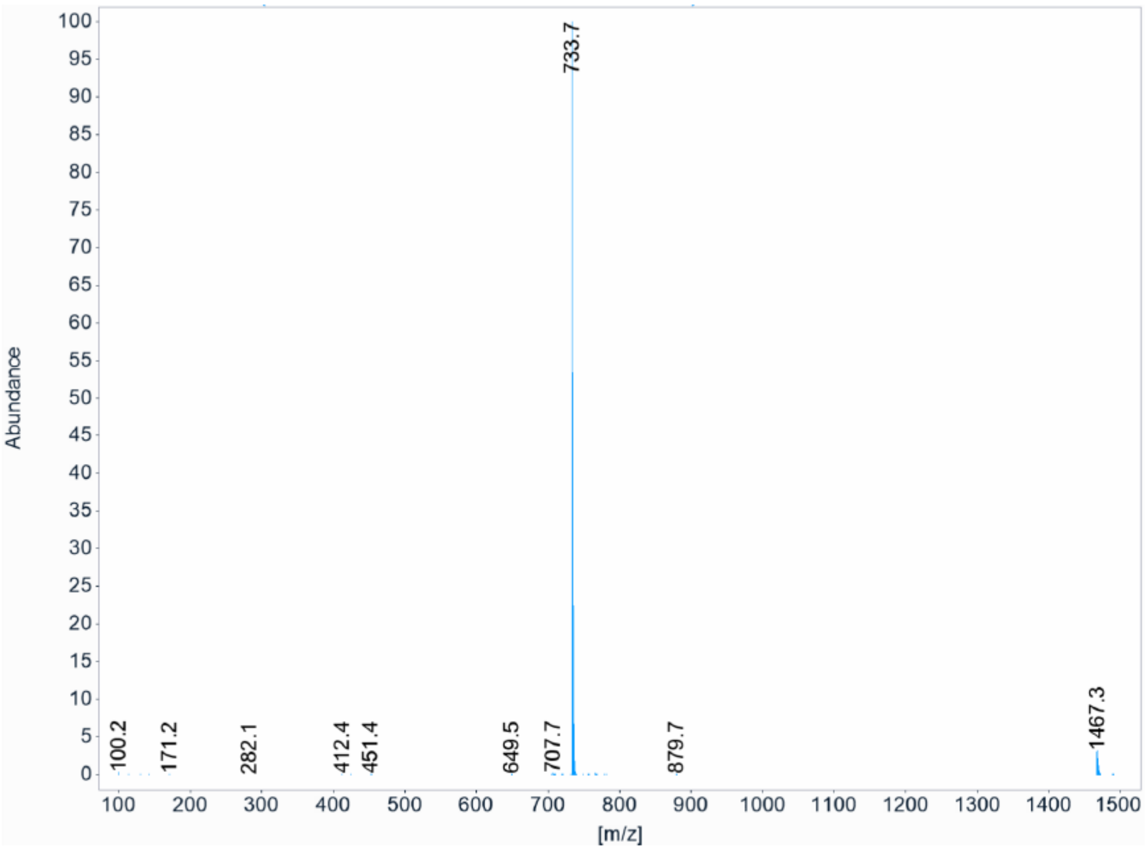


*Figure 33: MS spectrum of A1C18_D9_D12*

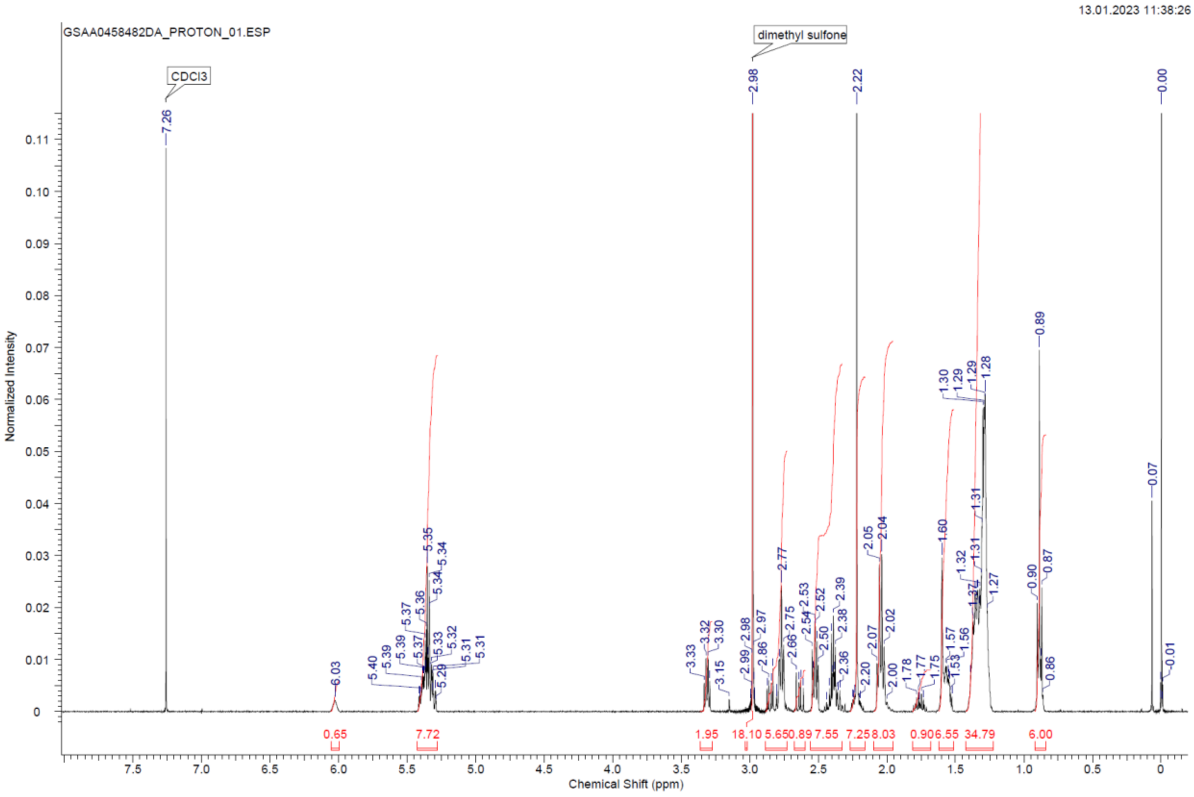


*Figure 34: 1H-NMR of compound A1C18_D9_D12*

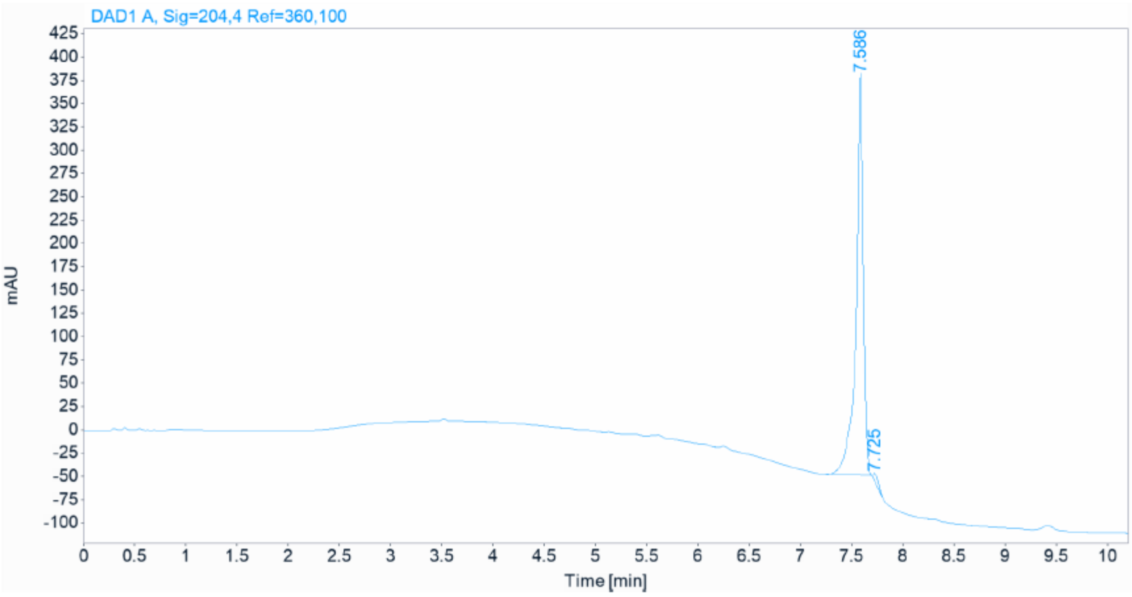


*Figure 35: HPLC trace of A1C18_D9*

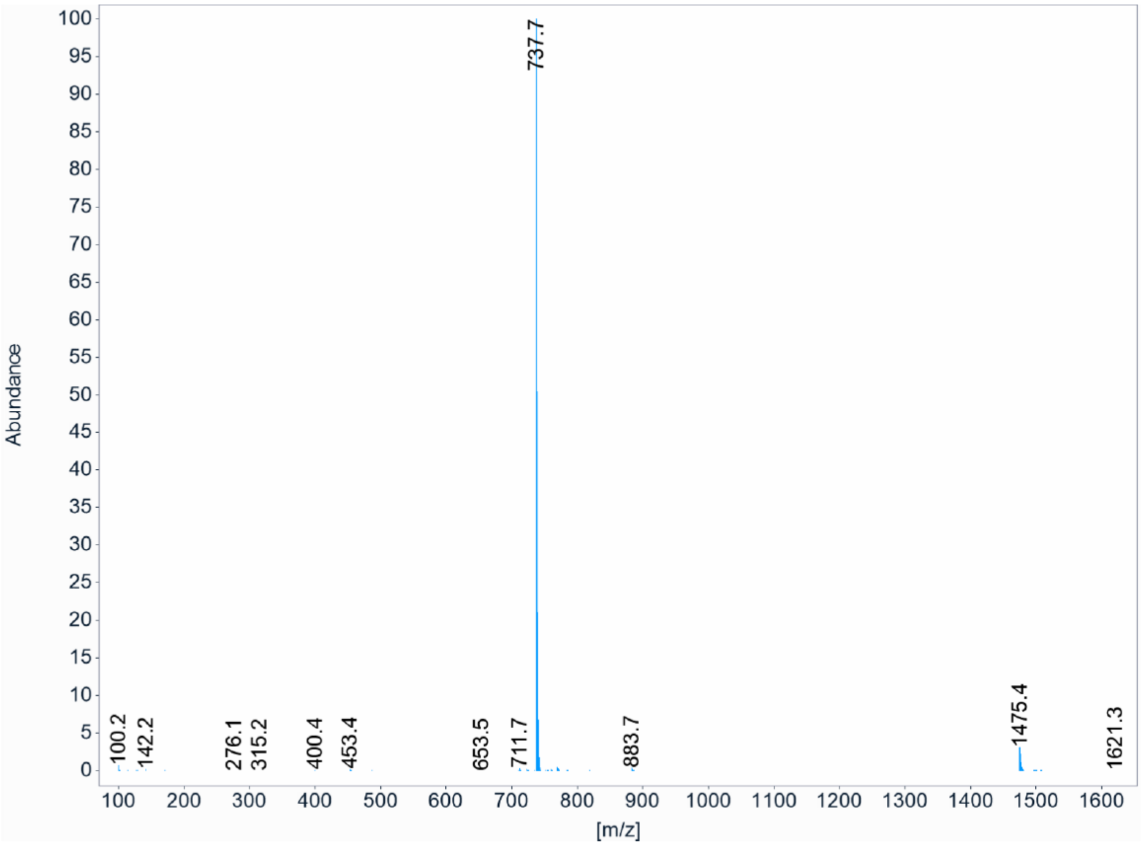


*Figure 36: ESI-MS of A1C18_D9*

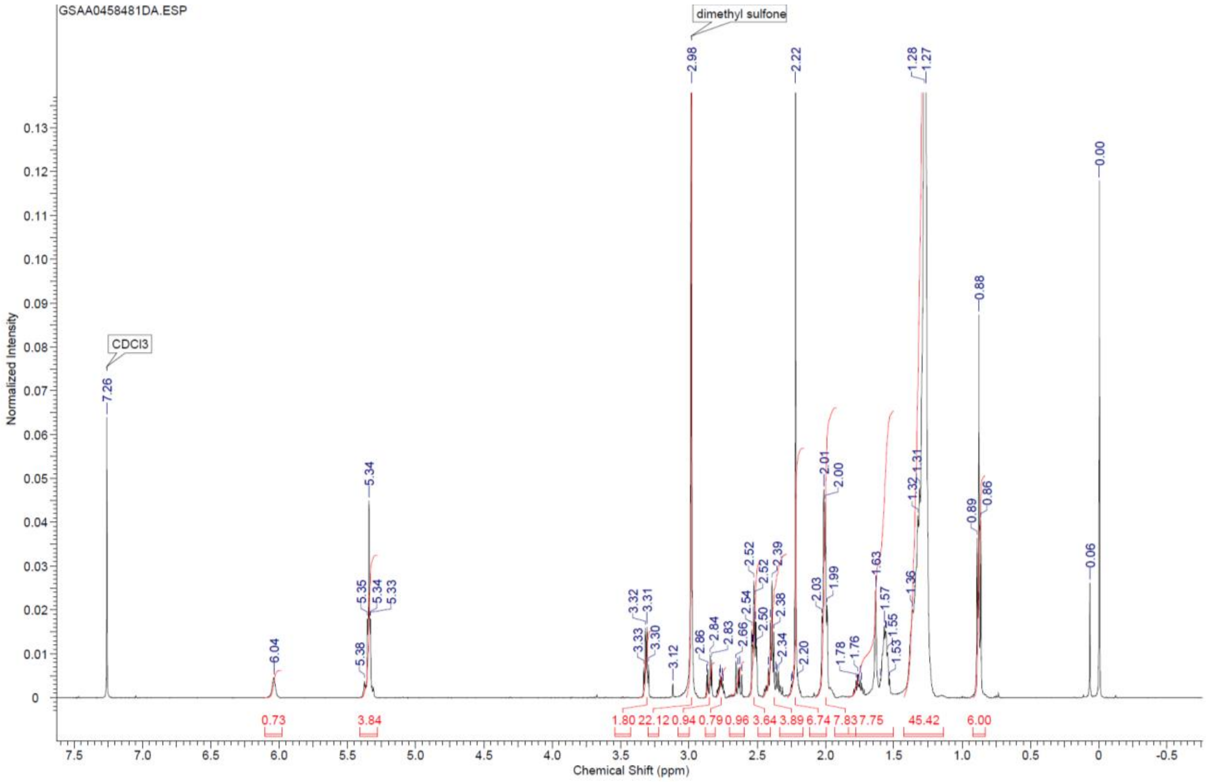


*Figure 37: 1H-NMR of compound A1C18_D9*

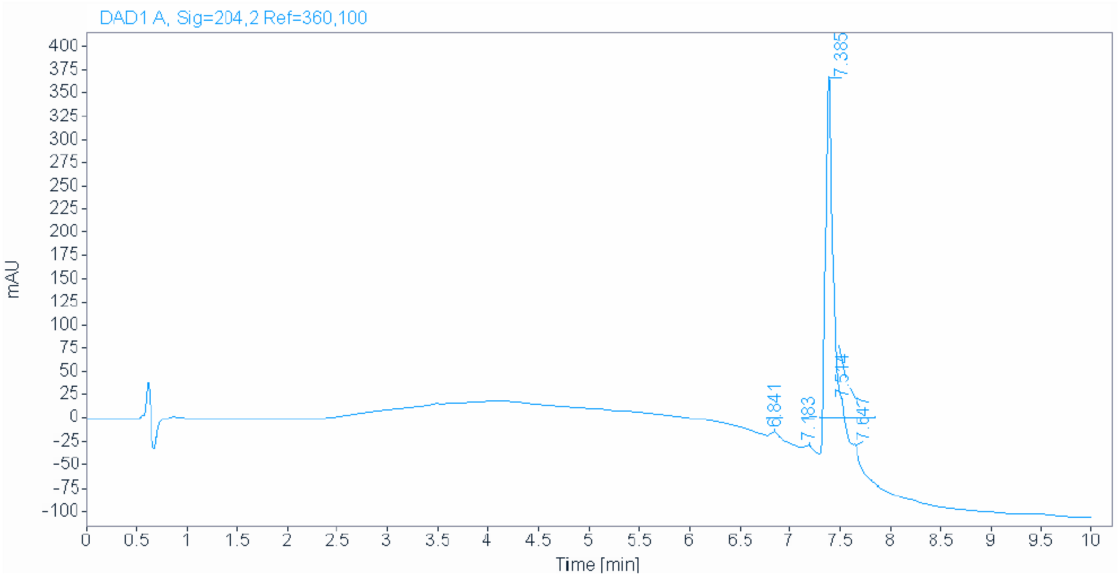


*Figure 38: HPLC trace of compound A3C18_D5*

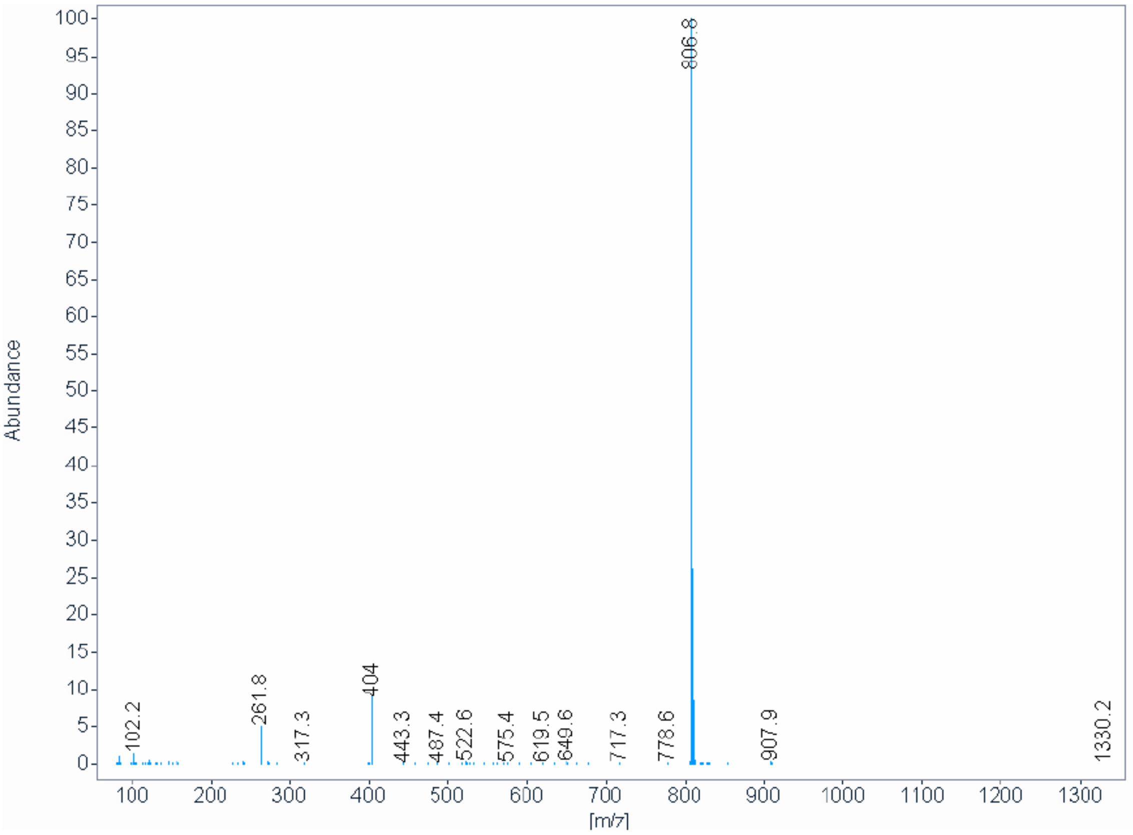


*Figure 39: ESI-MS of A3C18_D5*

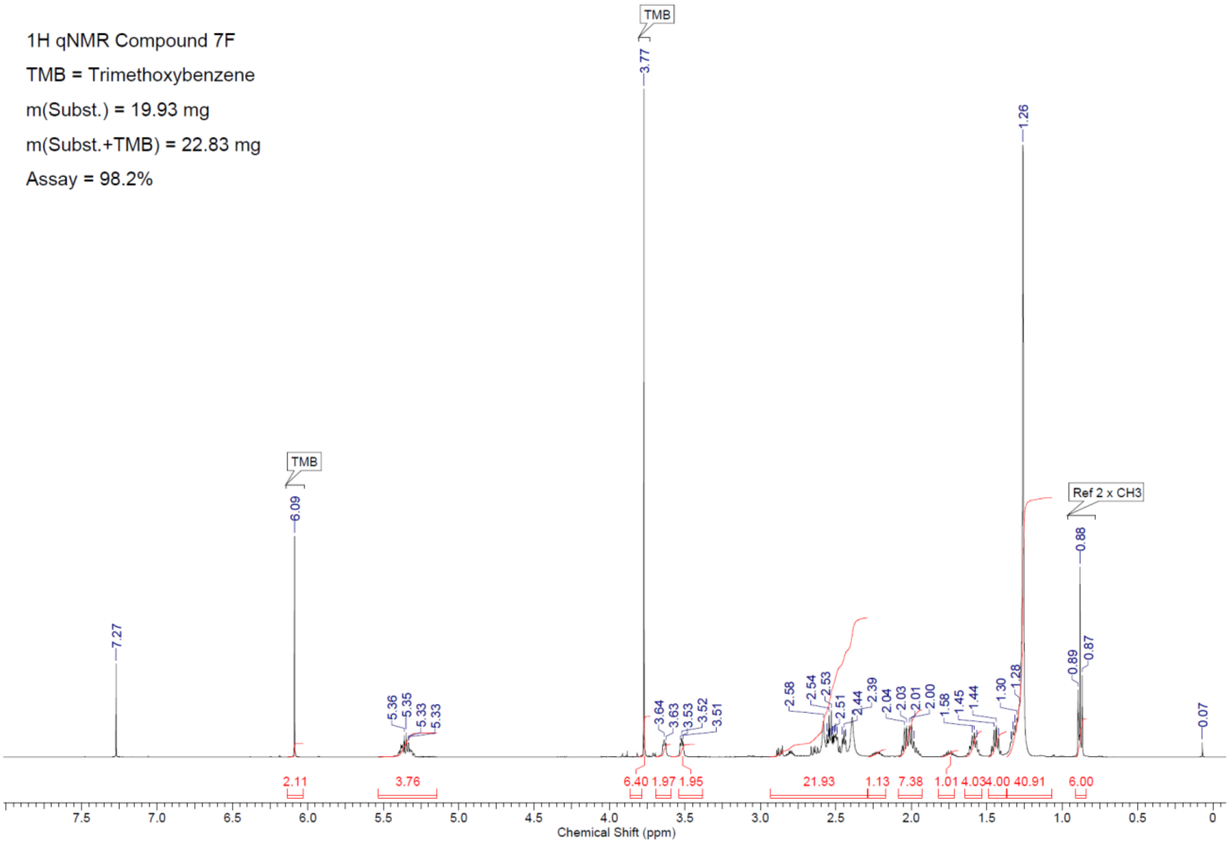


*Figure 40: 1H-NMR of compound A3C18_D5*

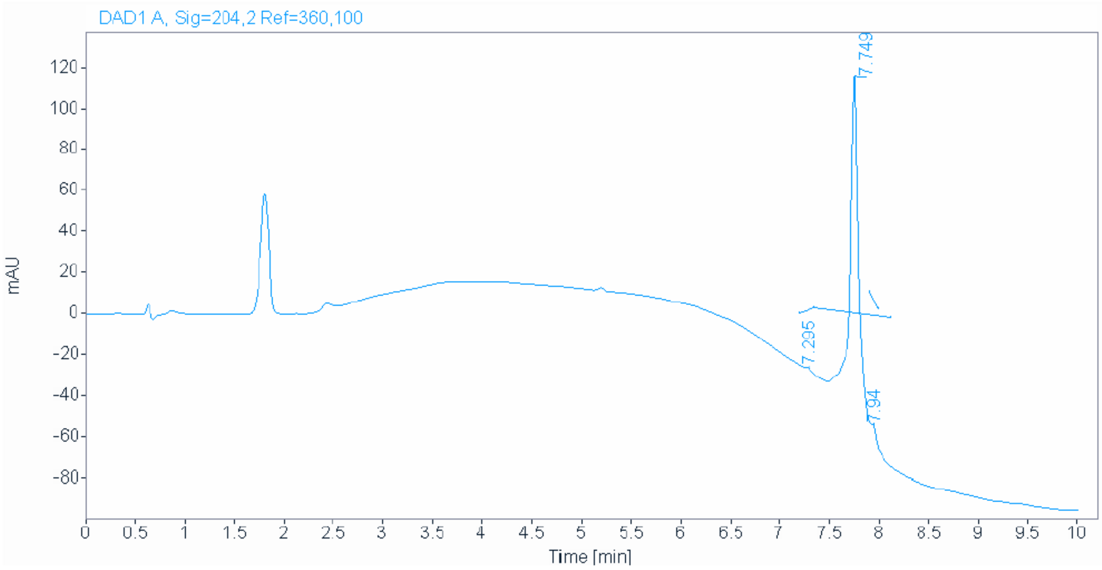


*Figure 41: HPLC trace of compound A2C18_D5*

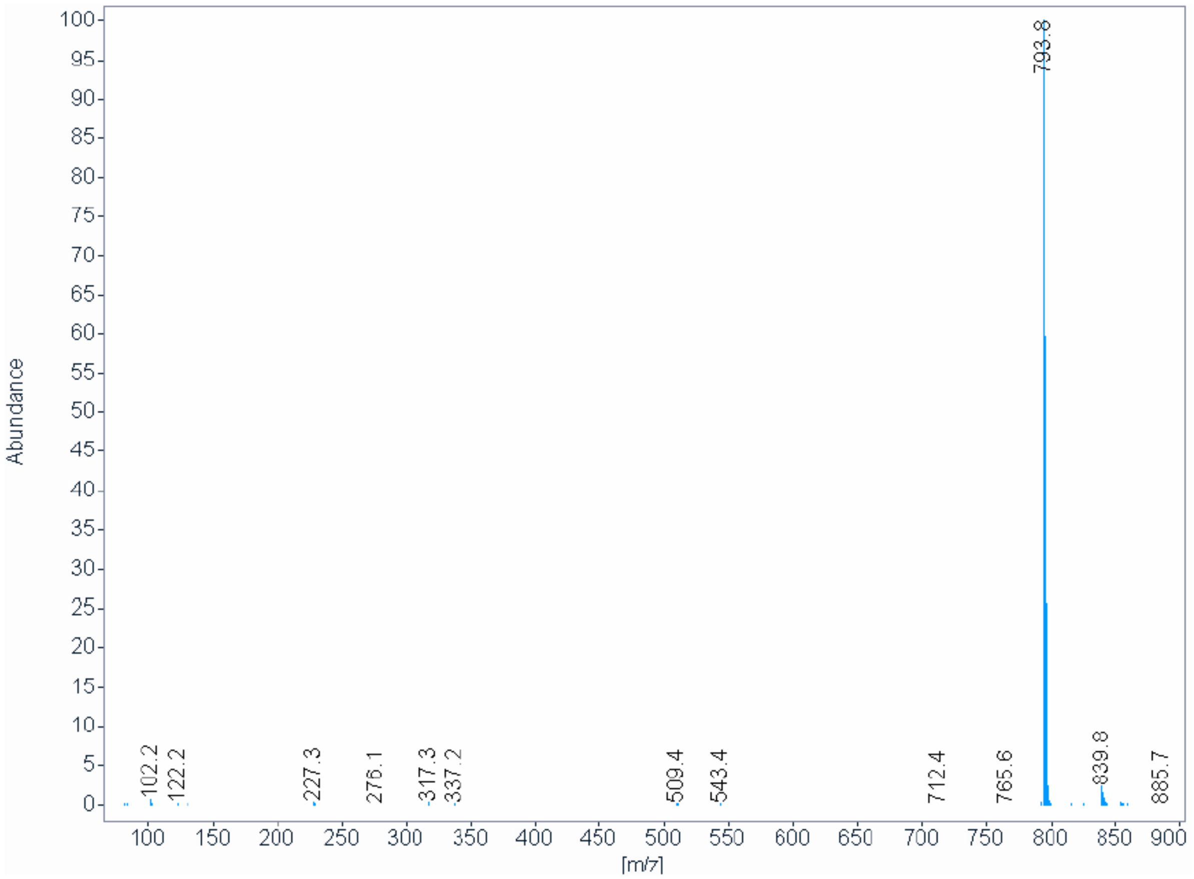


*Figure 42: ESI-MS of compound A2C18_D5*

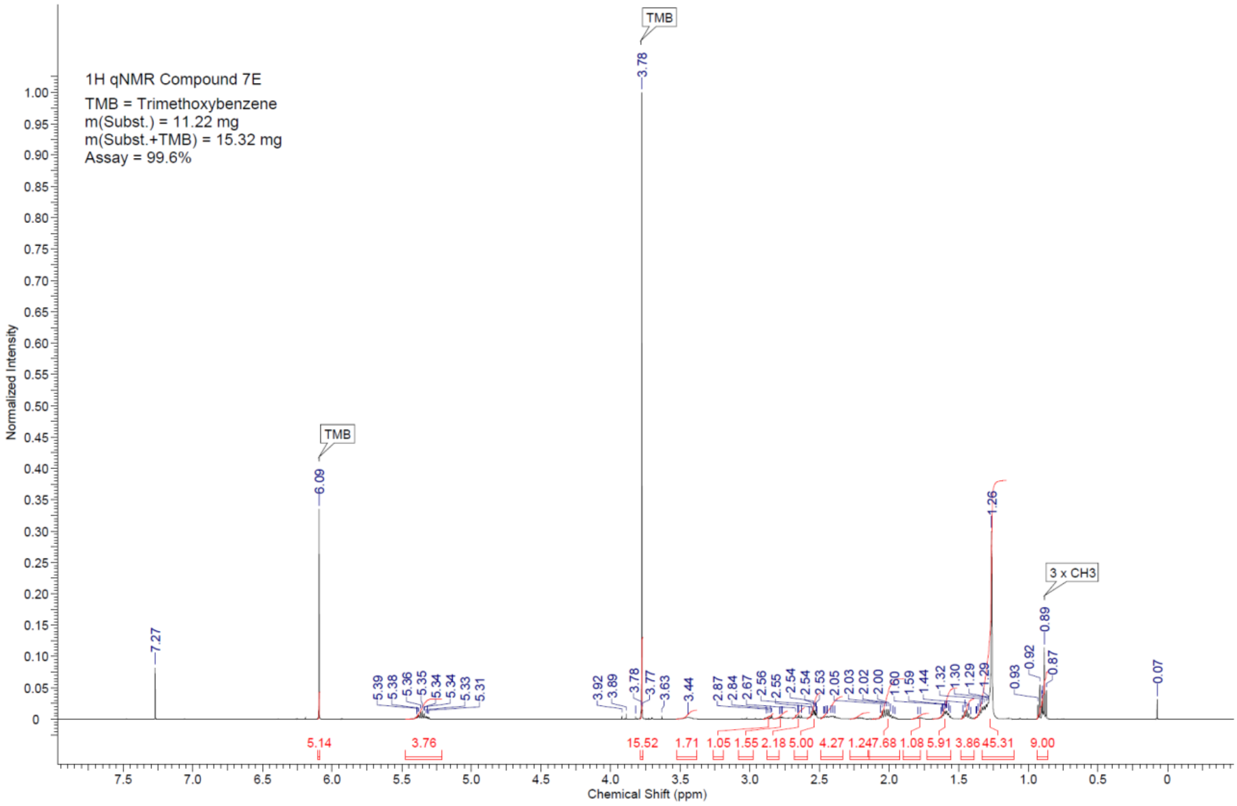


*Figure 43: 1H-NMR of compound A2C18_D5*

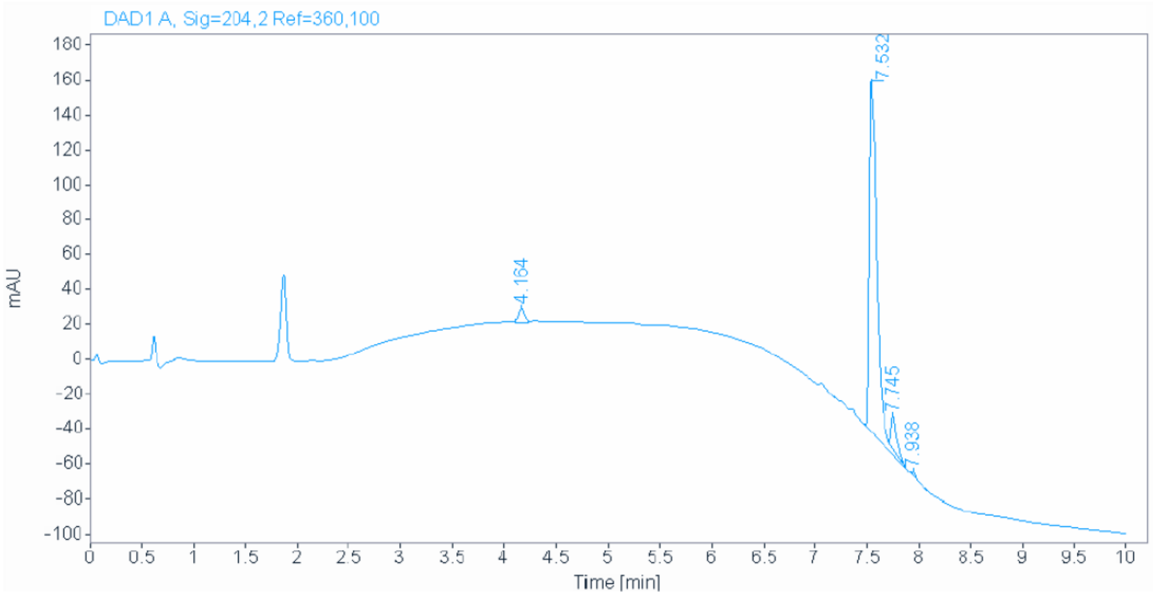


*Figure 44: HPLC trace of compound A4C18_D5*

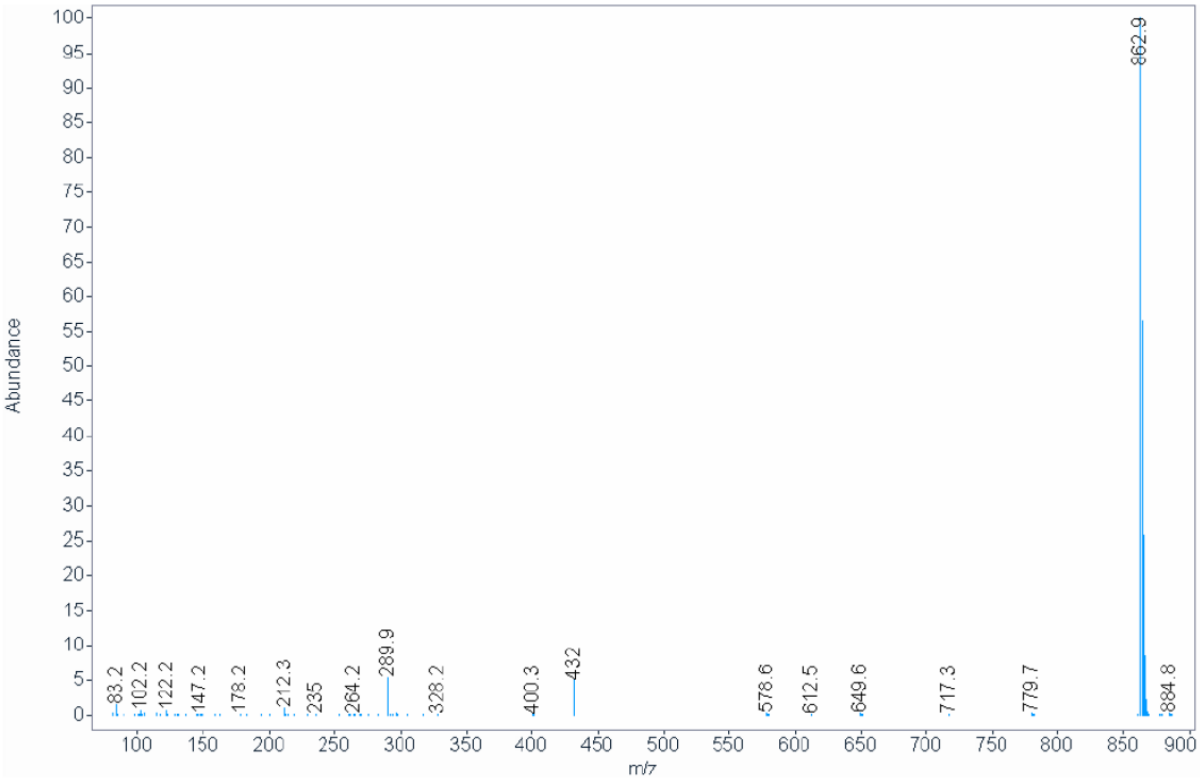


*Figure 45: ESI-MS of compound A4C18_D5*

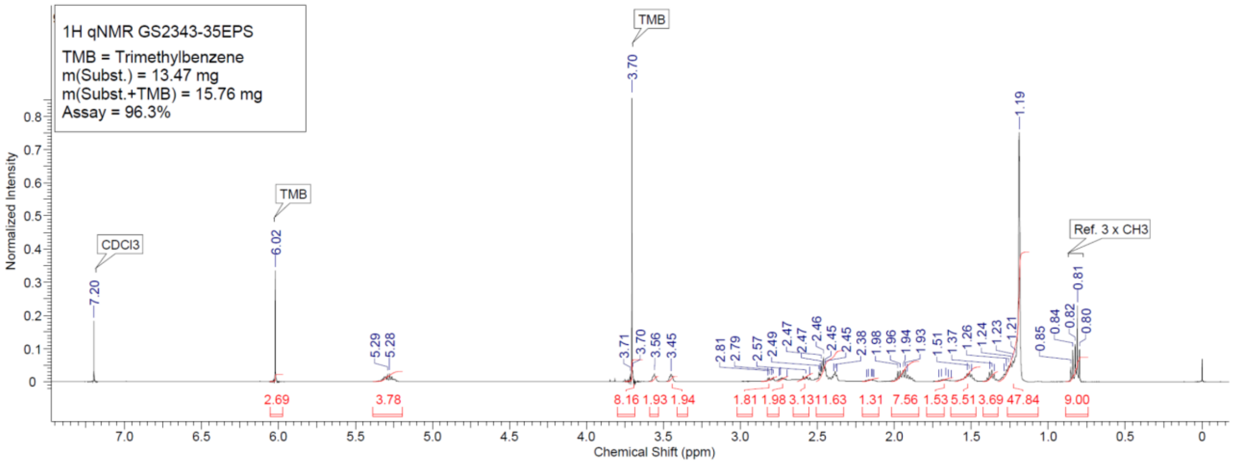


*Figure 46: 1H-NMR of compound A4C18_D5*

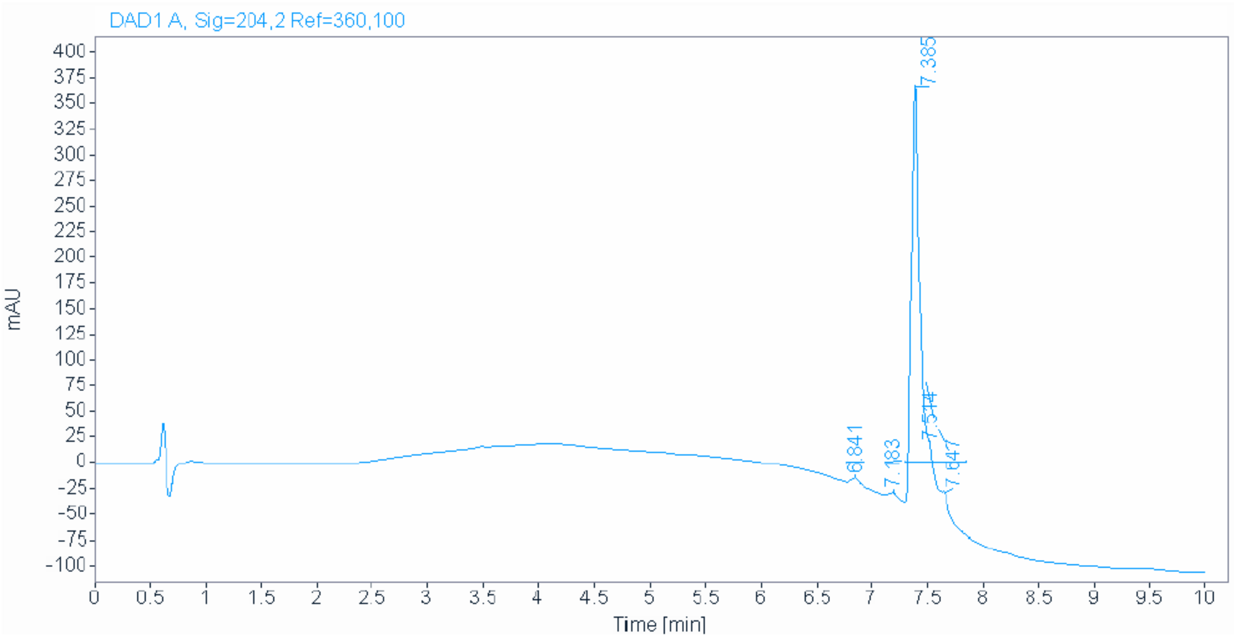


*Figure 47: HPLC trace of compound A3C18_D5*

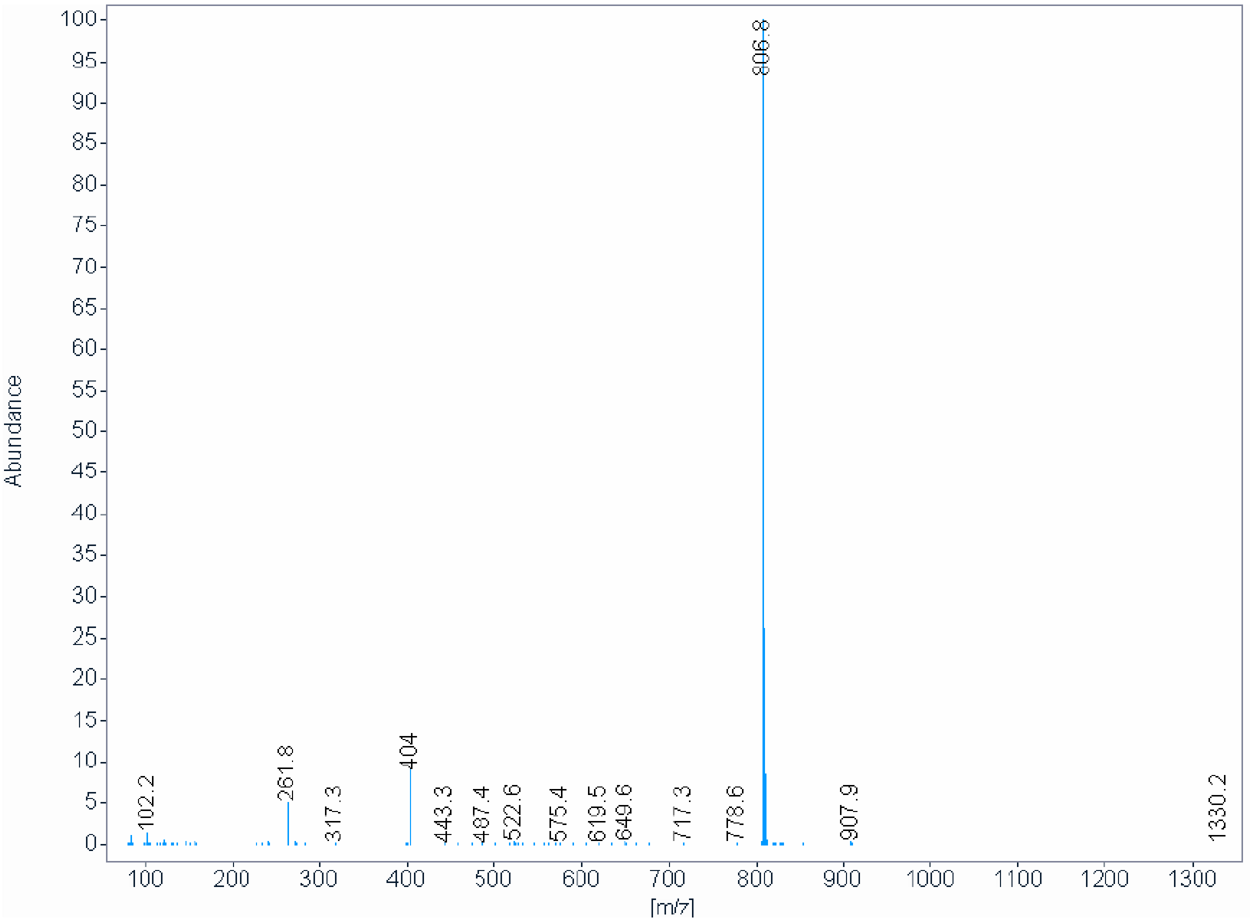


*Figure 48: ESI-MS of compound A3C18_D5*

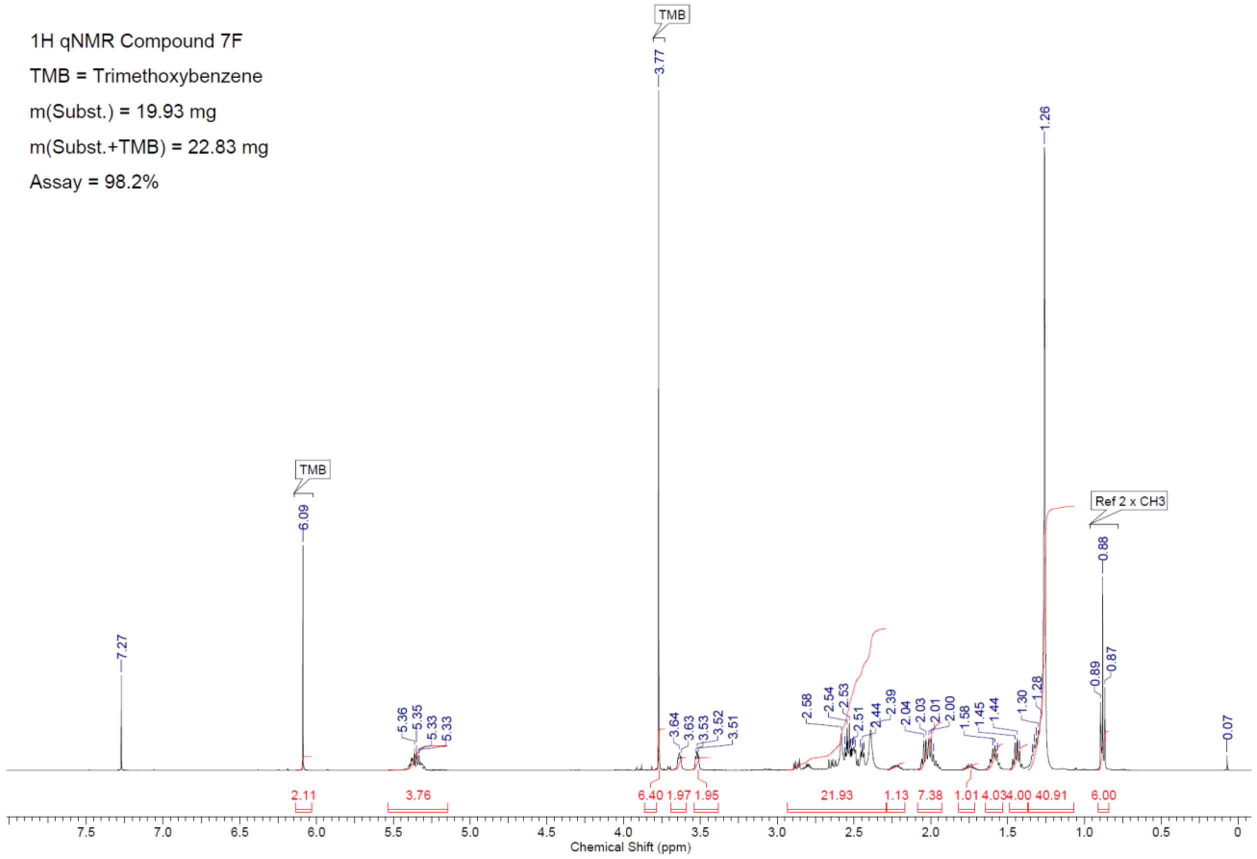


*Figure 49: 1H-NMR of compound A3C18_D5*

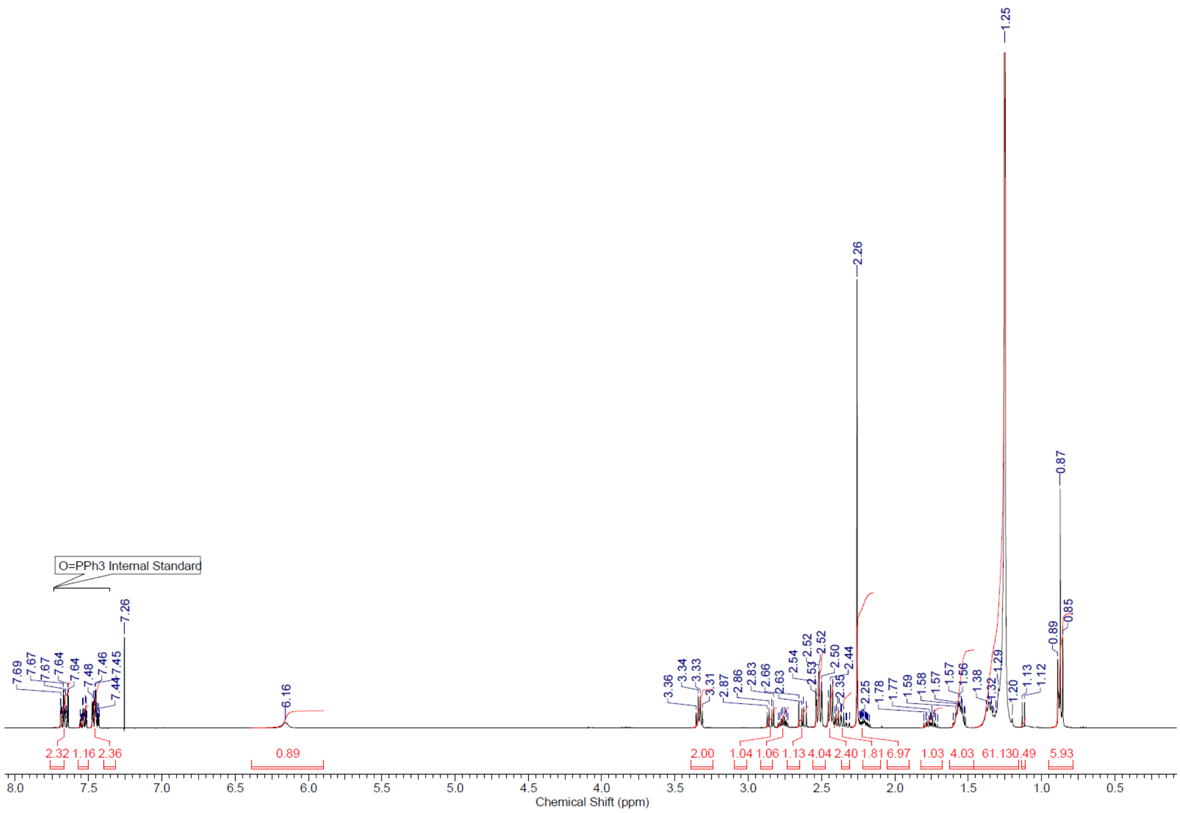


*Figure 50: 1H-NMR of compound A1C18*

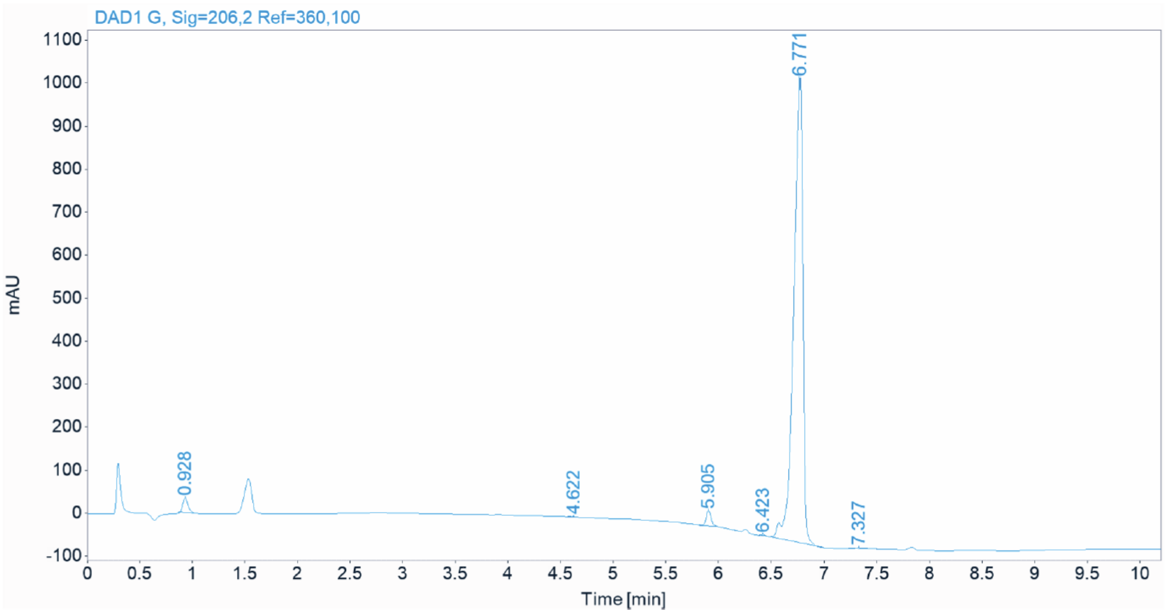

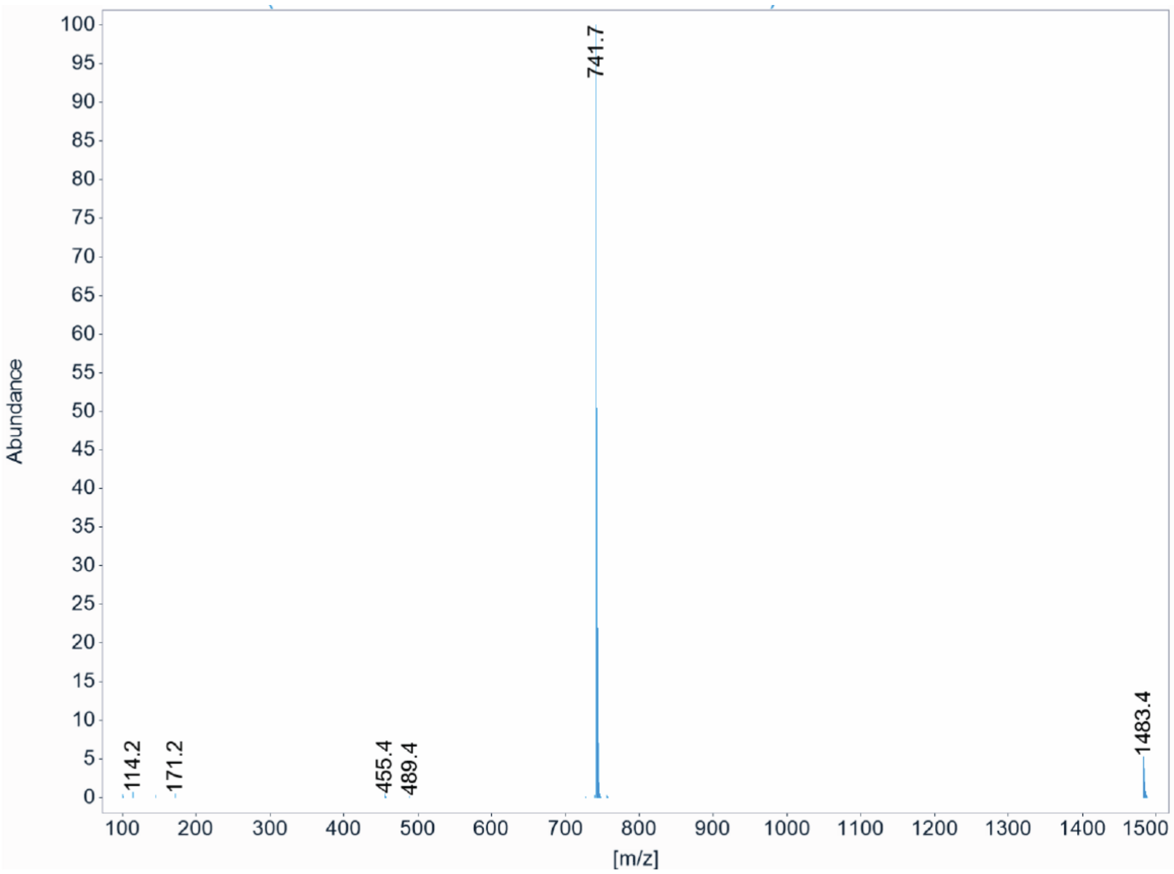


*Figure 51: HPLC-MS of compound A1C18*

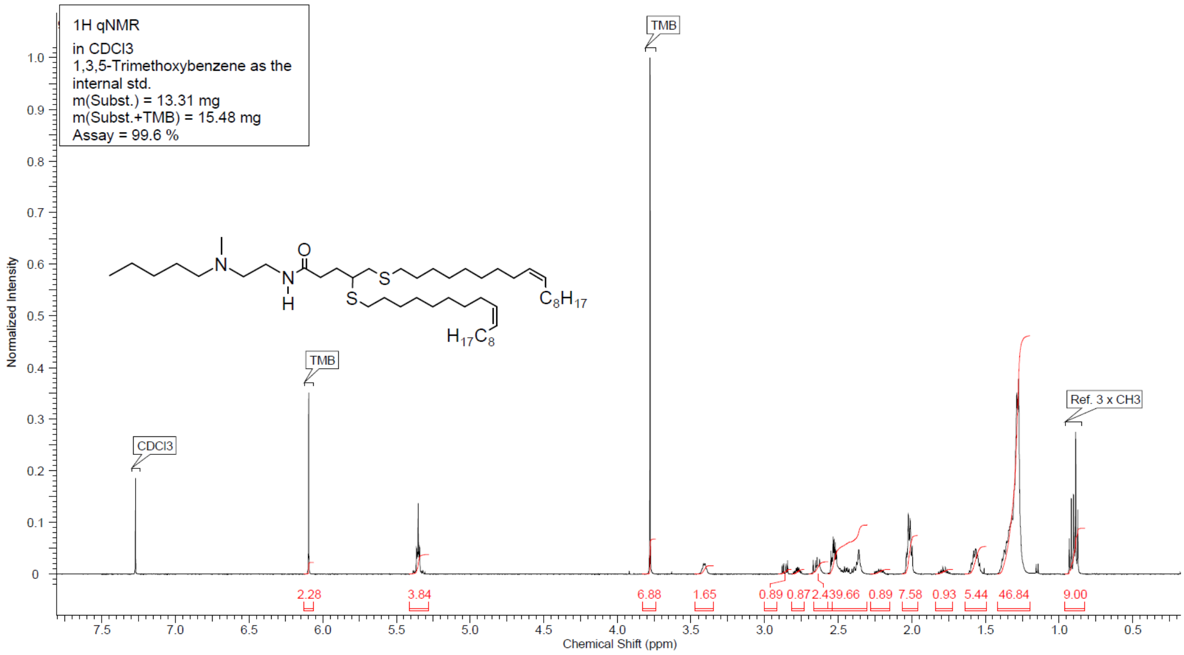


*Figure 52: 1H-NMR of compound A2C18_D9*

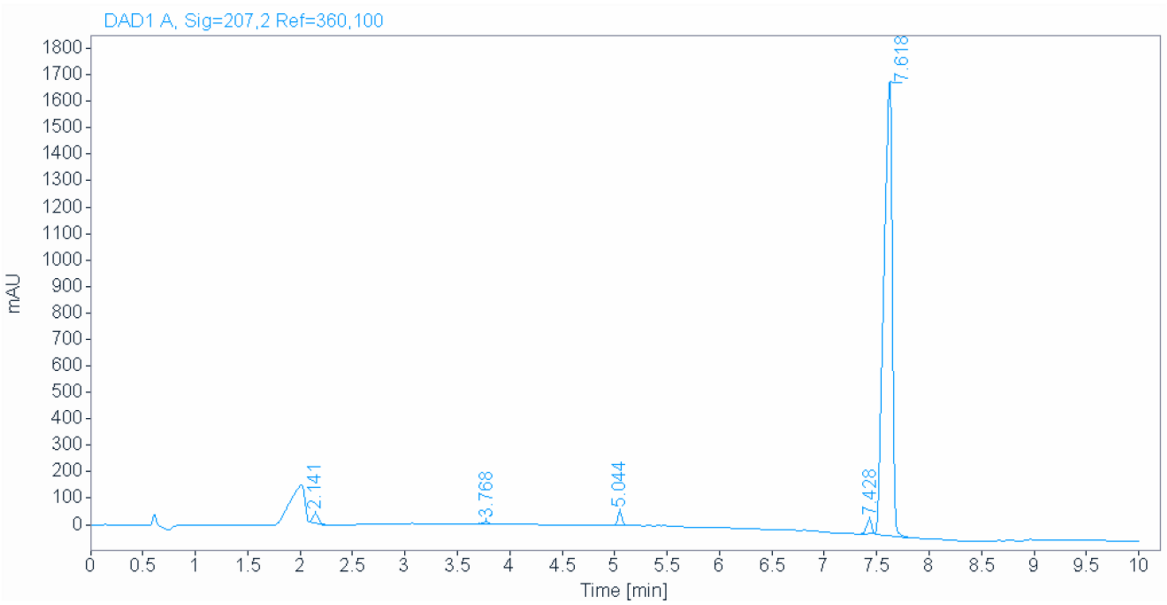


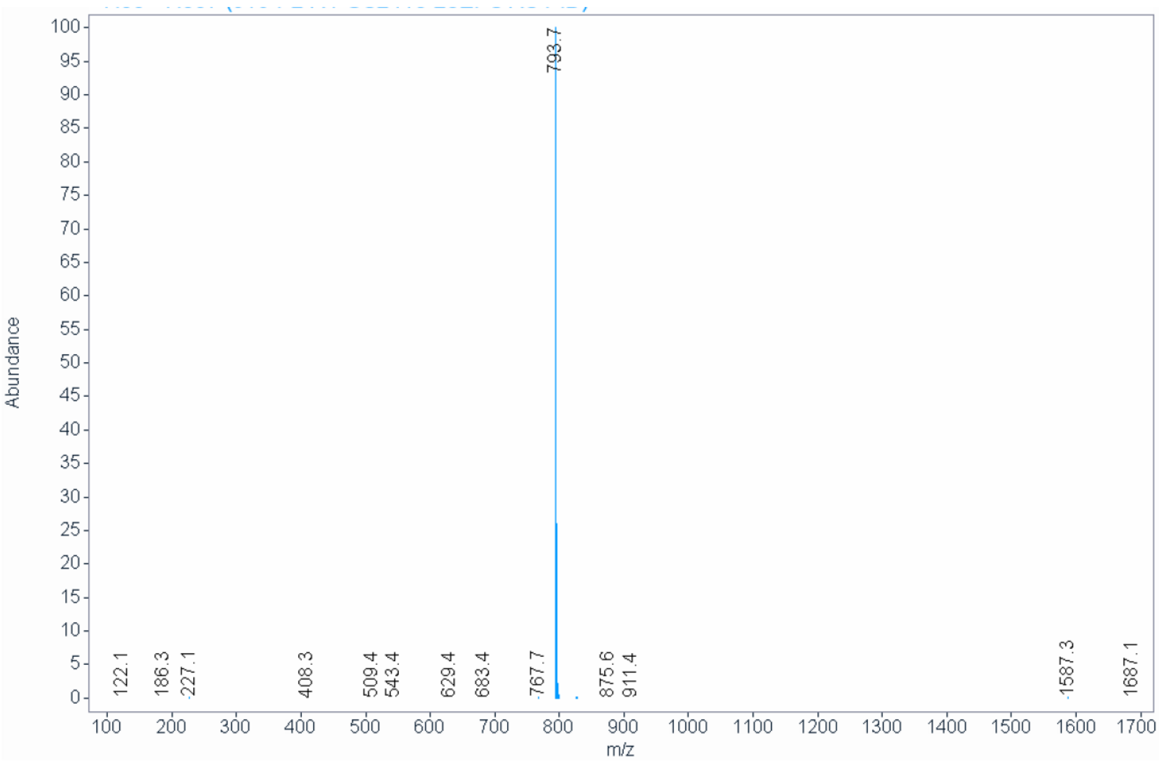


*Figure 53: HPLC-MS of compound A2C18_D9*

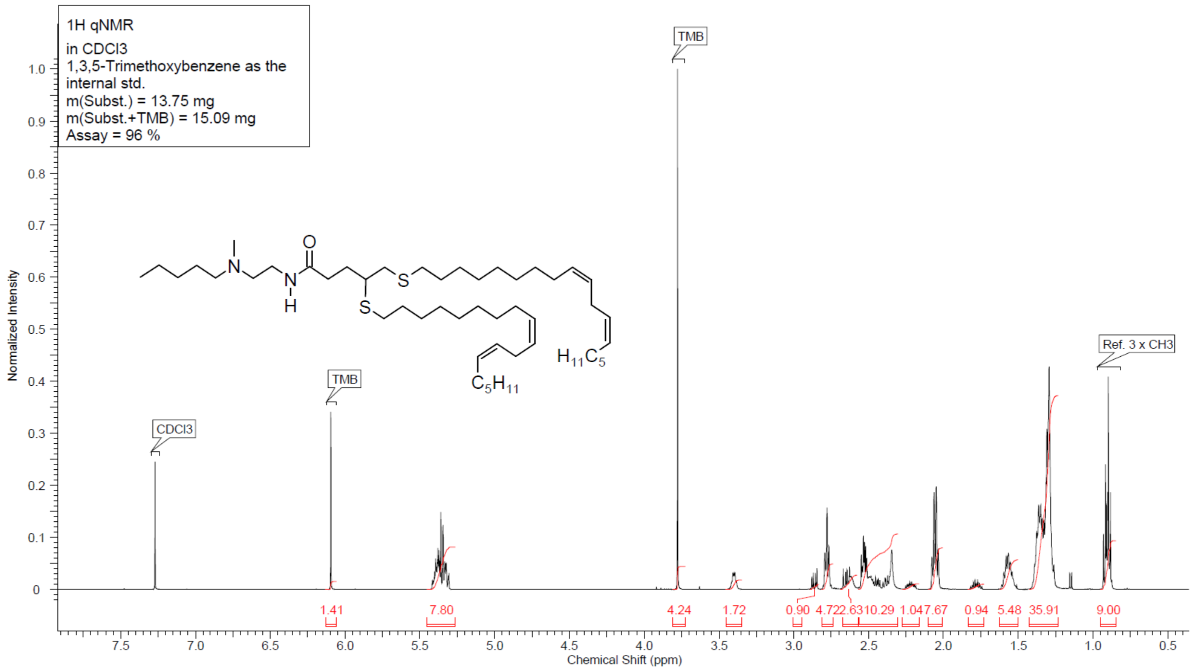


*Figure 54: 1H-NMR of compound A2C18_D9_D12*

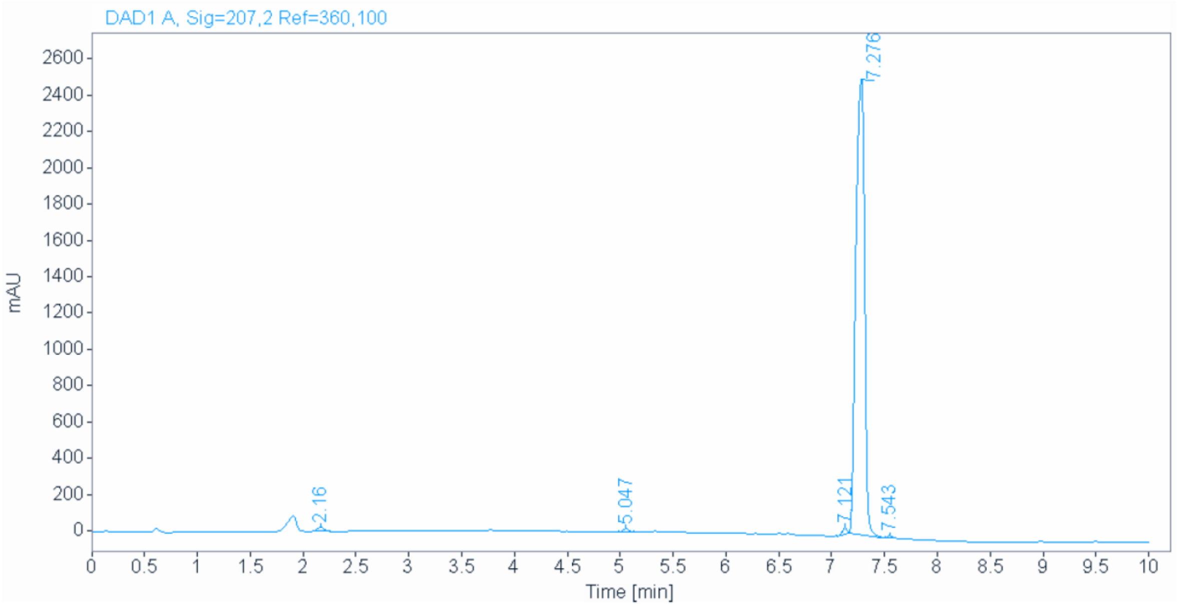

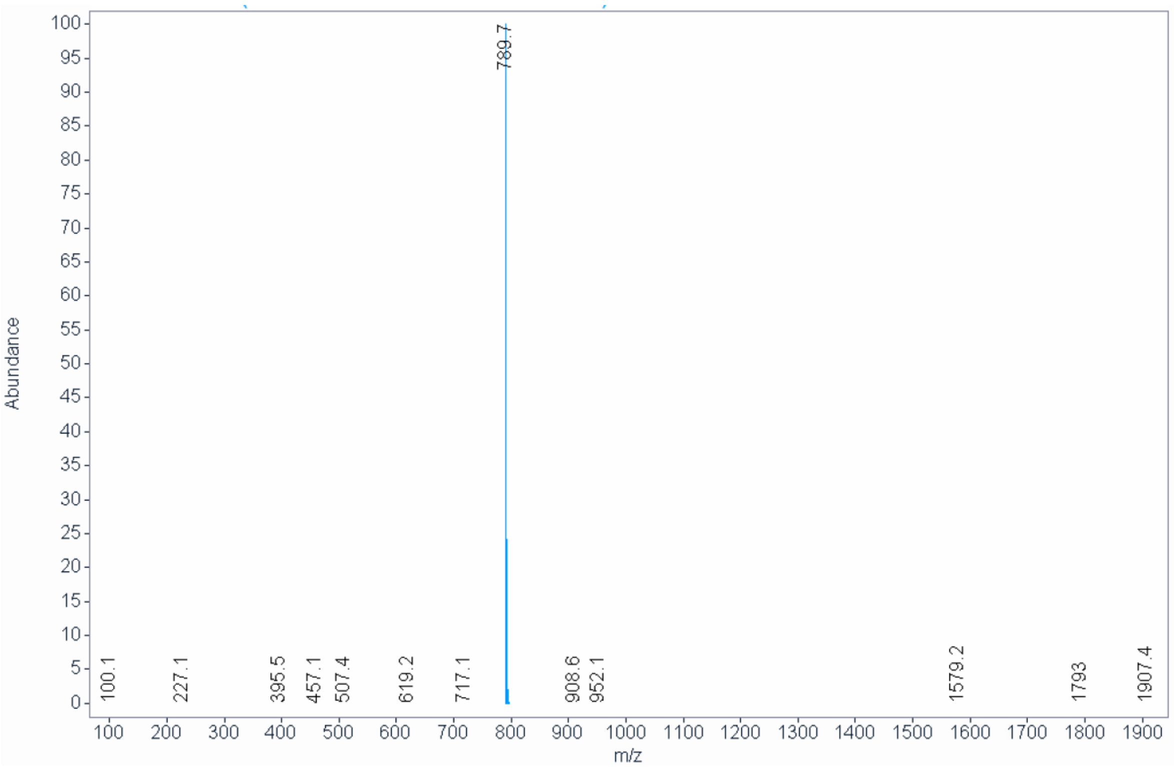


*Figure 55: HPLC-MS of compound A2C18_D9_D12*
